# All *Staphylococcus aureus* bacteraemia strains have the potential to cause infective endocarditis: results of GWAS and experimental animal studies

**DOI:** 10.1101/2022.05.16.491111

**Authors:** Sylvère Bastien, Severien Meyers, Wilmara Salgado-Pabón, Stefano Giulieri, Jean-Phillipe Rasigade, Laurens Liesenborghs, Kyle J. Kinney, Florence Couzon, Patricia Martins-Simoes, Vincent Le Moing, Xavier Duval, Natasha E Holmes, Niels Eske Bruun, Robert Skov, Benjamin P Howden, Vance G. Fowler, Peter Verhamme, Paal Skytt Andersen, Coralie Bouchiat, Karen Moreau, François Vandenesch

## Abstract

**Aims:** Infective endocarditis (IE) complicates 10-20% of *Staphylococcus aureus* bacteraemia (SAB). We aimed to determine whether IE strains of *S. aureus* are genotypically different or behave differently in experimental endocarditis models as compared to non-IE SAB strains.

**Methods and Results:** We conducted a genome wide association study (GWAS) of 924 *S. aureus* genomes from IE (274) and non-IE (650) SAB patients, and tested a subset of strains in two experimental animal models of IE, one studying the early step of bacterial adhesion to inflamed mice valves, the second evaluating the local and systemic developmental process of IE on mechanically damaged rabbit valves. The genetic profile of *S. aureus* IE and non-IE SAB strains did not differ when considering single nucleotide polymorphisms, coding sequences and k-mers analyses in GWASs. In the inflammation-induced IE model in mice no difference was observed between IE and non-IE SAB strains both in adhesion to the cardiac valves and in the propensity to cause IE; in the mechanical IE-induced rabbit model, there was no difference between IE and non-IE SAB strains regarding vegetation size and CFU.

**Conclusion:** *S. aureus* isolates from SAB patients with and without IE were indistinguishable, by GWAS and by two *in vivo* models of IE. Thus, *S. aureus* strain variation is not the primary determinant of IE. Pending the possible identification of host factors predisposing to IE, all strains of *S. aureus* must be considered in patients as capable of causing this common, lethal infection once they have accessed the bloodstream.

**Translational Perspective:** *Staphylococcus aureus* endocarditis (IE) is a deadly complication of *S. aureus* bacteraemia (SAB). Beyond well-identified host related IE risk factors, whether bacterial features may influence the occurrence of IE in the course of bacteraemia remain elusive. We analysed the genomes of 924 *S. aureus* strains from IE and non-IE SAB and compared some in two *in vivo* IE models. We demonstrated that the propensity of *S. aureus* to cause IE in the course of bacteraemia does not depend on the intrinsic genetic or virulence factors of *S. aureus*. These findings are of importance for the management of *S. aureus* bacteraemia.

## Introduction

*Staphylocococcus aureus* bacteremia (SAB) is a severe condition with incidence ranging from 38.2 to 45.7 per 100,000 person-years in the US (1,2). In 10 – 25% of SAB, the infection localises to the valves or endocardial surfaces producing infective endocarditis (IE) (3,4) which is responsible for enhanced morbidity and mortality. To exclude IE upon diagnosis of SAB, use of echocardiography either trans-thoracic or transoesophageal is recommended (5,6) but is far from being systematic: in a recent study analysing 668.423 hospitalizations with *S. aureus* bacteremia from US National Inpatient Sample database (2001-2014) 86.387 (12.9%), only 11% in 2001 to 15% in 2014 had echocardiogram (7). In this context, score-based prediction rules such as the VIRSTA, PREDICT or POSITIVE scores have been proposed to quantify the risk of IE in patient with SAB diagnosis and guide the use of echocardiography (8–10). However, none of these scores take into account the possibility that certain strains of *S. aureus* are more likely to cause IE. Previous studies have compared clinical IE strains with other disease isolates in order to point out which *S. aureus* traits and virulence factors are actually crucial in causing endocarditis. Studies on such bacterial determinants or shared genetic background/lineage by specific multilocus sequence typing (MLST) or clonal complex (CC) yielded contradictory results (11–14). CC12 and CC20 were associated with IE isolates compared with non-IE SAB and skin and soft tissue infection (SSTI) isolates (11). Similarly, CC30 and several adhesins and enterotoxins were associated with IE compared with SSTI isolates and nasal carriage isolates, respectively, but not with non-IE SAB isolates (12,15). The invasive isolates responsible for non-IE SAB were found to be less cytotoxic than non-invasive or colonising isolates, the differences between diseases were supported by polymorphisms in several loci (16). However, when comparing IE with non-IE bloodstream isolates of *S. aureus*, weak (17) or no significant difference between strains was observed (18). Likewise phenotypic traits known or hypothesised to be involved in IE show no significant differences in IE versus non-IE SAB strains (17). More recently, a genome wide association study (GWAS) performed on 241 strains (120 definite IE and 121 non-IE SAB) using a variety of bioinformatics approaches including single nucleotide polymorphisms (SNPs) analysis, accessory genome analysis and k-mers based analysis did not identify clear *S. aureus* genetic markers associated with the occurrence of IE in the course of bacteremia (19). However, the study by Lilje *et al*. had some limitations, including small population size, and limited geographical distribution, emphasising the need to conduct GWAS on a larger sample size.

In order to provide a more robust assessment of the potential role of *S. aureus* factors in the occurrence of IE during bacteraemia, we performed a GWAS analysis of 924 *S. aureus* genomes whether or not *S. aureus* may have caused IE in humans, and completed this genomic approach with two experimental models of IE exploring both early and late events of aortic valve infection.

## Methods

### Bacterial strains

The strain collection was designed to represent the worldwide genetic diversity of *S. aureus* bloodstream isolates isolated from both IE and uncomplicated bacteremia (non-IE SAB). To this end, several patient cohorts were combined (see supplementary material Text S1), achieving the number of 274 definite infective endocarditis according to the modified Duke criteria (20) and 650 uncomplicated bacteraemia (non-IE SAB) patients. All patients underwent either transthoracic or transoesophageal echocardiography and patients with intracardiac devices were excluded. For animal experiments using batch inoculum, a subset of 12 IE and 12 non-IE SAB CC5 strains as well as 10 IE and 9 non-IE SAB CC45 strains were selected within the collection of 924 strains to account for the genomic diversity of the CC5 and CC45 lineages. For single strain experiments, four CC5 (two IE and two non-IE SAB) or eight CC5 (four IE and four non-IE SAB) were arbitrarily chosen for the mouse and rabbit models, respectively.

### Bacterial whole genome sequencing

Libraries were prepared using Nextera® or Truseq® DNA (Illumina, San Diego, CA, United States). Bacterial whole genome sequencing was conducted, depending on centres, on MiSeq® or NextSeq® or HiSEQ-1500® (Illumina, San Diego, CA, United States) platforms with a read length of 2 × 300 bp, 2 × 250 bp or 2 × 150 bp. Further details are in supplementary material Text S1. The extraction and the sequencing of the 241 strains collected during the two prospective studies in Denmark and the US are described in Lilje et al. paper (19).

### Bioinformatic Methods

#### Quality control

First, raw reads were trimmed in order to remove adapters and other Illumina-specific sequences. Then, a sliding window with an average quality threshold trimming of 20 was processed using Trimmomatic v0.36 (21). Read length less than 30 nucleotides were removed. Data were subjected to a quality check through FastQC v.0.11.6 (22) and summarised using MultiQC v1.3 (23).

#### Assembly and Sequence Typing

Assemblies were processed by SPAdes v3.11.1 (24), enabling the automatic coverage cutoff and the mismatch/short indels reducing step. Contigs were broken in k-mers through Kraken v1.1.1 (25) in order to remove contigs which did not share at least one k-mer belonging to *S. aureus*. Finally, contigs less than 300 nucleotides were manually removed. An assembly quality check was visualised through Quast v4.6.1 (26) and visualised by R v3.6.3 (27). Sequence types (STs) were determined by MLST v2.9 (28,29) software and STs were grouped in clonal complexes (CCs).

#### Single nucleotide polymorphisms (SNPs) analysis

Variant calling analysis was performed on the entire data collection using TCH60 (ST30) as the reference genome (NC_017342.1). First of all, this genome was aligned against itself via MUMmer v4.0.0 (30) in order to locate duplicate regions. Then, Snippy v4.4.5 (31) software with a minimum of 10x variant site coverage and a new allele proportion set to 0.5 was used on each data. Result produced a presence/absence matrix for SNP, insertion or deletion positions for the whole genome. Genomic modifications detected in duplicate regions were removed. The remaining genomic changes were encoded as “1” when they were different from the reference. By combining the alignment of core SNPs purged of recombinations in nucleotide sequences by Gubbins v2.4.1 (32) and FastTree v2.1.9 (33), a phylogenetic tree was inferred with a maximum likelihood method and a generalised time reversible model. This global tree was then visualised by iTOL v5.6.1 (34).

#### Gene and non-coding RNA analysis

Annotations were performed by Prokka v1.14.5 (35) with NCTC8325 (NC_007795.1) GenBank annotation file as first protein database. Coding sequences (CDS) identification was reduced to a presence/absence matrix using Roary v3.13.0 (36) with the split paralogs option enabled. A specific database containing 659 genomic non-coding RNA sequences from five strains of *S. aureus* was constructed as follows: 538 ncRNAs from NCTC8325 (http://srd.genouest.org/) were included in the main database. Each ncRNA sequence from other genomes was mapped to the main database using blast v2.2.31 (37) with parameters set to 90% identity and coverage. Finally, 47, 51, 16, and 7 new genomic sequences were added from N315, JKD6008, Newman, and USA300_FPR3757 respectively. This database was then run through Abricate v0.7 (38) with the alignment process set at 90% coverage and identity. The results were transformed into a second presence/absence matrix that was then merged with the one containing the presence/absence of the genes.

#### Genome-wide association study

GWAS was performed using DBGWAS (for *De Bruijn Graph GWAS)* v0.5.2 (39), a method relying on k-mers (all nucleotide substrings of length k found in the genomes) and De Bruijn graphs which connect overlapping k-mers (here DNA fragments), yielding a compact summary of all variations across a set of genomes. Each k-mer presence was calculated both in IE samples and non-IE SAB strain collection. In order to take into account the population structure, the phylogenetic tree produced by variant calling analysis was used. K-mers were set with 3 different values (21, 31 and 41), minor allele frequency filter set to 0.01, and the neighbourhood node number set to 5.

#### Statistics

All statistical analyses were performed by R v3.6.3 (27). Principal component analysis (PCA) and hierarchical clustering (HC) with complete method and Euclidean distance for both rows and columns were performed to screen for possible stratification of the data. Fisher exact tests were used for proportion comparisons, distribution comparisons were screened by Wilcoxon-Mann-Whitney tests and one sample t-tests were used to compare the mean of our data to a known value. Corrections to the p-values of multiple tests were taken into account using a Bonferroni correction method. A threshold of significant p-value was set at 0.05.

A common heritable trait independently from the genetic background was measured by the Pagel lambda phylogenetic value; in this model, the lambda value varies between 0 (absence of phylogenetic signal on trait distributions) to 1 (strong phylogenetic signal) (see Supplementary material Text S1).

The presence or absence of coding sequences and ncRNAs allowing the discrimination of IE strains with non-IE SAB strains was tested by the Least Absolute Shrinkage and Selection Operator (LASSO) regression, accounting for possible bias due to the different geographical origins. LASSO regression is a modification of linear regression where the loss of function is modified to minimise the complexity of the model by limiting the sum of the absolute values of the model coefficients (see Supplementary material Text S1). To examine robustness, Cohen’s Kappa coefficient (a quantitative measure of reliability for two raters evaluating the same object, after adjusting this value for what could be expected from chance alone) was calculated from the best model prediction. The value of Cohen’s Kappa coefficient was compared to the distribution obtained by the 500 predictions of the random assignment of the observed responses while maintaining the link between the phenotype (IE and non-IE SAB) and the geographic origin. Cohen’s Kappa coefficient can range from <0 (no agreement) to 1 (perfect agreement).

Several R packages were used: ade4 v1.7-18 (40); ggplot2 v2.2.1 (41); ComplexHeatmap v2.2.0 (42); factoextra v1.0.7; phytools v0.7-90 (43); permute v0.9-5; glmnet v4.1-2 (44); questionr v0.7.6; effsize v0.8.1 (45); doParallel v1.0-11 (46).

#### Rabbit models

New Zealand white rabbits included both sexes (approximately 60% male) and weighed 2–3 kg. For all experiments, bacterial strains were cultured overnight to stationary phase and washed in PBS before intravenous inoculation. The combined IE/sepsis model was performed as previously described (47) wherein rabbits were injected through the marginal ear veins with either 1–4 × 10^7^ or 4.5–5.2 × 10^8^ CFUs *S. aureus* in sterile saline after damage to the aortic valve with a hard-plastic catheter that was then removed. The doses of *S. aureus* used were previously determined to assure vegetation development by 24 hours with increases in vegetation size occurring over the 4-day test period and to prevent early lethality (death before the 24 h test period). Hence, the following strains were inoculated only at 1–4 × 10^7^ CFU/rabbit: ST20101789 and ST20110560 (IE strains), ST20101791 and ST20120211 (non-IE SAB strains), whilst the four other strains (IE strains: ST20101420, ST20102295; non-IE SAB strains: ST20111372, ST20120206) were inoculated at 4.5–5.2 × 10^8^ CFUs. Animals were monitored 4 times daily during experimentation for development of clinical strokes defined as development of limb paralysis requiring premature euthanasia. Rabbit research was performed with approval given by the University of Iowa Institutional Animal Care and Use Committee. The approved animal protocol number were 1106140 and 4071100.

#### Mice models

To investigate whether IE strains already differ from non-IE SAB strains in initiating the early steps of endocarditis, we examined if these strains adhered differently on inflamed aortic valves by using an early adhesion mouse model as previously described (48). C57BL/6 wild type mice were intravenously injected with 2.10^7^ CFU fluorescent bacteria. Subsequently, a 32-gauge polyurethane catheter (Thermo Fisher, Waltham, USA; 10 lM) was introduced beyond the aortic valve via the right carotid artery and used to inflame the endothelium via an infusion of histamine (200 mM, infusion rate 10 mL/min for 5 min). Afterwards, mice were immediately sacrificed, hearts were harvested and cryosections of the aortic valve were made. Adhesion was visualised by confocal microscopy (LSM880, Carl-Zeiss) and quantified using Imaris (Bitplane, Zurich, Switzerland). To analyse the IE-development, a long-term inflammation-induced endocarditis model was used as previously described (48). In this model, mice were either inoculated with 2.10^6^ CFU *S. aureus* via the tail vein (producing a low proportion of IE) or inoculated with 4.10^6^ CFU *S. aureus* via the transaortic catheter used for histamine infusion (leading to a higher proportion of IE). Mice were followed for three days. The institutional Ethical Committee (KU Leuven) approved all animal experiments (licence number 189/2017).

## Results

### Assemblies and sequence typing

The entire data collection included 274 IE and 650 non-IE SAB from four geographical origins. An assembly summary is reported in Figure S1 and in the Supplementary file. One hundred and six sequence types (STs) were identified among the 924 *S. aureus* strains in the collection. Each isolate was assigned to one of the 28 clonal complexes (Table S1). The data set was essentially composed of six clonal complexes, namely CC30, CC45, CC8, CC5, CC1, and CC15, each accounting for 15.8 to 8.0% of the entire collection and overall representing 74.2% of this collection. The six largest clonal complexes and the five largest sequence type distributions among the geographical origin are visualised in Figure S2.

### Single nucleotide polymorphisms (SNPs) analysis

A total number of 15,436 SNPs were identified among the 924 *S. aureus* samples using the TCH60 (ST30) as reference (Refseq: NC8017342.1) after duplicated regions and recombination corrections. 1127 and 1043 SNPs were respectively present in more than five and ten percent of the entire data collection. Phylogeny of all the 924 strains according to infection type, CCs and STs assignment is shown in Figure 1. Discrimination between IE and non-IE SAB was primary searched through the Pagel lambda phylogenetic signal measurement within the entire strain collection and within each of the four geographical origins (Table S2). The lambda (phylogenetic signal) value of the analysis of the global data set reached 0.9441 and was above the one calculated through random permutations (0.7397 [95% CI: 0.7329, 0.7466]) (t-test, p-value < 2.2e-16). Further analysis showed that the unequal ratio of IE/non-IE SAB between the different cohorts biased the model and that eventually there was no relevant phylogenetic signal explaining the IE and non-IE SAB phenotypes in the whole collection (Supplementary Text S2).

**Figure 1.**
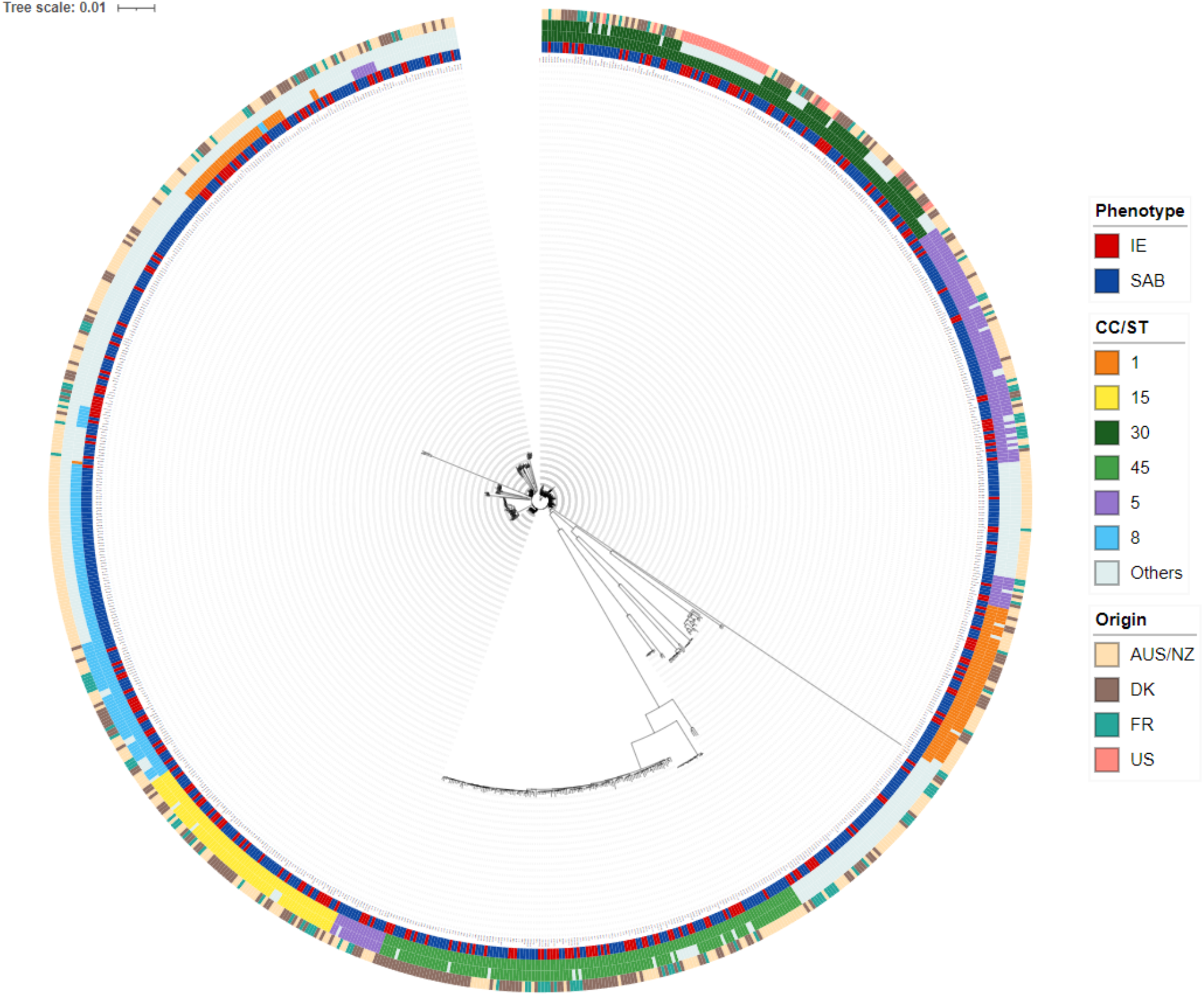
Rooted Phylogenetic tree on all the 15,436 SNPs from the 924 *S. aureus* strains. The tree is rooted by TCH60 *S. aureus* strain (ST30; Refseq: NC8017342.1) using a maximum likelihood method and a generalised time-reversible model after repeated region and recombination corrections. The inner circle represents IE (red) and non-IE SAB (blue) strains. The six largest clonal complexes and five largest sequence types are colored as followed: CC/ST1: orange; CC/ST15: yellow; CC/ST30: dark green; CC/ST45: light green; CC/ST5: purple; CC/ST8: light blue; other CCs or STs are pooled together (grey). The outer strip refers to the four geographical origins of the samples (AUS/NZ: beige; DK: brown; FR: turquoise; US: pink).

### Coding sequences and non-coding RNAs analysis

Beyond single nucleotide polymorphisms, the entire coding sequences and non-coding RNAs were tested in search for a discriminant signal between IE and non-IE SAB strains. Amongst the 20079 different genomic elements that were annotated by Prokka (35) and blasted (37) through the 924 samples, 1002 genes and/or ncRNAs were discarded since they were shared by all samples in the entire strain collection. 13692 genes were found in less than 5% of the strains and were considered as accessory genes. Finally, 4757 (∼23%) genes and/or ncRNAs were found in more than 5% and less than 95% of the samples and were used for analyses.

Hierarchical clustering (Figure 2) and PCA plots (Fig. S4) on the 4757 coding sequences and ncRNAs revealed a major contribution of clonal complex membership to the presence/absence of coding sequences and ncRNAs. Conversely, samples were not distinguished according to their clinical phenotype as it can be seen in the PCA plots (Fig. S4).

**Figure 2.**
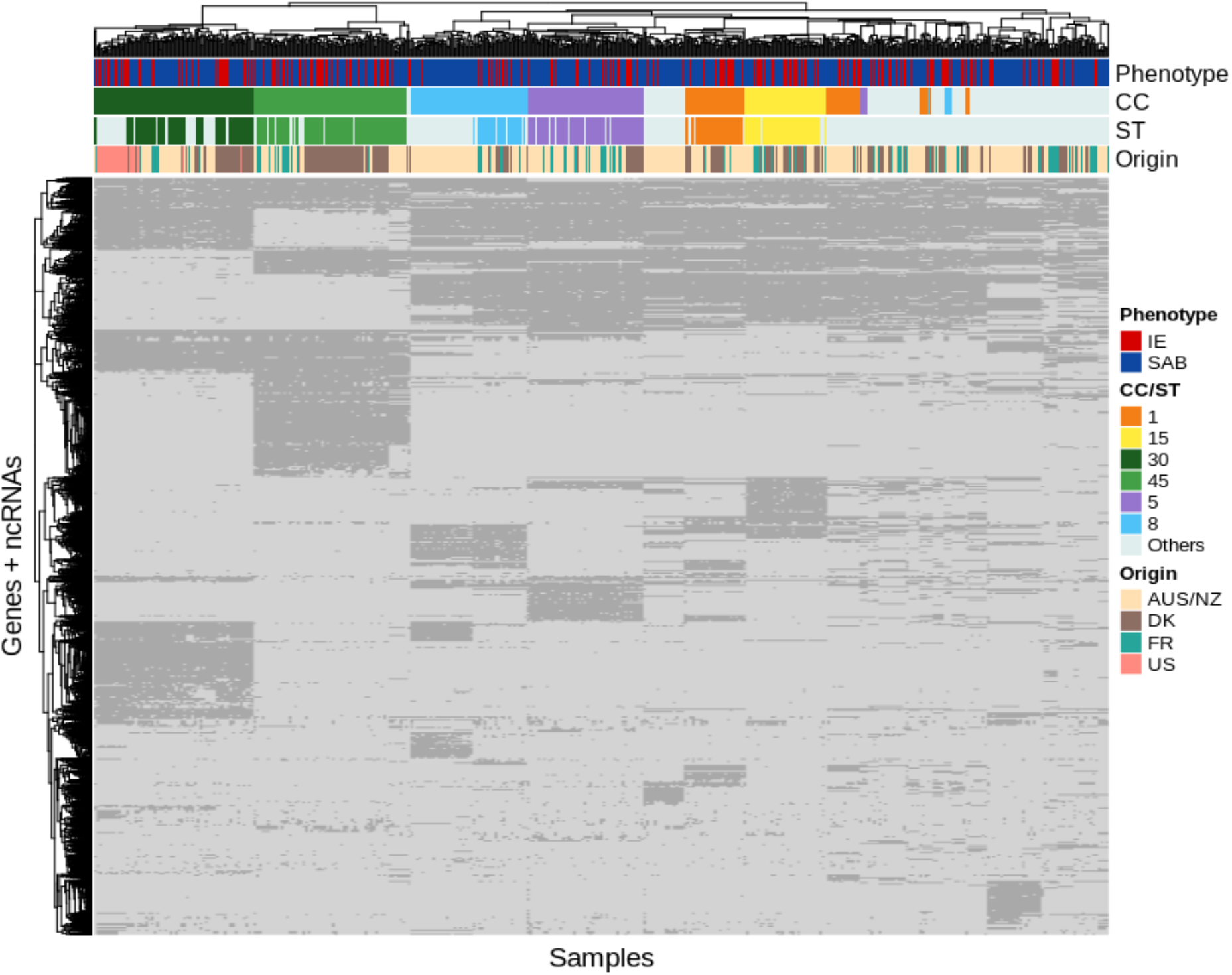
Hierarchical clustering on the 4757 coding sequences and non-coding RNAs presence (dark grey), and absence (light grey) on the 924 *S. aureus* strains. The blue/red stripe on top shows the infection type, either IE (red) or non-IE SAB (blue). The 2 strips below correspond to the six major clonal complexes and the five major sequence types respectively (CC/ST1: orange; CC/ST15: yellow; CC/ST30: dark green; CC/ST45: light green; CC/ST5: purple; CC/ST8: light blue; Others: grey). The last strip refers to the four geographical origins of the samples (AUS/NZ: beige; DK: brown; FR: turquoise; US: pink).

The presence or absence of genes and ncRNAs allowing the discrimination of IE strains with non-IE SAB strains was tested by LASSO regression taking into account the various geographical origins (see Methods). Our entire procedure which was computed 1000 times with novel subsampling for training and validation sets reached a mean of -0.2274 [-0.4312, 0.0070] (of note Cohen’s Kappa coefficient can range from <0 (no agreement) to 1 (perfect agreement), see Methods) (Fig. S5). Although some of these Kappa values (before permutation) are above the Kappa distribution (i.e., Kappa value exceeding the mean of the null distribution) obtained through random permutation (red dots), these values remain far from significance (=1). Therefore, it is not possible to discriminate between IE and non-IE SAB strains based on presence/absence of genes and non-coding RNAs.

### K-mer analysis

IE and non-IE SAB assemblies were split in k-mers and tagged in order to search specific genomic signatures associated with IE through DBGWAS (39) including a correction for population structure. Very few K-mers reached significance and their relevance was ruled out by further analysis (Supplementary Text S2).

### Mice model of infective endocarditis

This model was chosen to test the initial adhesion of *S. aureus* to inflamed aortic valves in mice (48). Such a model is particularly relevant in this context where we hypothesized that the difference between IE and non-IE SAB strains take place at the early steps of the process when blood-circulating bacteria would seed and develop on cardiac valves. In a screening -broad-approach we have chosen to pool 12 CC5 IE strains in a single batch and 12 CC5 non-IE SAB strains in another one (Supplementary excel File). CC5 was chosen because it represented the most frequent lineage in the French VIRSTA collection where IE and non-IE SAB isolates were matched on age and sex in the same geographical setting (3). In a first experiment, we tested whether bacteraemia and endocarditis strains adhere differently to histamine-inflamed aortic valves. Results revealed that the endocarditis and bacteraemia pools adhere equally on inflamed valves (p = 0.5619) (Fig. 3).

**Figure 3.**
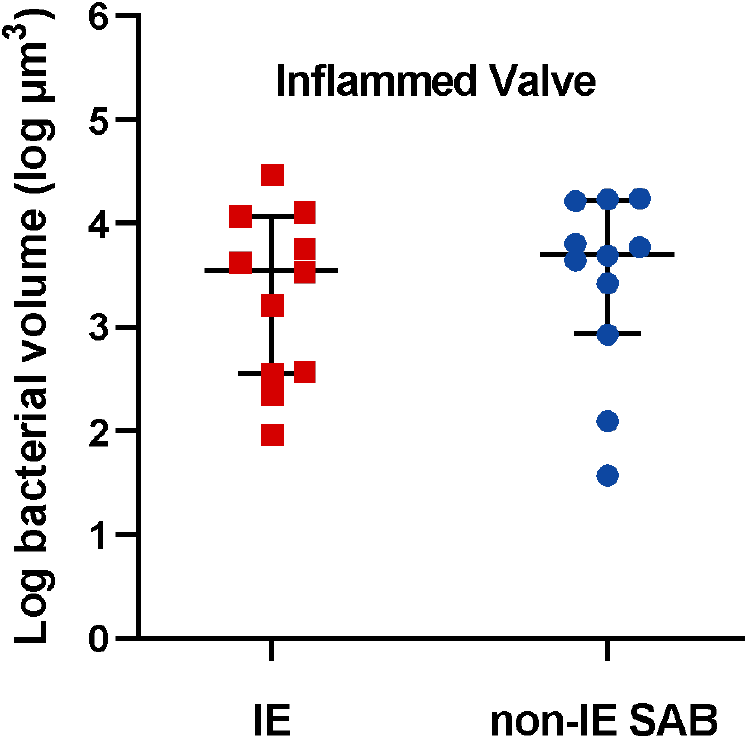
Adhesion of endocarditis and bacteraemia CC5 strain pools on inflamed cardiac valves. *Staphylococcus aureus* CC5 strain pool (2.10^7^ CFU/mouse) were intravenously injected in C57BL/6 mice. Subsequently, via a transaortic catheter, valves were inflamed (5 min of histamine infusion). Mice were immediately sacrificed and adhesion was quantified. Adhesion on inflamed valves of a *S. aureus* CC5 IE strain pool (12 CC5 IE strains, number of mice = 11) was compared with a *S. aureus* CC5 non-IE SAB strain pool (12 CC5 non-IE SAB strains, number of mice = 11, p-value = 0.5619). Results represent log-transformed vegetation volumes in single mice. Median values ± interquartile range are represented; *P < 0.05; Mann-Whitney Wilcoxon test.

We then tested whether the CC5 IE strain pool can cause more IE histologic lesions at Day 3 than the CC5 non-IE SAB strain pool. We observed that both pools equally caused IE on inflamed valves (CC5 OR [95 % CI]: 5.01 [0.85 – 54.43], p-value = 0.0738, Fig. 4). The challenge was repeated on four CC5 isolates distributed along the phylogeny of the CC5 collection (Supplementary excel File) using the inflammatory model as above and a more severe variation of this model obtained by injecting the inoculum via the transaortic catheter. Results revealed no significant differences between the IE and non-IE SAB strains (HS OR [95 % CI]: 2.14 [0.55 – 8.89], LS OR [95 % CI]: Inf. [0.02 – Inf.], Fig. 5).

**Figure 4.**
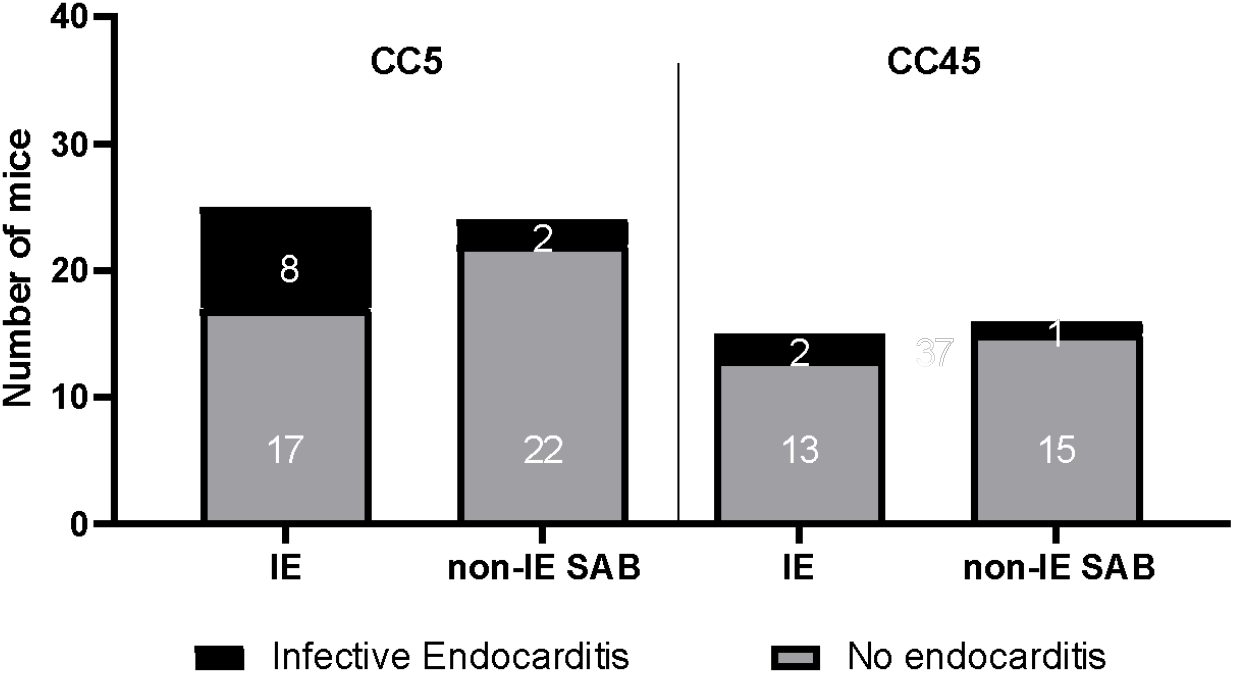
Propensity of endocarditis and bacteraemia CC5 and CC45 strain pools to cause IE on inflamed cardiac valves. *Staphylococcus aureus* CC5 or CC45 strain pools (2.10^6^ CFU/mouse) were intravenously injected in C57BL/6 mice. Subsequently, via a transaortic catheter, valves were inflamed (5 min of histamine infusion). Afterwards, the catheter was removed, mice were monitored for three days to see if endocarditis developed. Proportion of mice that developed endocarditis (i) in the CC5 strain pool vs. no endocarditis (IE strain pool: 12 IE strains, number of mice = 25; non-IE SAB strain pool: 12 non-IE SAB strains, number of mice = 24; OR [95 % CI]: 5.01 [0.85 – 54.43], p-value = 0.0738). (ii) in the CC45 strain pool vs. no endocarditis (IE strain pool: 10 IE strains, number of mice = 15; non-IE SAB strain pool: 9 non-IE SAB strains, number of mice = 16; OR [95 % CI]: 2.25 [0.11 – 144.93], p-value = 0.5996).

**Figure 5.**
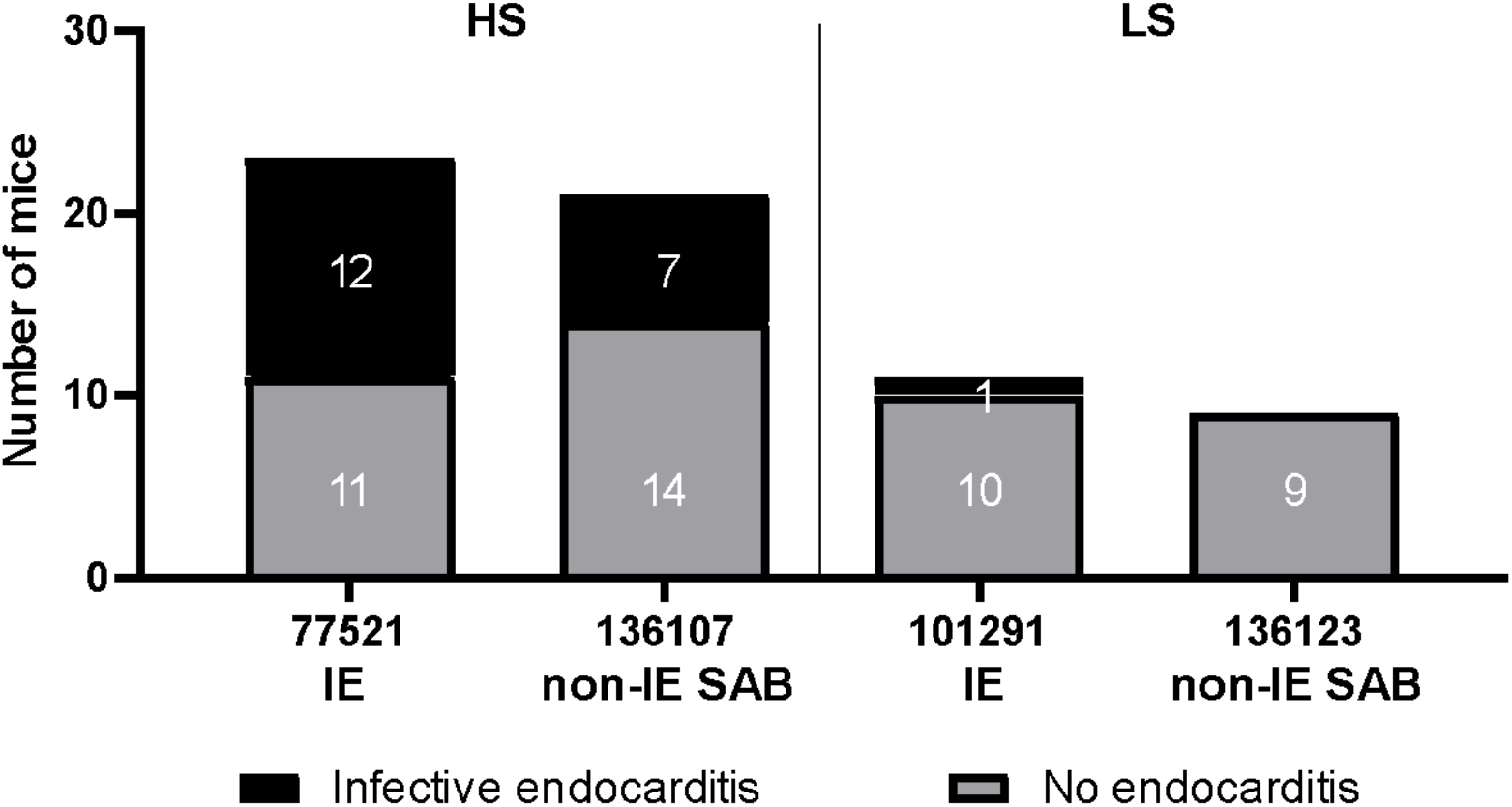
Propensity of endocarditis and bacteraemia single strains of CC5 to cause IE on inflamed cardiac valves. *S. aureus* IE and non-IE SAB strains (2-4.10^6^ CFU/mouse) were perfused via the tail vein catheter (strain 101291 and 136123) (low severity model, LS) or via a transaortic catheter (increasing the model severity, HS) (strain 77521 and 136107) in C57BL/6 mice. Valves were inflamed by histamine infusion (5 min) via a transaortic catheter. Afterwards, the catheter was removed and mice were monitored for one (strain 77521 and 136107) or three days (strain 101291 and 136123) to see if endocarditis developed. Proportions of endocarditis in mice infected with an IE strain (strain 77521, number of mice = 23) compared with a non-IE SAB strain (strain 136107, number of mice = 21) (HS OR [95 % CI]: 2.14 [0.55 – 8.89], p = 0.2387) and with an IE strain (101291, number of mice = 11) compared with a non-IE SAB strain (strain 136123, number of mice = 9) (LS OR [95 % CI]: Inf. [0.02 – Inf.], p > 0.9999). Fisher’s exact tests (P*<0.05).

The lack of robust evidence with CC5 lineage drove us to test CC45, another common CC in the VIRSTA cohort study. A pool of 10 IE isolates was tested in comparison to a pool of 9 non-IE SAB isolates in the inflammatory model. Both pools gave equal amounts of endocarditis (CC45 OR [95 % CI]: 2.25 [0.11 – 144.93], p = 0.5996, Fig. 4).

### Rabbit model of infective endocarditis

The above results suggested that the strain potential to initiate IE was similar between IE and non-IE SAB strains. Having established that IE and non-IE strains have similar ability to initiate IE in the setting of SAB, we sought to investigate whether there were strain-level differences in the developmental process of IE, such as the size of the vegetation or the blood bacterial load. To this end we used the mechanical IE-induced rabbit model, a robust experimental model in the field of IE pathogenesis (47). To optimise the signal/noise ratio, strains were selected within a single CC (CC5 which is one of the most common in the global collection). Four strains of IE and four strains of non-IE SAB were selected for the challenge in rabbits (Supplementary excel File). Bacteria were injected intravenously from 1 × 10^7^ – 5 × 10^8^ CFUs/rabbit, after mechanical damage of the aortic valves, and infection allowed to progress for a maximum of 4 days. The doses of *S. aureus* used were previously determined to ensure vegetation development by 24 hours with increases in vegetation size occurring over the 4-day test period and to prevent early lethality. Upon sacrifice at day 4, all rabbits presented with aortic vegetations with a broad variation with mean values across strains between 15-93 mg for vegetation size, 1.7 × 10^7^ - 2.6 × 10^9^ for vegetation CFU (most where in the 10^8^ range), and 1.3 × 10^3^ - 2.1 × 10^5^ for blood CFU (Figure 6A-C). Kruskal-Wallis tests were not statistically significant suggesting that none of the strains increase vegetation weight (p = 0.0699, IE vs. non-IE SAB effect size = -0.0277 [-0.6109, 0.5554]) and the number of viable bacteria either in vegetation (p = 0.1822, IE vs. non-IE SAB effect size = 0.0222 [-0.4413, 0.4857]) or in blood (p = 0.7356, IE vs. non-IE SAB effect size = 0.2145 [-0.3702, 0.7993]).

**Figure 6.**
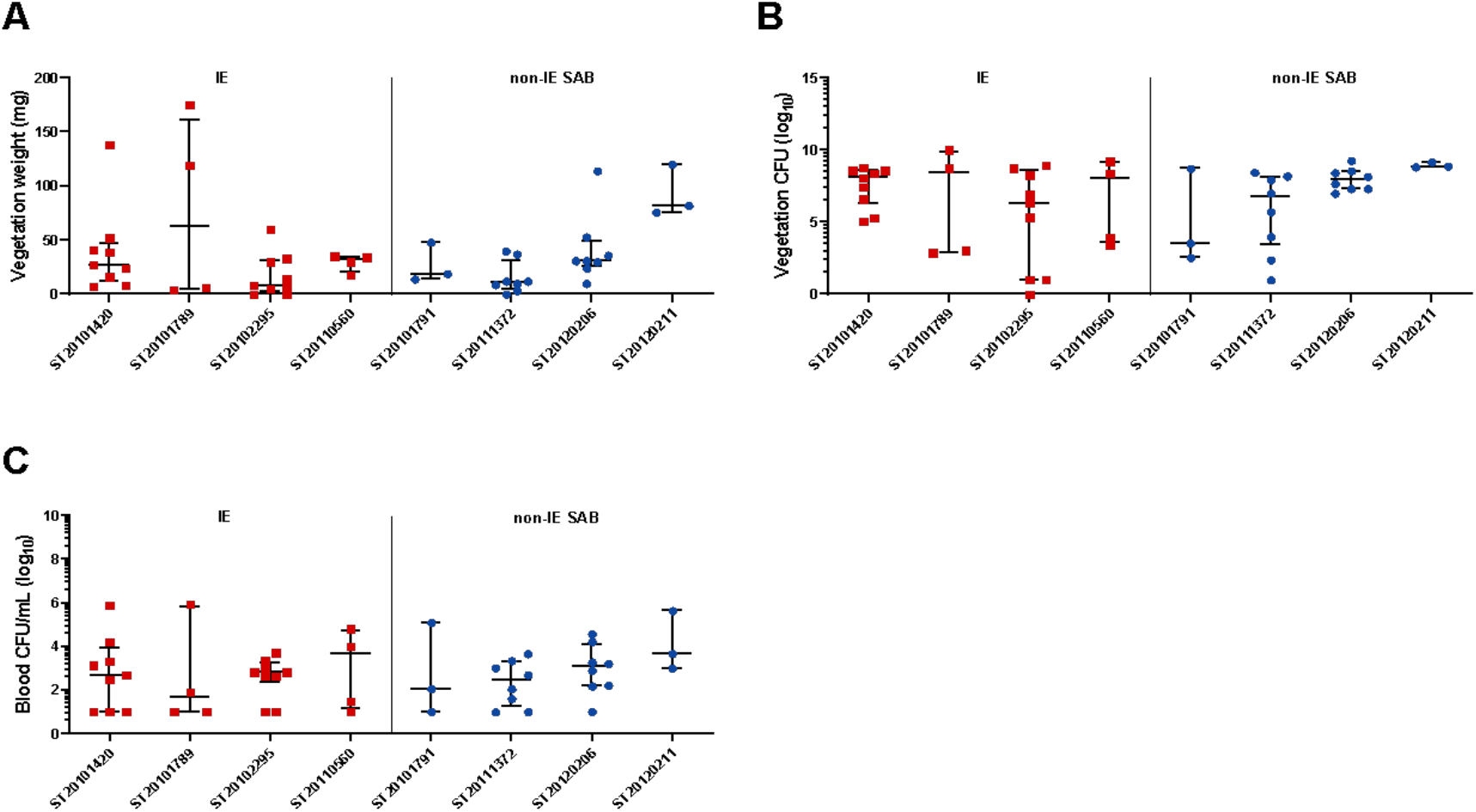
Native valve, infective endocarditis in rabbits with individual bacteraemia and infective endocarditis strains from the CC5 clonal group. Rabbits were infected intravenously with 1 × 10^7^ to 5 × 10^8^ CFU after mechanical damage of the aortic valves. (A) Total weight of vegetations dissected from aortic valves; (B) bacterial counts recovered from aortic valve vegetations shown in panel A; (C) Bacterial counts per millilitre of blood at end of experimentation. Horizontal lines and error bars represent median values with interquartile ranges. None of the strains are statistically different from each other. Kruskal-Wallis tests (P*<0.05; A, p=0.0699; B, p=0.1822; C, p=0.7356).

Altogether, the model of damage-induced infective endocarditis in rabbits showed that all strains, irrespective of their clinical origin in humans (IE or non-IE) induced IE of undistinguishable severity.

## Discussion

We herein aimed to address the question of whether specific *S. aureus* strains cause IE in the course of bacteraemia. To ensure a comprehensive answer, we based our approach on (i) a GWAS analysis and (ii) two different animal models exploring both early and late events of aortic valve infections, and both mechanical and inflammation damage induced IE. All these approaches concordantly reveal that IE isolates could not be discriminated from non-IE SAB isolates based on phylogeny, gene and non-coding RNA content, or i*n vivo* virulence.

We analysed 924 *S. aureus* strains of Duke-definite native valve IE and IE excluded-bacteraemia originating from France, Denmark, Australia/New-Zealand and the US, which is, to our knowledge the largest collection ever described of such well-defined isolates. The bacteraemia group contained only cases where the patient underwent transthoracic and/or transoesophageal echography, and the endocarditis group included only definite endocarditis on native valves according to the modified Duke criteria. Since we utilised different cohorts from various countries, there was a risk of a geographic bias. However, this potential bias was mitigated either by analysing each cohort independently or by including the geographic origin as a variable. Another result supporting this assertion is the fact that the genetic diversity of the global collection based on clonal complexes (CC) is in line with other studies with CC45, CC30, CC5, CC8, CC15, CC1 being the most prevalent (11,12,15,18).

Concerning genetic analyses, bioinformatics issues of every specific method were overcome using different strategies from gene and non-coding RNA presence/absence, single nucleotide polymorphisms and k-mers approaches. With the assumption that it is unlikely to find an association between IE with a single genetic event, a combination of markers was screened through the Pagel lambda and LASSO algorithm. We therefore, did not confirm the prevalence of specific virulence factors genes in IE strains as previously described by others (11,12,17). In a previous study, using microarray data from a small cohort (72 IE and 54 SAB) and discriminant analysis of principal component (DAPC) multivariate analysis method, we were able to discriminate IE from non-IE SAB and reassigned IE strains in 80.6% of the cases (17). The present study performed on a much larger cohort and relying on whole genome sequence data did not confirm our previous observations, which was likely biased by its small sample size and its unbalanced repartition of IE vs non-IE SAB (see Supplementary text S2 for the Pagel Lambda analysis).

Although based on over 900 genomes, the number of genetic markers tested is higher than the number of strains, which is a statistical limitation of our study. This limitation was taken into account in particular in the use of the LASSO algorithm, which eliminates non-essential variables from the final model.

If factors associated with the occurrence of IE do exist, we can conclude that there is no common heritable trait independently from the genetic background; rather, several independent pathways could be at play as an equivalent of “phenotypic convergence”, the phenotype being the propensity to cause IE. Other studies using GWAS to explore the transition from nasal carriage of *S. aureus* to SAB have not identified genomic predictors of bacteraemia versus carriage, perhaps due to the possible polygenic nature of this transition (49,50). Denamur *et al*. estimated that increasing sample size by an order of magnitude may provide enough power to detect subtle differences in a subset of strains (51). However, inclusion of such a number of patients will be challenging, and if genetic markers are identified, they are likely to be valid only for that one lineage (i.e., a specific ST or CC), and thus difficult to exploit in a diagnostic context. Some concerns in genome-wide association studies are the use of only one reference genome and therefore the risk of missing SNPs on genes not present in this genome. To overcome this bias, we performed k-mers analyses that did not require the use of a reference strain and which accounted for lineage effects in the linear mixed model by including the phylogenetic distance calculated from the variant calling analysis.

Previous *in vitro* experiments exploring numerous functions reported or hypothesised to be involved in staphylococcal IE pathogenesis (resistance to microbicidal peptides, adherence to fibrinogen and fibronectin, biofilm formation, staphylokinase production, platelet aggregation, CD69 superantigen-induced expression and adhesion to and internalisation by endothelial cells) failed to discriminate IE from non-IE SAB strains (17). To ensure an ultimate “functional” view of the issue, we explored IE animal models, using both murine and rabbit models, as well as both prior mechanical and inflammatory valve damage. By using the murine model developed by Liesenborghs *et al*. (48), that explores the initial stages of *S. aureus* IE, we showed that all tested *S. aureus* strains could adhere to inflamed valves and induce IE. Using this unique model, no significant difference was observed, whether using CC5 or CC45 strains. A weak signal was obtained when testing the CC5 strains in batches but was not confirmed when testing individual strains. We then used the damage-induced IE model in rabbits that showed that all strains, irrespective of their clinical origin in humans (IE or non-IE) induced IE of undistinguishable severity. Moreover, the apparent between-strains variations in vegetation weight, vegetation CFU and blood CFU never reached significance suggesting that all strains were equivalent regarding severity of IE.

A growing body of evidence suggests that human genetic variability may influence the risk for and severity of *S. aureus* infections, including complicated SAB and IE. Evidence supporting a genetic basis for susceptibility to SAB includes: i) higher rates of *S. aureus* infections in distinct ethnic populations (52–56); ii) familial clusters of *S. aureus* infection (57–60); iii) rare genetic conditions causing susceptibility to *S. aureus* (61–66); and iv) variable susceptibility to *S. aureus* in inbred mice (67–71) and cattle (72,73). The first GWAS evidence of human genetic susceptibility to *S. aureus* infection by comparing 4,701 cases of *S. aureus* infection and 45,344 matched controls was presented (74). These results were reinforced in an admixture mapping study identifying the HLA class II region on chromosome 6 associated with SAB susceptibility (75). Most recently investigators demonstrated a genetic basis for both complicated SAB (76) and persistent MRSA bacteremia (77). For example, Mba Medie employed a carefully matched group of patients with both persistent and resolving MRSA bacteraemia. They found that a single polymorphism in DNMT3A was significantly associated with: i) resolving MRSA bacteremia; ii) increased global methylation; and iii) reduced levels of IL-10 (77). In 2018, we published the first GWAS on 67 patients presenting definite native valve IE (cases) compared to 72 patients with SAB and identified 4 SNPs located on chromosome 3 associated with SaIE (78). However, this result did not reach conventional genome-wide significance, possibly due to the small size of the cohorts. This highlights the need to analyse large collections of samples, which can only be obtained as part of an international multicenter project.

Another explanation for the lack of association between bacterial or human genetic factors and IE might be that the “IE phenotype” is related to differential (bacterial or human) gene expression or epigenetic changes that cannot be captured through whole-genome sequencing.

One limitation of this study is the heterogeneous nature of the various cohorts, notably the ratio of IE to non-IE SAB, although all cases were either Duke-definite native valve IE or IE excluded-bacteraemia, which mitigates this bias. A second limitation is the interpretation of a negative result when considering biological exploration and even animal models. One cannot exclude that future study involving a novel animal experimental model or exploring some novel biological feature of bacterial strains may reveal significant discriminating features of IE versus non-IE SAB strains. As of today, the present study which compared the largest collection of *S. aureus* bacteraemia and endocarditis strains, indicates that these strains cannot be differentiated on the basis of genomic data or experimental animal models; rather, the conclusion is that all *S. aureus* isolated from blood culture could potentially cause infective endocarditis. In the management of *S. aureus* bacteraemia, where the place of ultrasound investigation, in particular transoesophageal ultrasound, is debated, our results argue for a more systematic investigation of SAB.

## Supporting information

Summary-genomes-assemblies

## Acknowledgments

We thank Bernard Iung at the Hopital Bichat (France) for critical reading.

## Data availability statement

The data underlying this article are available in several different repositories detailed in Supplementary Text S2

## Funding

This study did not receive dedicated funding. VGF was supported in part by grant 1R01AI165671 from the National Institutes of Health. SB was supported in part by the French National Research Agency (ANR IDAREV).

## Disclosures

FV reports research funding outside the scope of the present study by bioMérieux, two patents pending in antimicrobial resistance detection and shares in Weezion. VGF reports personal fees from Novartis, Novadigm, Durata, Debiopharm, Genentech, Achaogen, Affinium, Medicines Co., Cerexa, Tetraphase, Trius, MedImmune, Bayer, Theravance, Basilea, Affinergy, Janssen, xBiotech, Contrafect, Regeneron, Basilea, Destiny, Amphliphi Biosciences. Integrated Biotherapeutics; C3J, grants from NIH, MedImmune, Cerexa/Forest/Actavis/Allergan, Pfizer, Advanced Liquid Logics, Theravance, Novartis, Cubist/Merck; Medical Biosurfaces; Locus; Affinergy; Contrafect; Karius; Genentech, Regeneron, Basilea, Janssen, from Green Cross, Cubist, Cerexa, Durata, Theravance; Debiopharm, Royalties from UpToDate; and a patent pending in sepsis diagnostics.

## Notes

### Competing Interest Statement

FV reports research funding outside the scope of the present study by bioMerieux, two patents pending in antimicrobial resistance detection and shares in Weezion. VGF reports personal fees from Novartis, Novadigm, Durata, Debiopharm, Genentech, Achaogen, Affinium, Medicines Co., Cerexa, Tetraphase, Trius, MedImmune, Bayer, Theravance, Basilea, Affinergy, Janssen, xBiotech, Contrafect, Regeneron, Basilea, Destiny, Amphliphi Biosciences. Integrated Biotherapeutics; C3J, grants from NIH, MedImmune, Cerexa/Forest/Actavis/Allergan, Pfizer, Advanced Liquid Logics, Theravance, Novartis, Cubist/Merck; Medical Biosurfaces; Locus; Affinergy; Contrafect; Karius; Genentech, Regeneron, Basilea, Janssen, from Green Cross, Cubist, Cerexa, Durata, Theravance; Debiopharm, Royalties from UpToDate; and a patent pending in sepsis diagnostics.

