## Supplementary material for "All *Staphylococcus aureus* bacteraemia strains have the potential to cause infective endocarditis: results of GWAS and experimental animal studies"

#### Text S1. Supplementary Material and Methods

**Patient cohorts.** The final collection comprised 924 strains distributed as follows: 130 strains (72 IE and 58 non-IE SAB cases) from the French national prospective multicentre cohort VIRSTA (1); 26 strains (13 IE and 13 non-IE SAB) from the Danish National *Staphylococcus aureus* Bacteraemia Repository (Statens Serum Institute), 40 strains (20 IE and 20 non-IE SAB) obtained from the *Staphylococcus aureus* Bacteraemia Group (SABG) biorepository at Duke University (2), 487 strains (49 IE and 438 non-IE SAB) from two Australasian cohorts of SAB, namely the vancomycin sub study of the Australian and New Zealand Cooperative on Outcome of Staphylococcal Sepsis (ANZCOSS) (3) and the Vancomycin Efficacy in Staphylococcal Sepsis in Australasia (VANESSA) study (4); and 241 strains (120 IE and 121 non-IE SAB) collected during two prospective studies in the Denmark and the US (5,6) and already analysed by GWAS (7).

**Bacterial whole genome sequencing**. For the IE and non-IE SAB collected from the two Australasian cohorts of SAB, extraction of genomic DNA from single colonies was performed using the Janus® automated workstation (PerkinElmer) or manually using Invitrogen PureLink genomic DNA kit or the Sigma GenElute kit. Library was prepared using Nextera® XT DNA (Illumina, San Diego, CA, United States), DNA concentration was normalised to 0.2 ng/µl. For the strains collected from the French cohort, extraction of genomic DNA from single colonies was performed using either using Qiagen DNeasy Blood and Tissue Kit (Qiagen, Hilden, Germany) or the Roche Magnapure 96 automated workstation (Roche) both with a customised pre-treatment step with lysozyme (20 mg/ml) and lysostaphin (100 µg/mL) for a 30 min incubation period at 37 C or using the UltraClean® Microbial DNA Isolation Kit (MO BIO, Carlsbad, CA, United States).

Libraries were prepared using Nextera® XT DNA (Illumina, San Diego, CA, United States) and Nextera® XT index kit (Illumina, San Diego, CA, United States) or Nextera® DNA Flex/ DNA Prep (Illumina, San Diego, CA, United States) or Truseq® DNA (Illumina, San Diego, CA, United States) from 2µg of sample with an addition of 1% of PhiX. Bacterial whole genome sequencing was conducted on MiSeq® or NextSeq® or HiSEQ-1500® (Illumina, San Diego, CA, United States) platforms with a read length of 2 x 300 bp, 2 x 250 bp or 2 x 150 bp.

**Statistics: Pagel lambda phylogenetic analysis.** Pagel lambda values were calculated from the phylogenetic tree obtained from the variant calling analysis using phylosig from phytools v0.7-80 R package (8). In the model, the lambda value transforms a tree by reducing and compressing the inner branches relative to the tip branches which remain unchanged. The lambda value can range from 1 (no transformation) to 0 (all tip branches equal in length and all inner branches equal to 0). When the estimated parameter lambda is equal to 1, the model corresponds to a Brownian motion model evidencing a high phylogenetic signal (i.e., random genetic drift). In contrast, a lambda value less than 1 suggests that shared history had less influence on trait values at the tips. Finally, a lambda value equals to 0 suggests the absence of a phylogenetic signal on trait distributions (i.e., traits evolve independently of phylogeny). Lambda values were calculated over the entire data collection and within the four geographic origins. An additional analysis was performed by randomly subsampling 1000 times the strains from the cohorts with IE/non-IE SAB ratios different from 1 to obtain an overall IE/non-IE SAB ratio set to 1. Each of the lambda values was compared to the distribution obtained by randomly assigning the response variable (i.e., IE or non-IE SAB) 500 times.

**Statistics: Gene and ncRNA analysis by LASSO regression.** The first step in building our lasso model was to divide our data into two subsets (i.e., the training set representing 70% of the data and the validation set 30%) while maintaining the ratio of IE/non-IE SAB. To account for bias due to the different geographical origins, a variable "origin" was added to the matrix of presence/absence of genes and ncRNAs. The main idea was to find the optimal penalty parameter called “lambda”. To this end, a range of values was defined from 10^-3 to 10 in 0.1 steps. To determine which value to use for lambda, we performed a cross-validation of 215 folds (corresponding to 1/3 of the training set) and identified the lambda value that produced the lowest mean square error (MSE). The best lambda value was then used in a final model. To examine the robustness of the model, Cohen's Kappa coefficient calculated from the best model prediction was compared to the distribution obtained by the 500 predictions of the random assignment of the observed responses while maintaining the link between the phenotype and the origin variable. As for phylogenetic signal identification, this entire procedure was computed 1000 times with novel subsampling for training and validation sets.

### Text S2: Supplementary results

**Pagel lambda analysis on SNPs.** No significant phylogenetic signal could be highlighted in the Denmark and US cohorts since lambdas were either very low (US with an IE/non-IE SAB ratio at 1.00 and lambda at 0.0985 (true phenotypic assignation) vs 0.0285 (random phenotypic permutation) [95% CI: 0.0261, 0.0309]) (t-test, p-value < 2.2e-16) or smaller than the one calculated through random permutations (DK with an IE/non-IE SAB ratio at 0.99 and lambda at 0.9600 (true) vs 0.9781 (random permutation) [95% CI: 0.9777, 0.9785]) (t-test, p-value = 1). In contrast, lambdas values obtained for the two Australasian cohorts (0.8392 (true) vs 0.1985 (random permutation) [95% CI: 0.1888, 0.2081] (t-test, p-value < 2.2e-16)) and the French cohort (0.9377 (true) vs 0.8900 (random permutation) [95% CI: 0.8870, 0.8931] (t-test, p-value < 2.2e-16)) were both closer to 1 and higher than the ones calculated after permutations (Table S2). However, these two cohorts presented an unbalanced repartition in favour of non-IE SAB for AUS/NZ (ratio 0.11) or IE for FR (ratio 1.24), limiting the relevance of the potential signal identified. The impact of strain repartition was therefore assessed by randomly subsampling AUS/NZ and FR cohorts to adjust their IE/non-IE SAB ratio to 1 and then combined to the two other cohorts. No significant lambda values were obtained in these conditions [95% CI: 0.6679, 0.9338] (Fig. S3), confirming the absence of relevant phylogenetic signal in the AUS/NZ and FR cohorts and globally in the whole collection.

**K-mer analysis**. Nine overlapping k-mer sequences issued from 3 analyses were found statistically significant between the two infection types. These nine-overlapping k-mer sequences were mapped on TCH60 (Refseq: NC8017342.1) and all nine matched between a rRNA-5S and tRNA-Asn (1092881 - 1092964bp) genomic region. Each significant k-mer had other matches in TCH60 due to the number of rRNA-5S and tRNA-Asn present in the genome. To further investigate this result, the 9 k-mer overlapping sequences were concatenated to obtain a sequence of 84 nucleotides. This sequence was then blasted against each of the 924 assemblies by setting the coverage and identity thresholds to 80%. This sequence is extremely prevalent in all strains sequenced at 2x250 bp and 2x300 bp (345 / 347 strains); it is found with a lower frequency in strains sequenced at 2x150 bp

(Table S3), which is probably related to differences in sequencing depth between the different cohorts. Indeed, if we test independently each pair of cohort-reads length against the IE/non-IE phenotypes, none of the associations were significant (Table S4). In total, this signal cannot be considered as significant and must be considered as artefactual.

Table S1. Clonal complexes distribution among infectious endocarditis (IE) and bacteremia (non-IE SAB) patients.

| CC | IE (%)<br>n = 274 (29.65%) | non-IE SAB (%)<br>n = 650 (70.35%) | Total (%)<br>n = 924 | Ratio<br>IE / non-IE SAB |
|---|---|---|---|---|
| CC30 | 52 (19.0%) | 94 (14.5%) | 146 (15.8%) | 0.55 |
| CC45 | 52 (19.0%) | 87 (13.4%) | 139 (15.0%) | 0.60 |
| CC8 | 25 (9.1%) | 91 (14.0%) | 116 (12.6%) | 0.27 |
| CC5 | 27 (9.9%) | 87 (13.4%) | 114 (12.3%) | 0.31 |
| CC1 | 34 (12.4%) | 63 (9.7%) | 97 (10.5%) | 0.54 |
| CC15 | 27 (9.9%) | 47 (7.2%) | 74 (8.0%) | 0.57 |
| CC22 | 6 (2.2%) | 42 (6.5%) | 48 (5.2%) | 0.14 |
| CC78 | 5 (1.8%) | 32 (4.9%) | 37 (4.0%) | 0.16 |
| CC12 | 10 (3.6%) | 14 (2.2%) | 24 (2.6%) | 0.71 |
| CC20 | 4 (1.5%) | 13 (2.0%) | 17 (1.8%) | 0.31 |
| CC121 | 3 (1.1%) | 12 (1.8%) | 15 (1.6%) | 0.25 |
| CC59 | 2 (0.7%) | 11 (1.7%) | 13 (1.4%) | 0.18 |
| CC97 | 4 (1.5%) | 9 (1.4%) | 13 (1.4%) | 0.44 |
| CC25 | 4 (1.5%) | 7 (1.1%) | 11 (1.2%) | 0.57 |
| CC101 | 3 (1.1%) | 8 (1.2%) | 11 (1.2%) | 0.38 |
| CC7 | 4 (1.5%) | 6 (0.9%) | 10 (1.1%) | 0.67 |
| CC93 | 0 (0.0%) | 10 (1.5%) | 10 (1.1%) | 0.00 |
| CC398 | 6 (2.2%) | 3 (0.5%) | 9 (1.0%) | 2.00 |
| CC509 | 1 (0.4%) | 3 (0.5%) | 4 (0.4%) | 0.33 |
| CC182 | 1 (0.4%) | 2 (0.3%) | 3 (0.3%) | 0.50 |
| CC291 | 0 (0.0%) | 3 (0.5%) | 3 (0.3%) | 0.00 |
| CC50 | 1 (0.4%) | 1 (0.2%) | 2 (0.2%) | 1.00 |
| CC123 | 0 (0.0%) | 2 (0.3%) | 2 (0.2%) | 0.00 |
| CC395 | 2 (0.7%) | 0 (0.0%) | 2 (0.2%) | - |

| | | | | |
|---|---|---|---|---|
| CC80 | 0 (0.0%) | 1 (0.2%) | 1 (0.1%) | 0.00 |
| CC152 | 0 (0.0%) | 1 (0.2%) | 1 (0.1%) | 0.00 |
| CC1093 | 0 (0.0%) | 1 (0.2%) | 1 (0.1%) | 0.00 |
| CC2867 | 1 (0.4%) | 0 (0.0%) | 1 (0.1%) | - |

Since the whole collection had been enriched in CC45 and CC3O strains from Denmark and the USA respectively, no statistics was performed on this distribution.

Table S2. Pagel lambda phylogenetic signal measurement within the entire strain collection and within each of the four geographical origins.

| | | Pagel lambda value | | | | CI 95% from 500 permutations | | |
|---|---|---|---|---|---|---|---|---|
| Cohort | Ratio IE/non-IE SAB | Lambda | logL (Lambda) | LR (Lambda=0) | P-value (LR test) | lower | mean | upper |
| Global | 0.42 | 0.9441 | -750.247 | 147.546 | 5.9625e-34 | 0.7329 | 0.7397 | 0.7466 |
| AUS/NZ | 0.11 | 0.8392 | -173.424 | 17.5224 | 2.8395e-05 | 0.1888 | 0.1985 | 0.2081 |
| DK | 0.99 | 0.9600 | -309.762 | 64.3774 | 1.0273e-15 | 0.9777 | 0.9781 | 0.9785 |
| FR | 1.24 | 0.9377 | -141.061 | 22.0126 | 2.7087e-6 | 0.8870 | 0.8900 | 0.8931 |
| US | 1.00 | 0.0985 | -29.5175 | 0.4037 | 0.5252 | 0.0261 | 0.0285 | 0.0309 |

Table S3. Result of the presence/absence counts of the concatenation of the nine genomic sequences found in the DBGWAS analysis.

| | | Sequencing length | | |
|---|---|---|---|---|
| | Cohort | 2x150bp (n=577) | 2x250bp (n=291) | 2x300bp (n=56) |
| Presence in IE strains | AUS/NZ (n=49) | 37 / 41 | 4 / 4 | 4 / 4 |
| | DK (n=133) | 4 / 46 | 86 / 86 | 1 / 1 |
| | FR (n=72) | 9 / 66 | 6 / 6 | 0 / 0 |
| | US (n=20) | 0 / 1 | 19 / 19 | 0 / 0 |
| | Global (n=274) | 50 / 154 | 115 / 115 | 5 / 5 |
| Presence in non-IE SAB strains | AUS/NZ (n=438) | 290 / 321 | 65 / 66 | 50 / 51 |
| | DK (n=134) | 7 / 50 | 84 / 84 | 0 / 0 |
| | FR (n=58) | 6 / 52 | 6 / 6 | 0 / 0 |
| | US (n=20) | 0 / 0 | 20 / 20 | 0 / 0 |
| | Global (n=650) | 303 / 423 | 175 / 176 | 50 / 51 |

The 84-nucleotide sequence was blasted in each of the genome assemblies with parameters set at 80% coverage and identity.

Table S4. Fisher exact tests between IE and non-IE SAB strains on the presence/absence of the 84-nucleotide sequence found in the DBGWAS analysis.

| | 2x150bp | | 2x250bp | | 2x300bp | | [illegible] |
|---|---|---|---|---|---|---|---|
| Cohort | Odds Ratio (95% CI) | P-value | Odds Ratio (95% CI) | P-value | Odds Ratio (95% CI) | P-value | [illegible] |
| AUS/NZ | 1.0113 [0.2456, 3.1008] | 1 | 0 [0, 637.1041] | 1 | 0 [0, 493.424] | 1 | 1.090[illegible] |
| DK | 1.6999 [0.3973, 8.5206] | 0.528 | 0 [0, Inf] | 1 | 0 [0, Inf] | 1 | 0.976[illegible] |
| FR | 0.8274 [0.2249, 2.8264] | 0.7876 | 0 [0, Inf] | 1 | 0 [0, Inf] | 1 | 0.991[illegible] |
| US | 0 [0, Inf] | 1 | 0 [0, Inf] | 1 | 0 [0, Inf] | 1 | In[illegible] |
| Global | 5.2349 [3.4642, 7.9926] | < 2.2e-16 | 0 [0, 59.6323] | 1 | 0 [0, 395.3488] | 1 | 0[illegible] |

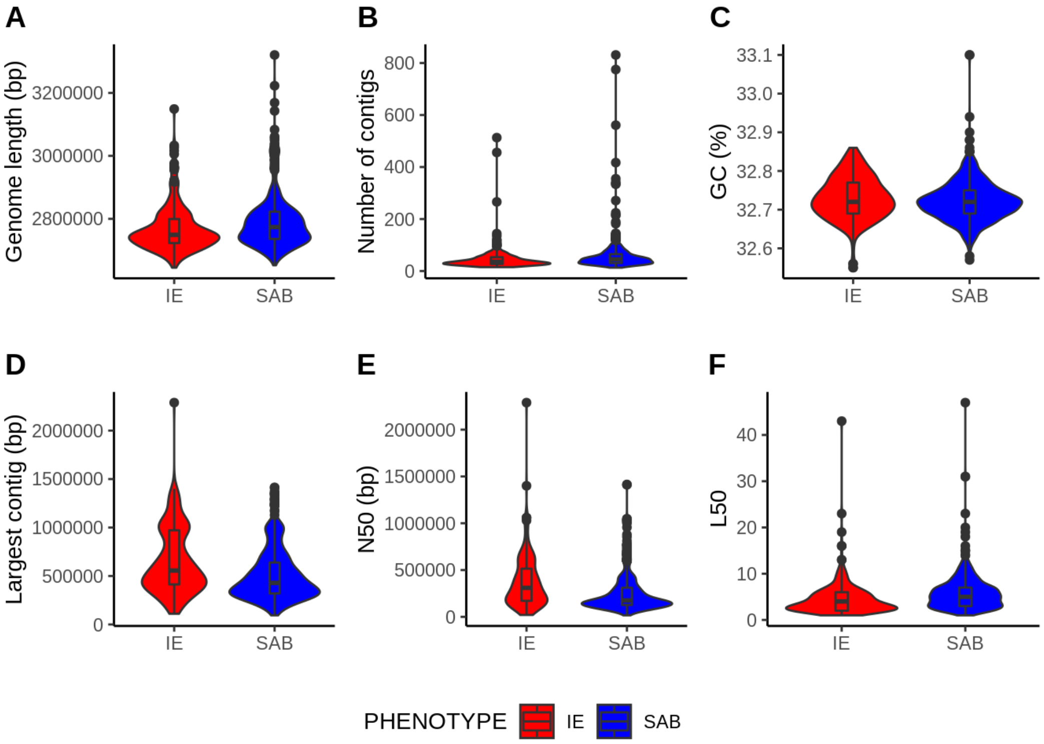


Figure S1. Summary of the 924 assemblies in the *Staphylococcus aureus* collection. A) Genome length in bp, B) Number of contigs, C) Percentage of GC, D) Largest contig in bp, E) N50 in bp. N50 refers to the sequence length of the contig, when they are summed from the longest to the smallest, which overtakes half of the total genome length. F) L50 is defined as the number of contigs required that exceed half of the genome size.

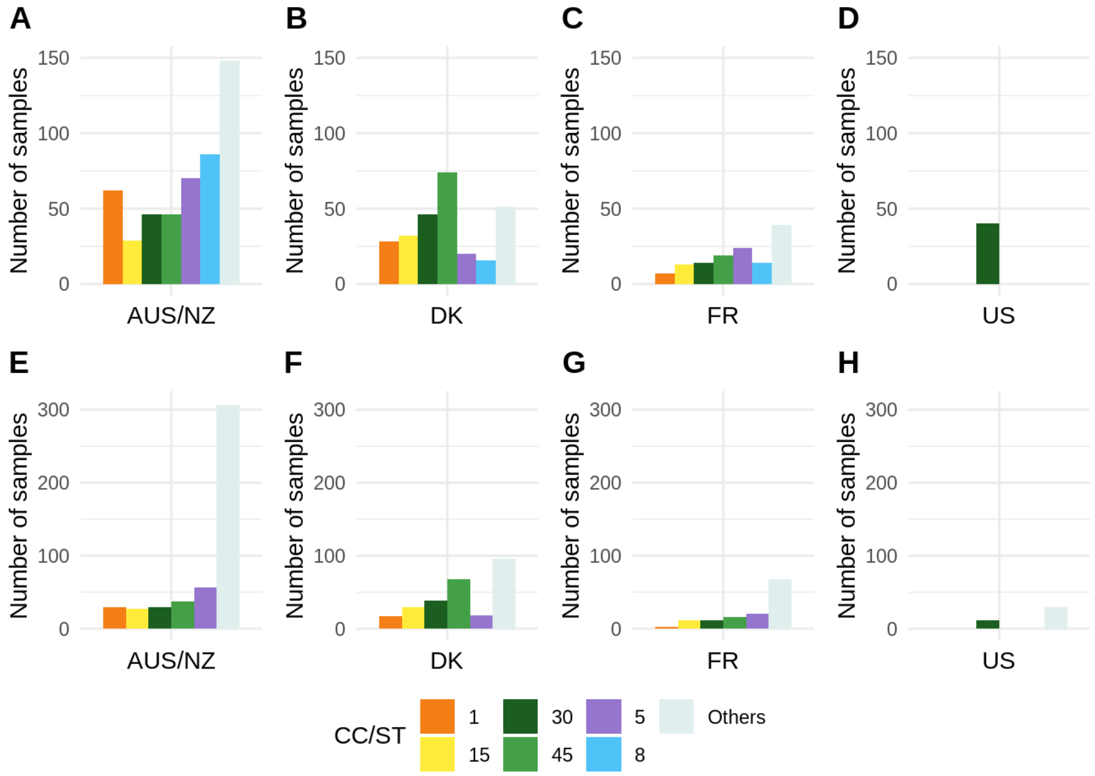


Figure S2. The six largest clonal complexes (A – D) and the five largest sequence types (E – H) distributions among the geographical origin of the sample: AUS/NZ (A, E), DK (B, F), FR (C, G) and US (D, H). The six largest clonal complexes and five largest sequence types are colored as followed: CC/ST1: orange; CC/ST15: yellow; CC/ST30: dark green; CC/ST45: light green; CC/ST5: purple; CC8: light blue; other CCs or STs are pooled together (grey).

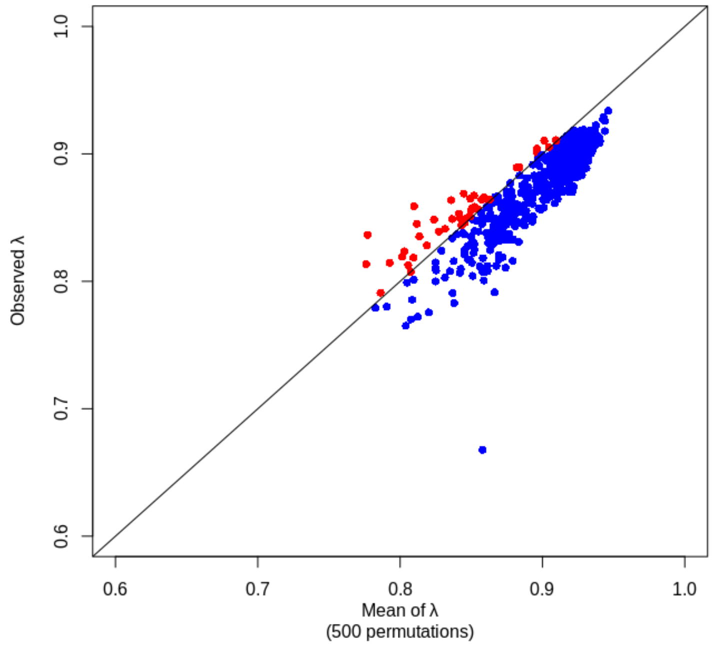


Figure S3. Investigation of a common heritable trait for IE vs non-IE SAB by comparing the observed Pagel's lambda measurements based on 15 436 SNPs, with the distribution obtained after 500 permutations. The cohorts with IE/non-IE SAB ratios different from 1 were randomly subsampled 1000 times to obtain an overall IE/non-IE SAB ratio set to 1, and combined to the 2 other cohorts. Each lambda value was compared to the mean of the distribution obtained by randomly assigning the response variable 500 times. Red dots (4.4%) and blue dots (95.6%) correspond to observed lambda values that are greater or lower than the average of the 500 lambda values obtained after permutations. No relevant phylogenetic signal in the whole collection could discriminate IE vs non-IE SAB.

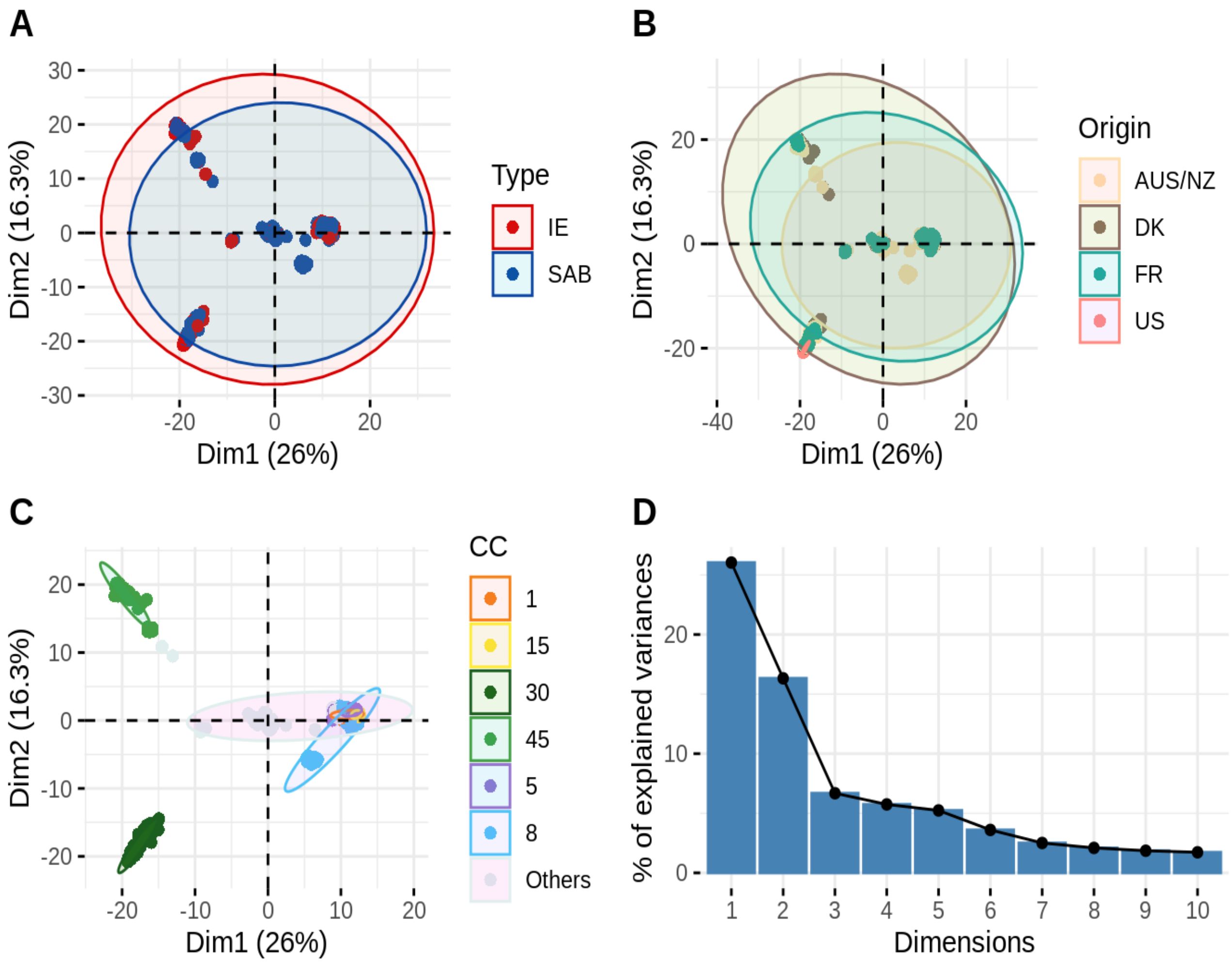


Figure S4. Two individual principal component analysis plots using the two first principal components (A – C), focusing on 4757 coding sequences and non-coding RNAs shared by more than five percent and less than 95 percent of the 924 *S. aureus* strains. Each dot corresponds to one sample and is coloured according to (A) infection type, IE (red) and non-IE SAB (blue); (B) geographical origin, AUS/NZ (beige), DK (brown), FR (turquoise) and US (pink); (C) clonal complex membership, the six largest clonal complexes with their sequence types are coloured as followed: CC1: orange; CC15: yellow; CC30: dark green; CC45: light green; CC5: purple; CC8: light blue; other CCs are pooled together (grey). D) Eigenvalues.

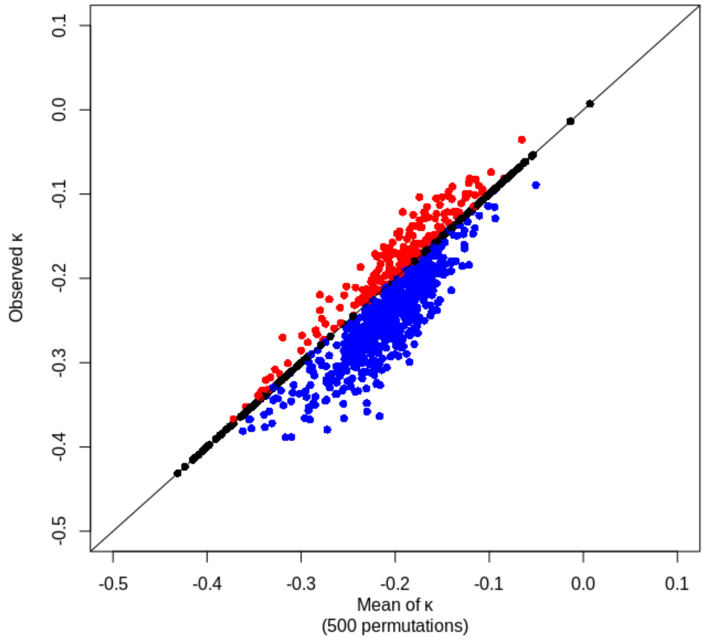


Figure S5. Investigation of an association between the presence / absence of 4757 coding sequences and ncRNAs with IE vs non-IE SAB by the comparison of Cohen's Kappa coefficient measurements with the distribution obtained by 500 permutations of Kappa values from LASSO regressions. Cohen's Kappa coefficient measures the agreement between the prediction and the actual state after adjusting this value for what could be expected from chance alone. It was computed 1000 times with novel subsampling for training and validation sets. Each kappa value was compared to the mean of the distribution obtained by randomly assigning the response variable 500 times while maintaining the link between the phenotype and the geographic origin variable. Red dots (19.8%), blue dots (60.2%) and black dots (20%) correspond to observed kappa values that are greater, lower or equal than the average of the 500 kappa values obtained after permutations, respectively. Cohen's Kappa coefficient measurements remain far from significance (=1), assessing the lack of signal discriminating IE vs non-IE SAB.

#### DATA SUMMARY

1. 130 strains (72 IE and 58 non-IE SAB cases) from the French national prospective multicenter cohort VIRSTA. Read data is deposited in the European Nucleotide Archive (ENA): study accession numbers PRJEB48298 and PRJEB49354 (http://www.ebi.ac.uk/ena/data/view/PRJEB48298, http://www.ebi.ac.uk/ena/data/view/PRJEB49354).

2. 26 strains (13 IE and 13 non-IE SAB) from the Danish National *Staphylococcus aureus* Bacteremia Repository (Statens Serum Institut), 40 strains (20 IE and 20 non-IE SAB) obtained from the *Staphylococcus aureus* Bacteremia Group (SABG) biorepository at Duke University. Read data is deposited in the European Nucleotide Archive (ENA): study accession number will be provided upon request.

3. 487 strains (49 IE and 438 non-IE SAB) from two Australasian cohorts of SAB, namely the vancomycin substudy of the Australian and New Zealand Cooperative on Outcome of Staphylococcal Sepsis (ANZCOSS) and the Vancomycin Efficacy in Staphylococcal Sepsis in Australasia (VANESSA) study. Read data is deposited in the European Nucleotide Archive (ENA): study accession numbers PRJEB22792 (http://www.ebi.ac.uk/ena/data/view/PRJEB22792).

4. 241 strains (120 IE and 121 non-IE SAB) collected during two prospective studies in the Denmark and the US. Read data is deposited in the European Nucleotide Archive (ENA): study accession number ERP023934 (http://www.ebi.ac.uk/ena/data/view/ERP023934).

5. Code, R-scripts and data files used when running the scripts can be accessed at Sourceforge, https://sourceforge.net/projects/###### /files/.
