## Supplementary material for "All *Staphylococcus aureus* bacteraemia strains have the potential to cause infective endocarditis: results of GWAS and experimental animal studies": Summary-genomes-assemblies

| FILE | PHENOTYPE | GEOGRAPHICAL ORIGIN | ANIMAL EXPERIMENT | CC | ST | CONTIGS NUMBER (>=0bp) | LENGTH (>=0bp) | CONTIGS NUMBER (>=500bp) | LENGTH (>=500bp) | LARGEST CONTIG | N50 | L50 | GC (%) | SEQUENCING | INFORMATIONS |
|---|---|---|---|---|---|---|---|---|---|---|---|---|---|---|---|
| 1701 | SAB | US | | 30 | 36 | 60 | 2961817 | 54 | 2959355 | 422199 | 151221 | 7 | 32.70 | 2x250bp | |
| 1702 | SAB | US | | 30 | 30 | 54 | 2864851 | 46 | 2861809 | 409007 | 151136 | 6 | 32.75 | 2x250bp | |
| 1703 | SAB | US | | 30 | 30 | 74 | 2785212 | 67 | 2782581 | 297420 | 85255 | 9 | 32.75 | 2x250bp | |
| 1704 | SAB | US | | 30 | 30 | 102 | 2877363 | 76 | 2868830 | 330115 | 67618 | 11 | 32.74 | 2x250bp | |
| 1705 | SAB | US | | 30 | 30 | 59 | 2777035 | 44 | 2772015 | 316949 | 150889 | 7 | 32.70 | 2x250bp | |
| 1706 | SAB | US | | 30 | 36 | 73 | 2954528 | 56 | 2948606 | 422357 | 134600 | 7 | 32.69 | 2x250bp | |
| 1707 | SAB | US | | 30 | 543 | 48 | 2808272 | 45 | 2807043 | 316699 | 151221 | 7 | 32.74 | 2x250bp | |
| 1708 | SAB | US | | 30 | 36 | 69 | 2958543 | 58 | 2954149 | 419207 | 121883 | 7 | 32.71 | 2x250bp | |
| 1709 | SAB | US | | 30 | 346 | 50 | 2960499 | 47 | 2959342 | 422155 | 170884 | 5 | 32.71 | 2x250bp | |
| 1710 | SAB | US | | 30 | 30 | 53 | 2866511 | 49 | 2864935 | 337199 | 124223 | 7 | 32.75 | 2x250bp | |
| 1711 | SAB | US | | 30 | 346 | 81 | 2970370 | 52 | 2959345 | 345800 | 151096 | 7 | 32.70 | 2x250bp | |
| 1712 | SAB | US | | 30 | 30 | 48 | 2822852 | 38 | 2819117 | 422158 | 171184 | 5 | 32.72 | 2x250bp | |
| 1713 | SAB | US | | 30 | 36 | 54 | 2961211 | 47 | 2958401 | 422152 | 150878 | 7 | 32.70 | 2x250bp | |
| 1714 | SAB | US | | 30 | 36 | 93 | 2964346 | 68 | 2955377 | 422267 | 130389 | 6 | 32.77 | 2x250bp | |
| 1715 | SAB | US | | 30 | 36 | 73 | 2968437 | 50 | 2959525 | 422214 | 142273 | 6 | 32.71 | 2x250bp | |
| 1716 | SAB | US | | 30 | 346 | 53 | 2958274 | 46 | 2955342 | 392538 | 139391 | 7 | 32.71 | 2x250bp | |
| 1717 | SAB | US | | 30 | 30 | 59 | 2828132 | 51 | 2825035 | 321269 | 151067 | 7 | 32.72 | 2x250bp | |
| 1718 | SAB | US | | 30 | 36 | 54 | 2961965 | 48 | 2959576 | 422214 | 208782 | 5 | 32.71 | 2x250bp | |
| 1719 | SAB | US | | 30 | 36 | 50 | 2879783 | 45 | 2877790 | 300845 | 171334 | 7 | 32.72 | 2x250bp | |
| 1720 | SAB | US | | 30 | 30 | 43 | 2811316 | 39 | 2809789 | 479409 | 167048 | 6 | 32.73 | 2x250bp | |
| 1721 | IE | US | | 30 | 36 | 59 | 2961794 | 53 | 2959256 | 397764 | 150636 | 6 | 32.71 | 2x250bp | |
| 1722 | IE | US | | 30 | 36 | 50 | 2920563 | 44 | 2918180 | 422214 | 151233 | 7 | 32.75 | 2x250bp | |
| 1723 | IE | US | | 30 | 30 | 107 | 2904126 | 82 | 2895484 | 314074 | 154334 | 7 | 32.76 | 2x250bp | |
| 1724 | IE | US | | 30 | 36 | 43 | 2877561 | 38 | 2875569 | 397937 | 208813 | 5 | 32.72 | 2x250bp | |
| 1725 | IE | US | | 30 | 36 | 60 | 2957659 | 46 | 2952382 | 422114 | 209715 | 5 | 32.70 | 2x250bp | |
| 1726 | IE | US | | 30 | 36 | 74 | 2900469 | 62 | 2895985 | 392298 | 118536 | 7 | 32.65 | 2x150bp | |
| 1727 | IE | US | | 30 | 346 | 57 | 2962859 | 49 | 2959814 | 422177 | 151096 | 6 | 32.70 | 2x250bp | |
| 1728 | IE | US | | 30 | 346 | 80 | 2967605 | 57 | 2958682 | 392603 | 151219 | 6 | 32.71 | 2x250bp | |
| 1729 | IE | US | | 30 | 346 | 55 | 2963350 | 47 | 2960193 | 422199 | 151220 | 6 | 32.70 | 2x250bp | |
| 1730 | IE | US | | 30 | 346 | 62 | 2955928 | 50 | 2951643 | 418449 | 151000 | 6 | 32.70 | 2x250bp | |
| 1731 | IE | US | | 30 | 346 | 91 | 2910094 | 77 | 2904667 | 417811 | 105830 | 7 | 32.68 | 2x250bp | |
| 1732 | IE | US | | 30 | 30 | 51 | 2817713 | 46 | 2815665 | 413724 | 163503 | 5 | 32.77 | 2x250bp | |
| 1733 | IE | US | | 30 | 346 | 68 | 2962274 | 62 | 2959805 | 417568 | 150607 | 6 | 32.77 | 2x250bp | |
| 1734 | IE | US | | 30 | 36 | 96 | 2913435 | 83 | 2908432 | 337964 | 105103 | 9 | 32.69 | 2x250bp | |
| 1735 | IE | US | | 30 | 346 | 70 | 2916976 | 62 | 2913873 | 392543 | 111547 | 8 | 32.76 | 2x250bp | |
| 1736 | IE | US | | 30 | 36 | 88 | 2976032 | 68 | 2968821 | 421994 | 100773 | 9 | 32.74 | 2x250bp | |
| 1737 | IE | US | | 30 | 36 | 73 | 2911990 | 62 | 2908155 | 422102 | 142171 | 6 | 32.74 | 2x250bp | |
| 1738 | IE | US | | 30 | 30 | 55 | 2831174 | 48 | 2828475 | 392082 | 151068 | 6 | 32.74 | 2x250bp | |
| 1739 | IE | US | | 30 | 36 | 48 | 2921553 | 43 | 2919569 | 417677 | 180508 | 5 | 32.76 | 2x250bp | |
| 1740 | IE | US | | 30 | 36 | 110 | 3016427 | 65 | 2999098 | 422339 | 171084 | 6 | 32.72 | 2x250bp | |
| 41273 | IE | DK | | 45 | 45 | 36 | 2806522 | 27 | 2802954 | 963307 | 601043 | 2 | 32.84 | 2x250bp | Danish Repository |
| 41885 | IE | DK | | 15 | 15 | 34 | 2692096 | 25 | 2688259 | 567974 | 438060 | 3 | 32.72 | 2x250bp | Danish Repository |
| 43139 | IE | DK | | 45 | 45 | 29 | 2743207 | 24 | 2741062 | 983223 | 422985 | 2 | 32.80 | 2x250bp | Danish Repository |
| 46235 | IE | DK | | 45 | 45 | 27 | 2695011 | 20 | 2692097 | 1282729 | 985681 | 2 | 32.80 | 2x250bp | Danish Repository |
| 46524 | IE | DK | | 45 | 45 | 19 | 2759466 | 16 | 2758075 | 1025373 | 874228 | 2 | 32.80 | 2x250bp | Danish Repository |
| 50368 | IE | DK | | 45 | 45 | 30 | 2741642 | 21 | 2737824 | 726365 | 315266 | 3 | 32.81 | 2x250bp | Danish Repository |
| 55189 | IE | DK | | 45 | 45 | 26 | 2700080 | 21 | 2697919 | 807824 | 618716 | 2 | 32.81 | 2x250bp | Danish Repository |
| 56603 | IE | DK | | 395 | 395 | 33 | 2756048 | 20 | 2750973 | 1006748 | 575168 | 2 | 32.76 | 2x250bp | Danish Repository |
| 60668 | IE | DK | | 45 | 45 | 20 | 2742757 | 19 | 2742346 | 1039379 | 622628 | 2 | 32.85 | 2x250bp | Danish Repository |
| 60825 | SAB | DK | | 45 | 45 | 30 | 2738532 | 26 | 2736762 | 1035763 | 504251 | 2 | 32.80 | 2x250bp | Danish Repository |
| 62466 | IE | DK | | 5 | 5 | 27 | 2754247 | 22 | 2752166 | 1023362 | 702504 | 2 | 32.78 | 2x250bp | Danish Repository |
| 65934 | SAB | DK | | 45 | 45 | 28 | 2773760 | 21 | 2770830 | 1128039 | 1029417 | 2 | 32.76 | 2x250bp | Danish Repository |
| 66783 | SAB | DK | | 45 | 45 | 25 | 2696978 | 20 | 2694839 | 608037 | 438809 | 3 | 32.78 | 2x250bp | Danish Repository |
| 72061 | IE | DK | | 45 | 45 | 72 | 2675230 | 68 | 2673661 | 184775 | 68454 | 11 | 32.78 | 2x150bp | |
| 73582 | IE | DK | | 45 | 45 | 30 | 2742459 | 24 | 2740096 | 607649 | 338645 | 3 | 32.81 | 2x250bp | Danish Repository |
| 75171 | SAB | DK | | 45 | 45 | 36 | 2701498 | 26 | 2698207 | 471625 | 321292 | 4 | 32.81 | 2x250bp | Danish Repository |
| 76045 | IE | DK | | 1 | 3384 | 18 | 2755201 | 10 | 2752013 | 1399962 | 1399962 | 1 | 32.73 | 2x250bp | |
| 76233 | IE | DK | | 30 | 30 | 64 | 2870476 | 50 | 2864857 | 394526 | 151038 | 7 | 32.76 | 2x250bp | |
| 76447 | IE | DK | | 7 | 7 | 42 | 2716351 | 33 | 2713408 | 324800 | 259214 | 5 | 32.72 | 2x150bp | |
| 76603 | IE | DK | | 12 | 12 | 23 | 2701850 | 16 | 2699004 | 709387 | 502243 | 3 | 32.75 | 2x250bp | |
| 76907 | IE | DK | | 45 | 45 | 30 | 2711449 | 22 | 2708049 | 674812 | 553079 | 3 | 32.79 | 2x250bp | |
| 76953 | IE | DK | | 45 | 45 | 24 | 2750709 | 19 | 2748514 | 991674 | 608013 | 2 | 32.83 | 2x250bp | |
| 77105 | IE | DK | | 59 | 59 | 56 | 2737863 | 40 | 2731260 | 317376 | 169527 | 6 | 32.76 | 2x250bp | |
| 77521 | IE | DK | Mice | 5 | 5 | 33 | 2778171 | 24 | 2774586 | 1033877 | 734126 | 2 | 32.74 | 2x250bp | |
| 77743 | IE | DK | | 8 | 8 | 17 | 2741858 | 12 | 2739682 | 1081251 | 1025687 | 2 | 32.66 | 2x250bp | |
| 77847 | IE | DK | | 45 | 4229 | 45 | 2761246 | 37 | 2758021 | 549727 | 182006 | 5 | 32.79 | 2x250bp | |
| 77871 | IE | DK | | 25 | 25 | 16 | 2770489 | 15 | 2770038 | 979177 | 748265 | 2 | 32.71 | 2x250bp | |
| 78369 | SAB | DK | | 45 | 45 | 29 | 2757394 | 17 | 2752920 | 1244407 | 597892 | 2 | 32.80 | 2x250bp | Danish Repository |
| 78577 | IE | DK | | 15 | 582 | 33 | 2730302 | 25 | 2727042 | 599619 | 320352 | 3 | 32.74 | 2x250bp | |
| 78586 | SAB | DK | | 30 | 30 | 107 | 2833560 | 58 | 2815884 | 486136 | 136296 | 6 | 32.73 | 2x250bp | Danish Repository |
| 78611 | IE | DK | | 15 | 15 | 40 | 2644869 | 30 | 2641101 | 427120 | 234606 | 5 | 32.72 | 2x150bp | |
| 78790 | SAB | DK | | 45 | 45 | 50 | 2780364 | 34 | 2774590 | 993365 | 498153 | 2 | 32.76 | 2x250bp | Danish Repository |
| 80033 | SAB | DK | | 45 | 45 | 36 | 2684976 | 23 | 2680090 | 898488 | 601734 | 2 | 32.79 | 2x250bp | Danish Repository |
| 80330 | SAB | DK | | 45 | 45 | 28 | 2761251 | 23 | 2759131 | 616295 | 421908 | 3 | 32.78 | 2x250bp | Danish Repository |
| 81401 | IE | DK | | 12 | 12 | 19 | 2706599 | 16 | 2705281 | 974985 | 665165 | 2 | 32.75 | 2x250bp | |
| 81867 | IE | DK | | 15 | 15 | 58 | 2712060 | 52 | 2709571 | 232106 | 127260 | 8 | 32.69 | 2x150bp | |
| 82003 | IE | DK | | 1 | 188 | 70 | 2719381 | 60 | 2715765 | 199882 | 82187 | 11 | 32.67 | 2x150bp | |

| | | | | | | | | | | | | | | | |
|---|---|---|---|---|---|---|---|---|---|---|---|---|---|---|---|
| 82259 | IE | DK | | 45 | 45 | 21 | 2747220 | 20 | 2746732 | 993477 | 385012 | 2 | 32.82 | 2x250bp | |
| 82273 | SAB | DK | | 45 | 45 | 31 | 2749051 | 20 | 2744830 | 612777 | 421584 | 3 | 32.80 | 2x250bp | Danish Repository |
| 82513 | IE | DK | | 30 | 30 | 53 | 2817745 | 46 | 2814990 | 421072 | 150511 | 6 | 32.73 | 2x250bp | |
| 82649 | IE | DK | | 30 | 30 | 31 | 2758053 | 27 | 2756589 | 467511 | 197072 | 5 | 32.74 | 2x250bp | |
| 84551 | IE | DK | | 5 | 6 | 36 | 2822050 | 31 | 2819822 | 645089 | 506315 | 3 | 32.76 | 2x250bp | |
| 85243 | IE | DK | | 1 | 1 | 52 | 2722655 | 48 | 2721196 | 219951 | 115322 | 9 | 32.69 | 2x150bp | |
| 85756 | SAB | DK | | 45 | 10007* | 25 | 2738399 | 23 | 2737534 | 433955 | 393440 | 4 | 32.84 | 2x250bp | ‹nown ST; Danish Repository |
| 85820 | SAB | DK | | 45 | 45 | 47 | 2717364 | 35 | 2712967 | 599908 | 135792 | 5 | 32.77 | 2x250bp | Danish Repository |
| 85835 | IE | DK | | 12 | 12 | 23 | 2723185 | 18 | 2721099 | 1296367 | 661287 | 2 | 32.71 | 2x250bp | |
| 87677 | IE | DK | | 45 | 3396 | 36 | 2741071 | 21 | 2735662 | 526892 | 460424 | 3 | 32.80 | 2x250bp | Danish Repository |
| 88833 | SAB | DK | | 45 | 45 | 31 | 2747647 | 27 | 2745989 | 602463 | 283826 | 4 | 32.82 | 2x250bp | Danish Repository |
| 89969 | IE | DK | | 1 | 188 | 23 | 2737989 | 15 | 2734687 | 753489 | 566420 | 3 | 32.70 | 2x250bp | |
| 90547 | IE | DK | | 30 | 30 | 47 | 2807057 | 41 | 2804684 | 422175 | 164890 | 6 | 32.75 | 2x250bp | |
| 90913 | IE | DK | | 15 | 15 | 25 | 2692881 | 22 | 2691492 | 611419 | 307616 | 4 | 32.69 | 2x250bp | |
| 90973 | IE | DK | | 1 | 188 | 84 | 2758336 | 74 | 2754576 | 236388 | 81872 | 11 | 32.72 | 2x150bp | |
| 91589 | IE | DK | | 8 | 8 | 39 | 2817625 | 29 | 2813373 | 896807 | 867837 | 2 | 32.66 | 2x250bp | |
| 92039 | IE | DK | | 509 | 509 | 99 | 2844653 | 84 | 2838393 | 224506 | 81830 | 12 | 32.72 | 2x250bp | |
| 92381 | IE | DK | | 8 | 8 | 64 | 2743685 | 53 | 2739775 | 479570 | 125891 | 6 | 32.65 | 2x150bp | |
| 93317 | IE | DK | | 45 | 45 | 27 | 2725626 | 25 | 2724727 | 1238217 | 414106 | 2 | 32.78 | 2x250bp | |
| 93877 | IE | DK | | 5 | 6 | 55 | 2765054 | 50 | 2763113 | 351461 | 146334 | 6 | 32.72 | 2x150bp | |
| 94495 | IE | DK | | 15 | 15 | 48 | 2726052 | 43 | 2724090 | 251760 | 123807 | 7 | 32.68 | 2x150bp | |
| 94519 | IE | DK | | 45 | 45 | 27 | 2724351 | 22 | 2722156 | 606529 | 423443 | 3 | 32.77 | 2x250bp | |
| 94572 | IE | DK | | 45 | 45 | 37 | 2697091 | 26 | 2693107 | 525832 | 288696 | 4 | 32.80 | 2x250bp | Danish Repository |
| 95457 | IE | DK | | 1 | 1 | 20 | 2784203 | 13 | 2781386 | 1030108 | 768480 | 2 | 32.70 | 2x250bp | |
| 95703 | IE | DK | | 15 | 15 | 27 | 2728949 | 24 | 2727673 | 668936 | 315021 | 3 | 32.73 | 2x250bp | |
| 100135 | IE | DK | | 15 | 15 | 36 | 2664679 | 31 | 2662519 | 275604 | 193375 | 6 | 32.68 | 2x150bp | |
| 100597 | IE | DK | | 45 | 45 | 25 | 2746709 | 18 | 2743685 | 991390 | 606687 | 2 | 32.82 | 2x250bp | |
| 101291 | IE | DK | Mice | 5 | 5 | 38 | 2837635 | 25 | 2832465 | 1034855 | 419019 | 2 | 32.77 | 2x250bp | |
| 101541 | IE | DK | | 22 | 22 | 48 | 2848826 | 39 | 2845000 | 404927 | 211624 | 5 | 32.68 | 2x250bp | |
| 102965 | IE | DK | | 25 | 25 | 32 | 2766593 | 27 | 2764806 | 428072 | 322747 | 4 | 32.69 | 2x150bp | |
| 103237 | IE | DK | | 45 | 45 | 22 | 2751989 | 19 | 2750683 | 1325101 | 1029385 | 2 | 32.81 | 2x250bp | |
| 103483 | IE | DK | | 59 | 59 | 46 | 2749563 | 30 | 2742917 | 453198 | 282257 | 4 | 32.74 | 2x250bp | |
| 103701 | IE | DK | | 15 | 15 | 36 | 2661889 | 34 | 2661211 | 350946 | 184227 | 6 | 32.70 | 2x150bp | |
| 103767 | IE | DK | | 30 | 30 | 55 | 2782302 | 46 | 2778747 | 369279 | 151219 | 7 | 32.71 | 2x250bp | |
| 103921 | IE | DK | | 182 | 182 | 25 | 2792040 | 16 | 2788464 | 1328195 | 1060503 | 2 | 32.70 | 2x250bp | |
| 103995 | IE | DK | | 30 | 30 | 61 | 2846196 | 49 | 2841255 | 577256 | 177974 | 5 | 32.77 | 2x250bp | |
| 104319 | IE | DK | | 30 | 30 | 60 | 2856332 | 54 | 2853871 | 366125 | 150565 | 6 | 32.77 | 2x250bp | |
| 104343 | IE | DK | | 7 | 7 | 52 | 2770780 | 42 | 2767199 | 466233 | 198379 | 5 | 32.72 | 2x150bp | |
| 105343 | IE | DK | | 25 | 25 | 30 | 2703530 | 26 | 2702020 | 728330 | 475779 | 3 | 32.68 | 2x150bp | |
| 105445 | IE | DK | | 1 | 1 | 144 | 2779084 | 136 | 2776110 | 120215 | 39901 | 23 | 32.66 | 2x150bp | |
| 105547 | IE | DK | | 30 | 30 | 63 | 2821081 | 57 | 2818639 | 426094 | 134775 | 7 | 32.72 | 2x250bp | |
| 106141 | IE | DK | | 12 | 12 | 31 | 2681162 | 30 | 2680800 | 416637 | 187401 | 5 | 32.74 | 2x150bp | |
| 106461 | IE | DK | | 1 | 9 | 33 | 2778857 | 27 | 2776664 | 519726 | 425660 | 3 | 32.70 | 2x150bp | |
| 106533 | IE | DK | | 30 | 30 | 45 | 2823322 | 39 | 2820926 | 394769 | 170799 | 5 | 32.73 | 2x250bp | |
| 107191 | IE | DK | | 22 | 22 | 67 | 2760540 | 52 | 2755124 | 575858 | 150870 | 5 | 32.73 | 2x150bp | |
| 107635 | IE | DK | | 30 | 30 | 83 | 2806125 | 70 | 2801141 | 370324 | 81640 | 10 | 32.72 | 2x150bp | |
| 110231 | IE | DK | | 45 | 508 | 22 | 2780813 | 19 | 2779519 | 981964 | 540453 | 2 | 32.83 | 2x250bp | |
| 110255 | IE | DK | | 30 | 34 | 71 | 2918939 | 60 | 2914534 | 494351 | 164453 | 5 | 32.71 | 2x250bp | |
| 111069 | IE | DK | | 30 | 30 | 47 | 2773152 | 42 | 2771201 | 394628 | 175581 | 6 | 32.71 | 2x250bp | |
| 111189 | IE | DK | | 15 | 15 | 42 | 2683722 | 34 | 2680520 | 426206 | 217880 | 5 | 32.66 | 2x150bp | |
| 111697 | IE | DK | | 8 | 8 | 42 | 2802864 | 37 | 2801232 | 491609 | 148185 | 6 | 32.64 | 2x150bp | |
| 111765 | IE | DK | | 45 | 45 | 37 | 2761128 | 29 | 2757855 | 968371 | 626230 | 2 | 32.71 | 2x250bp | |
| 112005 | IE | DK | | 8 | 8 | 35 | 2747135 | 22 | 2742551 | 794891 | 313320 | 3 | 32.64 | 2x150bp | |
| 112553 | IE | DK | | 97 | 97 | 98 | 2717041 | 79 | 2710228 | 184486 | 68953 | 13 | 32.68 | 2x150bp | |
| 113165 | IE | DK | | 5 | 5 | 40 | 2734304 | 31 | 2730672 | 1022475 | 432726 | 2 | 32.70 | 2x250bp | |
| 113391 | IE | DK | | 1 | 9 | 40 | 2777905 | 34 | 2775710 | 344374 | 192195 | 5 | 32.69 | 2x150bp | |
| 113647 | IE | DK | | 15 | 15 | 33 | 2722911 | 30 | 2721580 | 422952 | 249788 | 5 | 32.70 | 2x150bp | |
| 114435 | IE | DK | | 30 | 34 | 53 | 2813140 | 45 | 2809902 | 408118 | 175519 | 6 | 32.70 | 2x250bp | |
| 114821 | IE | DK | | 45 | 45 | 29 | 2706071 | 22 | 2703381 | 599718 | 529307 | 3 | 32.78 | 2x250bp | |
| 116063 | IE | DK | | 12 | 12 | 24 | 2693460 | 18 | 2691536 | 830396 | 638672 | 2 | 32.73 | 2x150bp | |
| 116693 | IE | DK | | 45 | 45 | 30 | 2723743 | 23 | 2721424 | 1035729 | 421141 | 2 | 32.84 | 2x250bp | |
| 116745 | IE | DK | | 20 | 20 | 40 | 2743312 | 29 | 2739409 | 563808 | 245097 | 4 | 32.67 | 2x150bp | |
| 120725 | IE | DK | | 15 | 15 | 31 | 2690099 | 23 | 2686754 | 445327 | 315073 | 4 | 32.72 | 2x300bp | |
| 120897 | IE | DK | | 30 | 30 | 90 | 2813288 | 72 | 2807129 | 369131 | 134608 | 7 | 32.73 | 2x150bp | |
| 121881 | IE | DK | | 15 | 15 | 39 | 2669134 | 32 | 2666212 | 400561 | 181388 | 6 | 32.68 | 2x150bp | |
| 123327 | SAB | DK | | 1093 | 1093 | 45 | 2779143 | 35 | 2775495 | 397234 | 337649 | 4 | 32.78 | 2x150bp | |
| 123441 | SAB | DK | | 45 | 45 | 23 | 2746651 | 22 | 2746248 | 1035495 | 608121 | 2 | 32.83 | 2x250bp | |
| 124991 | IE | DK | | 1 | 188 | 31 | 2747055 | 24 | 2744670 | 539222 | 465821 | 3 | 32.75 | 2x150bp | |
| 125351 | SAB | DK | | 5 | 5 | 31 | 2693854 | 22 | 2690171 | 1017201 | 554254 | 2 | 32.76 | 2x250bp | |
| 126179 | IE | DK | | 5 | 5 | 40 | 2761150 | 29 | 2757411 | 1021273 | 385875 | 2 | 32.73 | 2x250bp | |
| 126181 | IE | DK | | 45 | 45 | 28 | 2696273 | 21 | 2693272 | 1006273 | 605399 | 2 | 32.77 | 2x250bp | |
| 126673 | IE | DK | | 45 | 45 | 37 | 2757801 | 29 | 2754284 | 601904 | 512701 | 3 | 32.80 | 2x250bp | |
| 126691 | IE | DK | | 45 | 45 | 49 | 2784240 | 33 | 2778388 | 902116 | 672372 | 2 | 32.82 | 2x150bp | |
| 127097 | IE | DK | | 22 | 22 | 41 | 2717281 | 36 | 2715250 | 517060 | 174374 | 5 | 32.75 | 2x250bp | |
| 130639 | SAB | DK | | 15 | 15 | 29 | 2764607 | 22 | 2761552 | 657831 | 432813 | 3 | 32.77 | 2x250bp | |
| 130641 | SAB | DK | | 1 | 573 | 44 | 2742432 | 32 | 2737373 | 593144 | 184996 | 4 | 32.70 | 2x250bp | |
| 130867 | SAB | DK | | 45 | 45 | 29 | 2709878 | 21 | 2706720 | 527651 | 465333 | 3 | 32.79 | 2x250bp | |
| 130939 | SAB | DK | | 25 | 10001* | 19 | 2733192 | 17 | 2732310 | 978765 | 661228 | 2 | 32.68 | 2x250bp | *Unknown ST |

| | | | | | | | | | | | | | | | |
|---|---|---|---|---|---|---|---|---|---|---|---|---|---|---|---|
| 130987 | IE | DK | | 97 | 97 | 33 | 2732642 | 26 | 2729749 | 701777 | 280610 | 3 | 32.71 | 2x250bp | |
| 130989 | SAB | DK | | 45 | 45 | 31 | 2759297 | 24 | 2757003 | 529982 | 243275 | 4 | 32.77 | 2x150bp | |
| 131271 | SAB | DK | | 7 | 7 | 54 | 2789894 | 32 | 2782532 | 318623 | 198373 | 6 | 32.71 | 2x150bp | |
| 131973 | SAB | DK | | 30 | 30 | 71 | 2812027 | 56 | 2806735 | 430248 | 125349 | 7 | 32.73 | 2x150bp | |
| 131975 | SAB | DK | | 25 | 25 | 23 | 2717828 | 17 | 2715605 | 1067463 | 560725 | 2 | 32.66 | 2x150bp | |
| 132093 | SAB | DK | | 8 | 630 | 42 | 2792794 | 35 | 2789731 | 453678 | 253787 | 4 | 32.65 | 2x250bp | |
| 132105 | IE | DK | | 25 | 25 | 26 | 2713152 | 21 | 2711519 | 902619 | 694068 | 2 | 32.68 | 2x150bp | |
| 132281 | IE | DK | | 1 | 1 | 21 | 2750255 | 15 | 2747681 | 2288594 | 2288594 | 1 | 32.67 | 2x250bp | |
| 132647 | SAB | DK | | 1 | 1 | 27 | 2798391 | 17 | 2794270 | 1008734 | 879258 | 2 | 32.69 | 2x250bp | |
| 132731 | SAB | DK | | 5 | 5 | 26 | 2726731 | 20 | 2724334 | 779913 | 617397 | 2 | 32.77 | 2x250bp | |
| 132735 | IE | DK | | 30 | 30 | 40 | 2713864 | 33 | 2711308 | 471053 | 186002 | 5 | 32.74 | 2x150bp | |
| 132751 | SAB | DK | | 30 | 30 | 60 | 2825033 | 51 | 2821399 | 422223 | 148103 | 7 | 32.75 | 2x250bp | |
| 132855 | SAB | DK | | 45 | 45 | 31 | 2760989 | 22 | 2757311 | 601420 | 437177 | 3 | 32.83 | 2x250bp | |
| 132895 | IE | DK | | 50 | 50 | 54 | 2767874 | 42 | 2763315 | 435336 | 212335 | 5 | 32.72 | 2x250bp | |
| 133057 | SAB | DK | | 12 | 12 | 36 | 2739813 | 29 | 2737406 | 638875 | 330084 | 3 | 32.77 | 2x150bp | |
| 133271 | SAB | DK | | 30 | 30 | 86 | 2856333 | 68 | 2849359 | 393418 | 116073 | 7 | 32.73 | 2x150bp | |
| 133593 | IE | DK | | 15 | 15 | 47 | 2742106 | 30 | 2736419 | 441129 | 227995 | 5 | 32.69 | 2x150bp | |
| 133611 | SAB | DK | | 8 | 8 | 28 | 2731492 | 25 | 2730089 | 807686 | 477518 | 3 | 32.69 | 2x250bp | |
| 133621 | SAB | DK | | 30 | 30 | 48 | 2767412 | 40 | 2764542 | 488139 | 155107 | 6 | 32.69 | 2x150bp | |
| 134033 | SAB | DK | | 45 | 45 | 52 | 2745539 | 49 | 2744354 | 301646 | 134241 | 7 | 32.80 | 2x150bp | |
| 134333 | IE | DK | | 30 | 34 | 74 | 2835582 | 64 | 2831469 | 348958 | 171202 | 6 | 32.72 | 2x250bp | |
| 134361 | SAB | DK | | 5 | 5 | 28 | 2711667 | 23 | 2709575 | 1020148 | 734340 | 2 | 32.74 | 2x250bp | |
| 134363 | IE | DK | | 8 | 8 | 19 | 2727732 | 14 | 2725509 | 1163753 | 1010038 | 2 | 32.68 | 2x250bp | |
| 134631 | SAB | DK | | 45 | 45 | 18 | 2695208 | 15 | 2693819 | 991418 | 606690 | 2 | 32.78 | 2x250bp | |
| 134633 | SAB | DK | | 15 | 15 | 60 | 2652506 | 54 | 2650463 | 245508 | 104027 | 9 | 32.68 | 2x150bp | |
| 134741 | SAB | DK | | 20 | 10002* | 39 | 2722869 | 32 | 2720260 | 523574 | 192453 | 5 | 32.69 | 2x150bp | *Unknown ST |
| 134787 | SAB | DK | | 30 | 30 | 46 | 2825455 | 42 | 2823758 | 533904 | 170265 | 5 | 32.72 | 2x250bp | |
| 134789 | IE | DK | | 1 | 573 | 41 | 2758886 | 37 | 2757265 | 590613 | 356092 | 3 | 32.78 | 2x250bp | |
| 134967 | IE | DK | | 45 | 45 | 31 | 2725424 | 20 | 2720900 | 1023213 | 615426 | 2 | 32.83 | 2x250bp | |
| 134975 | SAB | DK | | 59 | 59 | 60 | 2721165 | 48 | 2716617 | 330719 | 117893 | 8 | 32.76 | 2x150bp | |
| 134977 | IE | DK | | 5 | 5 | 26 | 2718648 | 22 | 2717205 | 982390 | 399440 | 2 | 32.72 | 2x250bp | |
| 135213 | SAB | DK | | 15 | 15 | 48 | 2754954 | 35 | 2750253 | 697800 | 236829 | 4 | 32.75 | 2x150bp | |
| 135239 | SAB | DK | | 15 | 15 | 28 | 2665273 | 23 | 2663247 | 498113 | 243710 | 4 | 32.68 | 2x150bp | |
| 135251 | SAB | DK | | 22 | 22 | 59 | 2770188 | 48 | 2766096 | 338085 | 150818 | 7 | 32.73 | 2x150bp | |
| 135387 | SAB | DK | | 15 | 15 | 27 | 2694507 | 21 | 2692160 | 585869 | 367651 | 3 | 32.69 | 2x250bp | |
| 135429 | SAB | DK | | 45 | 45 | 25 | 2709517 | 19 | 2706919 | 1032070 | 468613 | 2 | 32.82 | 2x250bp | |
| 135557 | IE | DK | | 12 | 12 | 26 | 2661085 | 16 | 2656916 | 1282066 | 660779 | 2 | 32.73 | 2x250bp | |
| 135627 | IE | DK | | 15 | 15 | 25 | 2702711 | 19 | 2700164 | 610839 | 436705 | 3 | 32.78 | 2x250bp | |
| 135631 | SAB | DK | | 59 | 59 | 52 | 2677356 | 43 | 2674058 | 329256 | 132757 | 7 | 32.74 | 2x150bp | |
| 135633 | IE | DK | | 8 | 1181 | 28 | 2789251 | 20 | 2785762 | 1236504 | 579434 | 2 | 32.65 | 2x250bp | |
| 135687 | SAB | DK | | 45 | 45 | 23 | 2736469 | 23 | 2736469 | 1227498 | 601530 | 2 | 32.83 | 2x250bp | |
| 135689 | IE | DK | | 45 | 45 | 25 | 2734757 | 20 | 2732596 | 1020674 | 467176 | 2 | 32.79 | 2x250bp | |
| 135703 | IE | DK | | 1 | 1 | 29 | 2732106 | 21 | 2729287 | 443241 | 296540 | 4 | 32.69 | 2x150bp | |
| 135721 | SAB | DK | | 15 | 15 | 29 | 2741003 | 20 | 2737245 | 642377 | 347129 | 3 | 32.75 | 2x250bp | |
| 135775 | SAB | DK | | 8 | 8 | 45 | 2768191 | 32 | 2763713 | 674858 | 256897 | 3 | 32.63 | 2x150bp | |
| 135777 | SAB | DK | | 30 | 30 | 48 | 2865123 | 42 | 2862680 | 534998 | 181835 | 5 | 32.74 | 2x250bp | |
| 135865 | SAB | DK | | 30 | 30 | 67 | 2847027 | 54 | 2842238 | 368968 | 101351 | 9 | 32.72 | 2x150bp | |
| 135897 | SAB | DK | | 30 | 30 | 58 | 2831944 | 53 | 2829989 | 254751 | 134771 | 8 | 32.74 | 2x250bp | |
| 135901 | SAB | DK | | 45 | 45 | 28 | 2672850 | 15 | 2668404 | 992458 | 603635 | 2 | 32.77 | 2x150bp | |
| 135911 | SAB | DK | | 45 | 45 | 36 | 2727468 | 25 | 2723430 | 998869 | 282276 | 3 | 32.74 | 2x150bp | |
| 135933 | SAB | DK | | 30 | 30 | 57 | 2824356 | 51 | 2821964 | 276290 | 156095 | 8 | 32.72 | 2x250bp | |
| 135989 | SAB | DK | | 30 | 30 | 47 | 2848605 | 43 | 2847054 | 422296 | 151202 | 7 | 32.76 | 2x250bp | |
| 136011 | SAB | DK | | 59 | 59 | 52 | 2743902 | 37 | 2737870 | 304243 | 170383 | 6 | 32.75 | 2x250bp | |
| 136031 | SAB | DK | | 5 | 5 | 35 | 2824390 | 29 | 2821857 | 628084 | 389992 | 3 | 32.76 | 2x250bp | |
| 136107 | SAB | DK | Mice | 5 | 5 | 25 | 2723004 | 23 | 2722017 | 677353 | 450805 | 3 | 32.70 | 2x250bp | |
| 136123 | SAB | DK | Mice | 5 | 5 | 34 | 2727256 | 27 | 2724379 | 641867 | 290548 | 3 | 32.71 | 2x250bp | |
| 136357 | SAB | DK | | 15 | 15 | 29 | 2705637 | 20 | 2701915 | 586200 | 315192 | 4 | 32.74 | 2x250bp | |
| 136361 | SAB | DK | | 509 | 89 | 105 | 2787702 | 77 | 2777802 | 219585 | 80252 | 12 | 32.75 | 2x150bp | |
| 136383 | SAB | DK | | 7 | 7 | 61 | 2730129 | 53 | 2726997 | 279232 | 126183 | 7 | 32.70 | 2x150bp | |
| 136463 | SAB | DK | | 45 | 10003* | 22 | 2697587 | 19 | 2696479 | 765418 | 423063 | 3 | 32.76 | 2x150bp | *Unknown ST |
| 136555 | SAB | DK | | 15 | 15 | 38 | 2672875 | 25 | 2668296 | 611695 | 246552 | 4 | 32.68 | 2x150bp | |
| 136707 | SAB | DK | | 30 | 30 | 47 | 2787183 | 41 | 2785079 | 368905 | 155103 | 7 | 32.72 | 2x150bp | |
| 136709 | SAB | DK | | 30 | 30 | 48 | 2823950 | 41 | 2821175 | 279368 | 151221 | 7 | 32.72 | 2x250bp | |
| 136735 | SAB | DK | | 15 | 15 | 26 | 2723368 | 21 | 2721261 | 656208 | 319640 | 3 | 32.72 | 2x250bp | |
| 136889 | SAB | DK | | 30 | 30 | 61 | 2767189 | 49 | 2762798 | 392804 | 121889 | 7 | 32.69 | 2x150bp | |
| 136923 | SAB | DK | | 8 | 27 | 31 | 2715755 | 18 | 2711348 | 779942 | 358448 | 3 | 32.66 | 2x150bp | |
| 136925 | SAB | DK | | 45 | 45 | 42 | 2690594 | 39 | 2689469 | 320615 | 150469 | 7 | 32.78 | 2x150bp | |
| 136927 | IE | DK | | 45 | 45 | 31 | 2695926 | 23 | 2693184 | 698827 | 407737 | 3 | 32.79 | 2x150bp | |
| 136929 | SAB | DK | | 5 | 5 | 33 | 2719123 | 24 | 2715440 | 766148 | 379143 | 3 | 32.73 | 2x250bp | |
| 136941 | IE | DK | | 45 | 45 | 28 | 2719474 | 24 | 2717761 | 992814 | 607868 | 2 | 32.77 | 2x250bp | |
| 136943 | IE | DK | | 30 | 30 | 61 | 2789067 | 52 | 2785978 | 423999 | 113859 | 8 | 32.71 | 2x150bp | |
| 137019 | SAB | DK | | 1 | 1 | 16 | 2750631 | 11 | 2748491 | 1292201 | 1048102 | 2 | 32.67 | 2x250bp | |
| 137021 | IE | DK | | 45 | 45 | 29 | 2717070 | 23 | 2714472 | 974762 | 607178 | 2 | 32.78 | 2x250bp | |
| 137091 | SAB | DK | | 1 | 9 | 48 | 2791923 | 37 | 2787763 | 355661 | 177331 | 6 | 32.67 | 2x150bp | |
| 137101 | SAB | DK | | 7 | 7 | 28 | 2725296 | 21 | 2722459 | 707740 | 319504 | 3 | 32.74 | 2x250bp | |
| 137145 | SAB | DK | | 45 | 45 | 28 | 2724710 | 20 | 2721777 | 841229 | 606891 | 2 | 32.84 | 2x250bp | |
| 137217 | SAB | DK | | 1 | 1 | 34 | 2799861 | 27 | 2796803 | 949648 | 854597 | 2 | 32.77 | 2x250bp | |
| 137221 | IE | DK | | 30 | 30 | 45 | 2753433 | 38 | 2750671 | 417275 | 173605 | 5 | 32.72 | 2x250bp | |

| | | | | | | | | | | | | | | |
|---|---|---|---|---|---|---|---|---|---|---|---|---|---|---|
| 137233 | SAB | DK | 59 | 59 | 59 | 2715687 | 42 | 2708767 | 304234 | 157180 | 6 | 32.72 | 2x250bp | |
| 137307 | SAB | DK | 15 | 15 | 23 | 2678089 | 19 | 2676401 | 612897 | 434138 | 3 | 32.71 | 2x250bp | |
| 137309 | SAB | DK | 15 | 15 | 37 | 2719744 | 23 | 2714857 | 611965 | 434528 | 3 | 32.74 | 2x250bp | |
| 137381 | SAB | DK | 30 | 30 | 50 | 2746989 | 39 | 2743110 | 417322 | 157560 | 6 | 32.71 | 2x250bp | |
| 137601 | SAB | DK | 15 | 582 | 41 | 2722843 | 27 | 2718115 | 541048 | 193385 | 5 | 32.72 | 2x150bp | |
| 137603 | SAB | DK | 45 | 45 | 28 | 2708544 | 21 | 2706151 | 1045601 | 343321 | 2 | 32.82 | 2x150bp | |
| 137605 | IE | DK | 1 | 1 | 29 | 2717753 | 26 | 2716693 | 567780 | 251242 | 4 | 32.71 | 2x150bp | |
| 137607 | SAB | DK | 30 | 39 | 52 | 2756834 | 43 | 2753346 | 401547 | 143974 | 7 | 32.76 | 2x150bp | |
| 137609 | SAB | DK | 22 | 22 | 40 | 2719224 | 33 | 2716311 | 419554 | 174320 | 6 | 32.75 | 2x250bp | |
| 137611 | IE | DK | 5 | 5 | 25 | 2728812 | 21 | 2727219 | 1018001 | 480340 | 2 | 32.80 | 2x250bp | |
| 137613 | SAB | DK | 121 | 121 | 63 | 2815725 | 44 | 2808876 | 266868 | 99307 | 9 | 32.69 | 2x150bp | |
| 137615 | SAB | DK | 45 | 45 | 28 | 2710255 | 20 | 2707337 | 986836 | 403029 | 2 | 32.76 | 2x150bp | |
| 137617 | SAB | DK | 25 | 25 | 31 | 2736791 | 23 | 2734140 | 746785 | 346871 | 3 | 32.66 | 2x150bp | |
| 137619 | SAB | DK | 1 | 1 | 17 | 2751368 | 12 | 2749077 | 1078256 | 732518 | 2 | 32.67 | 2x250bp | |
| 137621 | SAB | DK | 97 | 97 | 32 | 2739487 | 25 | 2736686 | 794547 | 671620 | 2 | 32.71 | 2x250bp | |
| 137623 | SAB | DK | 15 | 15 | 25 | 2698830 | 20 | 2696806 | 656228 | 315352 | 3 | 32.76 | 2x250bp | |
| 137625 | SAB | DK | 1 | 81 | 19 | 2792078 | 15 | 2790376 | 1352642 | 713515 | 2 | 32.71 | 2x250bp | |
| 140005 | IE | DK | 15 | 15 | 26 | 2718330 | 22 | 2716554 | 657052 | 437432 | 3 | 32.73 | 2x250bp | |
| 140057 | IE | DK | 30 | 30 | 70 | 2809070 | 63 | 2806616 | 314534 | 95851 | 9 | 32.71 | 2x150bp | |
| 140101 | SAB | DK | 45 | 45 | 27 | 2698909 | 21 | 2696348 | 992811 | 600974 | 2 | 32.77 | 2x250bp | |
| 140135 | SAB | DK | 45 | 45 | 27 | 2731588 | 21 | 2729059 | 1013039 | 608095 | 2 | 32.85 | 2x250bp | |
| 140141 | SAB | DK | 45 | 10004* | 51 | 2772939 | 44 | 2770667 | 334415 | 126891 | 8 | 32.75 | 2x150bp | *Unknown ST |
| 140175 | SAB | DK | 30 | 30 | 67 | 2801070 | 55 | 2796598 | 284519 | 126479 | 8 | 32.76 | 2x150bp | |
| 140177 | IE | DK | 45 | 45 | 32 | 2760404 | 23 | 2757105 | 1039733 | 424847 | 2 | 32.84 | 2x150bp | |
| 140179 | SAB | DK | 5 | 5 | 32 | 2711648 | 22 | 2707498 | 857117 | 766350 | 2 | 32.74 | 2x250bp | |
| 140181 | SAB | DK | 59 | 87 | 59 | 2764139 | 51 | 2761126 | 310725 | 102694 | 8 | 32.70 | 2x150bp | |
| 140189 | SAB | DK | 15 | 15 | 29 | 2726466 | 22 | 2723468 | 612002 | 319373 | 3 | 32.75 | 2x250bp | |
| 140473 | SAB | DK | 30 | 30 | 58 | 2811351 | 48 | 2807753 | 489253 | 123590 | 7 | 32.73 | 2x150bp | |
| 140491 | SAB | DK | 45 | 45 | 26 | 2707038 | 24 | 2706232 | 925859 | 557975 | 2 | 32.78 | 2x250bp | |
| 140497 | SAB | DK | 8 | 72 | 31 | 2712865 | 26 | 2711079 | 720220 | 256867 | 3 | 32.68 | 2x150bp | |
| 140579 | SAB | DK | 45 | 45 | 34 | 2735497 | 28 | 2732915 | 926096 | 483921 | 2 | 32.86 | 2x250bp | |
| 140601 | IE | DK | 30 | 30 | 75 | 2811734 | 53 | 2804416 | 198742 | 110352 | 10 | 32.71 | 2x150bp | |
| 140733 | SAB | DK | 45 | 45 | 21 | 2703697 | 19 | 2702795 | 1230412 | 600624 | 2 | 32.79 | 2x250bp | |
| 140751 | IE | DK | 45 | 45 | 28 | 2737680 | 19 | 2734034 | 992417 | 608302 | 2 | 32.85 | 2x250bp | |
| 140755 | SAB | DK | 1 | 1 | 21 | 2800575 | 14 | 2797527 | 1239679 | 1003860 | 2 | 32.69 | 2x250bp | |
| 140829 | SAB | DK | 1 | 1 | 51 | 2800073 | 34 | 2792990 | 1029481 | 811019 | 2 | 32.78 | 2x250bp | |
| 140843 | SAB | DK | 1 | 1 | 17 | 2770753 | 11 | 2768274 | 1350999 | 1045747 | 2 | 32.73 | 2x250bp | |
| 140861 | SAB | DK | 8 | 8 | 33 | 2785277 | 21 | 2780832 | 701713 | 380464 | 3 | 32.69 | 2x250bp | |
| 140863 | SAB | DK | 123 | 123 | 55 | 2773338 | 37 | 2766880 | 432074 | 132318 | 5 | 32.75 | 2x150bp | |
| 140865 | SAB | DK | 30 | 34 | 59 | 2834263 | 45 | 2829245 | 369017 | 150906 | 6 | 32.70 | 2x150bp | |
| 140991 | IE | DK | 30 | 30 | 52 | 2796989 | 43 | 2793777 | 392716 | 131212 | 7 | 32.75 | 2x150bp | |
| 140995 | SAB | DK | 20 | 20 | 64 | 2784007 | 32 | 2772395 | 757382 | 311440 | 3 | 32.72 | 2x250bp | |
| 141103 | SAB | DK | 45 | 45 | 28 | 2712959 | 22 | 2710838 | 528930 | 370985 | 3 | 32.73 | 2x150bp | |
| 141163 | SAB | DK | 45 | 45 | 29 | 2704454 | 24 | 2702292 | 976350 | 607445 | 2 | 32.79 | 2x250bp | |
| 141169 | IE | DK | 1 | 1 | 31 | 2747020 | 21 | 2743507 | 554408 | 415925 | 3 | 32.66 | 2x150bp | |
| 141373 | IE | DK | 8 | 10005* | 38 | 2862394 | 32 | 2859897 | 1008606 | 881297 | 2 | 32.56 | 2x250bp | *Unknown ST |
| 141377 | SAB | DK | 45 | 45 | 35 | 2697280 | 22 | 2692843 | 990653 | 420076 | 2 | 32.79 | 2x150bp | |
| 141423 | IE | DK | 30 | 30 | 41 | 2824494 | 37 | 2822854 | 607528 | 175566 | 4 | 32.74 | 2x250bp | |
| 141431 | SAB | DK | 30 | 30 | 56 | 2763704 | 46 | 2760015 | 370812 | 150417 | 7 | 32.70 | 2x150bp | |
| 141475 | SAB | DK | 30 | 30 | 63 | 2805799 | 52 | 2801275 | 488408 | 175177 | 5 | 32.72 | 2x250bp | |
| 141521 | SAB | DK | 5 | 5 | 27 | 2726156 | 18 | 2722389 | 863829 | 474152 | 3 | 32.72 | 2x250bp | |
| 141711 | SAB | DK | 509 | 89 | 74 | 2760542 | 65 | 2756840 | 270507 | 103151 | 9 | 32.76 | 2x250bp | |
| 141729 | SAB | DK | 22 | 22 | 59 | 2754023 | 45 | 2748916 | 252093 | 133923 | 8 | 32.76 | 2x150bp | |
| 141739 | SAB | DK | 182 | 182 | 88 | 2755879 | 74 | 2750938 | 243274 | 94243 | 10 | 32.66 | 2x250bp | |
| 142059 | SAB | DK | 12 | 12 | 26 | 2725681 | 22 | 2724068 | 675577 | 620523 | 3 | 32.72 | 2x250bp | |
| 142077 | IE | DK | 45 | 45 | 40 | 2734753 | 28 | 2730623 | 533779 | 382072 | 3 | 32.82 | 2x150bp | |
| 142083 | SAB | DK | 1 | 1 | 17 | 2771920 | 10 | 2769075 | 1413927 | 1413927 | 1 | 32.73 | 2x250bp | |
| 142297 | SAB | DK | 22 | 22 | 48 | 2753264 | 36 | 2748308 | 512957 | 176658 | 5 | 32.78 | 2x250bp | |
| 142313 | SAB | DK | 45 | 45 | 23 | 2750266 | 20 | 2748960 | 1035930 | 627723 | 2 | 32.84 | 2x250bp | |
| 142321 | SAB | DK | 22 | 737 | 65 | 2800594 | 40 | 2792260 | 518487 | 180239 | 5 | 32.73 | 2x150bp | |
| 142469 | IE | DK | 30 | 34 | 42 | 2813984 | 37 | 2812079 | 545833 | 175653 | 4 | 32.73 | 2x250bp | |
| 142583 | IE | DK | 5 | 5 | 34 | 2699065 | 24 | 2694991 | 647088 | 334178 | 3 | 32.75 | 2x250bp | |
| 142705 | SAB | DK | 15 | 15 | 24 | 2722142 | 20 | 2720521 | 655252 | 436623 | 3 | 32.72 | 2x250bp | |
| 142777 | IE | DK | 1 | 1 | 21 | 2748886 | 12 | 2745161 | 1368495 | 1028476 | 2 | 32.67 | 2x250bp | |
| 142813 | SAB | DK | 45 | 45 | 24 | 2670158 | 17 | 2667748 | 1225761 | 467873 | 2 | 32.77 | 2x150bp | |
| 143139 | IE | DK | 8 | 8 | 51 | 2864353 | 38 | 2858988 | 807512 | 307082 | 3 | 32.55 | 2x250bp | |
| 143369 | SAB | DK | 15 | 15 | 38 | 2699338 | 29 | 2695642 | 546943 | 252267 | 4 | 32.70 | 2x250bp | |
| 143373 | SAB | DK | 30 | 39 | 66 | 2826295 | 54 | 2821863 | 292314 | 91666 | 10 | 32.80 | 2x250bp | |
| 144667 | SAB | DK | 5 | 5 | 27 | 2765235 | 22 | 2763103 | 1022711 | 702661 | 2 | 32.74 | 2x250bp | |
| 144669 | SAB | DK | 45 | 45 | 24 | 2700549 | 20 | 2698757 | 608235 | 424242 | 3 | 32.81 | 2x250bp | |
| 144671 | IE | DK | 8 | 8 | 61 | 2795497 | 41 | 2788432 | 361724 | 239030 | 5 | 32.64 | 2x150bp | |
| 144673 | SAB | DK | 1 | 1 | 24 | 2789182 | 21 | 2787763 | 683001 | 380725 | 3 | 32.69 | 2x250bp | |
| 144675 | SAB | DK | 30 | 10006* | 215 | 2872728 | 192 | 2864081 | 116250 | 27326 | 31 | 32.72 | 2x150bp | *Unknown ST |
| 144677 | SAB | DK | 45 | 45 | 29 | 2696968 | 21 | 2694283 | 522980 | 423966 | 3 | 32.76 | 2x150bp | |
| BPH2702 | SAB | AUS/NZ | 15 | 15 | 45 | 2821612 | 39 | 2818890 | 437331 | 246733 | 5 | 32.78 | 2x250bp | |
| BPH2703 | SAB | AUS/NZ | 8 | 8 | 41 | 2774473 | 39 | 2773579 | 434227 | 212223 | 5 | 32.69 | 2x250bp | |
| BPH2704 | SAB | AUS/NZ | 93 | 93 | 30 | 2809823 | 27 | 2808621 | 820902 | 645629 | 2 | 32.68 | 2x250bp | |
| BPH2706 | SAB | AUS/NZ | 45 | 508 | 27 | 2747061 | 26 | 2746573 | 906038 | 540353 | 2 | 32.85 | 2x250bp | |

| | | | | | | | | | | | | | | |
|---|---|---|---|---|---|---|---|---|---|---|---|---|---|---|
| BPH2707 | SAB | AUS/NZ | 8 | 8 | 28 | 2747734 | 26 | 2746982 | 882277 | 511542 | 2 | 32.67 | 2x250bp | |
| BPH2708 | SAB | AUS/NZ | 97 | 97 | 25 | 2724939 | 24 | 2724482 | 508642 | 225683 | 4 | 32.71 | 2x250bp | |
| BPH2711 | SAB | AUS/NZ | 22 | 22 | 41 | 2750005 | 40 | 2749603 | 540937 | 164562 | 5 | 32.77 | 2x250bp | |
| BPH2713 | SAB | AUS/NZ | 45 | 45 | 76 | 2730634 | 71 | 2728774 | 146900 | 88467 | 12 | 32.82 | 2x250bp | |
| BPH2716 | SAB | AUS/NZ | 5 | 5 | 27 | 2829641 | 25 | 2828654 | 564158 | 243975 | 4 | 32.75 | 2x250bp | |
| BPH2718 | SAB | AUS/NZ | 8 | 239 | 68 | 3026645 | 60 | 3023596 | 299881 | 145058 | 7 | 32.64 | 2x250bp | |
| BPH2719 | SAB | AUS/NZ | 5 | 5 | 26 | 2736660 | 20 | 2733926 | 692499 | 379699 | 3 | 32.79 | 2x250bp | |
| BPH2720 | SAB | AUS/NZ | 22 | 22 | 46 | 2816057 | 46 | 2816057 | 286750 | 142962 | 8 | 32.71 | 2x250bp | |
| BPH2722 | SAB | AUS/NZ | 1 | 188 | 32 | 2737777 | 31 | 2737307 | 536466 | 188150 | 5 | 32.71 | 2x250bp | |
| BPH2723 | SAB | AUS/NZ | 8 | 239 | 76 | 2943153 | 66 | 2939227 | 518170 | 109384 | 7 | 32.69 | 2x250bp | |
| BPH2724 | SAB | AUS/NZ | 30 | 30 | 82 | 2764630 | 76 | 2762284 | 221962 | 99410 | 9 | 32.73 | 2x250bp | |
| BPH2725 | SAB | AUS/NZ | 45 | 45 | 22 | 2719520 | 20 | 2718576 | 905547 | 333908 | 3 | 32.80 | 2x250bp | |
| BPH2726 | SAB | AUS/NZ | 8 | 239 | 60 | 2937987 | 56 | 2936211 | 321398 | 155660 | 7 | 32.67 | 2x250bp | |
| BPH2728 | SAB | AUS/NZ | 45 | 45 | 45 | 2809272 | 39 | 2806683 | 437329 | 199177 | 5 | 32.71 | 2x250bp | |
| BPH2731 | SAB | AUS/NZ | 121 | 121 | 57 | 2752971 | 52 | 2751097 | 352985 | 147286 | 6 | 32.74 | 2x250bp | |
| BPH2732 | SAB | AUS/NZ | 45 | 45 | 45 | 2749050 | 39 | 2746756 | 299836 | 164605 | 6 | 32.80 | 2x250bp | |
| BPH2735 | SAB | AUS/NZ | 22 | 22 | 72 | 2823948 | 59 | 2818915 | 330638 | 114806 | 7 | 32.71 | 2x250bp | |
| BPH2736 | SAB | AUS/NZ | 45 | 45 | 61 | 2749593 | 55 | 2747233 | 243262 | 106833 | 9 | 32.78 | 2x250bp | |
| BPH2737 | SAB | AUS/NZ | 15 | 10008* | 87 | 2805914 | 75 | 2801231 | 192897 | 72171 | 12 | 32.79 | 2x250bp | *Unknown ST |
| BPH2739 | IE | AUS/NZ | 20 | 20 | 34 | 2749288 | 30 | 2747573 | 423153 | 338579 | 4 | 32.69 | 2x250bp | |
| BPH2741 | SAB | AUS/NZ | 1 | 109 | 57 | 2841726 | 50 | 2838935 | 315877 | 107774 | 9 | 32.72 | 2x250bp | |
| BPH2742 | SAB | AUS/NZ | 22 | 22 | 144 | 2817348 | 134 | 2813049 | 118461 | 43289 | 20 | 32.73 | 2x250bp | |
| BPH2743 | SAB | AUS/NZ | 5 | 5 | 40 | 2736088 | 33 | 2733365 | 288048 | 148708 | 7 | 32.69 | 2x250bp | |
| BPH2744 | SAB | AUS/NZ | 8 | 3376 | 44 | 2822986 | 39 | 2821016 | 694895 | 384455 | 3 | 32.64 | 2x250bp | |
| BPH2745 | IE | AUS/NZ | 8 | 72 | 51 | 2718296 | 45 | 2715923 | 202399 | 99828 | 9 | 32.71 | 2x250bp | |
| BPH2746 | SAB | AUS/NZ | 8 | 8 | 45 | 2743126 | 38 | 2740412 | 326395 | 126593 | 6 | 32.67 | 2x250bp | |
| BPH2747 | SAB | AUS/NZ | 8 | 239 | 75 | 3004480 | 68 | 3001707 | 291562 | 142844 | 7 | 32.67 | 2x250bp | |
| BPH2749 | SAB | AUS/NZ | 20 | 20 | 52 | 2767555 | 38 | 2762255 | 353817 | 135631 | 6 | 32.67 | 2x250bp | |
| BPH2750 | SAB | AUS/NZ | 30 | 39 | 64 | 2774995 | 60 | 2773302 | 244940 | 111390 | 10 | 32.79 | 2x250bp | |
| BPH2752 | SAB | AUS/NZ | 30 | 30 | 118 | 2784658 | 117 | 2784300 | 196623 | 46107 | 18 | 32.75 | 2x250bp | |
| BPH2753 | SAB | AUS/NZ | 121 | 121 | 52 | 2801562 | 50 | 2800669 | 446072 | 143813 | 6 | 32.73 | 2x250bp | |
| BPH2754 | SAB | AUS/NZ | 5 | 5 | 57 | 2729342 | 50 | 2726461 | 249571 | 117439 | 8 | 32.72 | 2x250bp | |
| BPH2756 | SAB | AUS/NZ | 1 | 567 | 25 | 2838988 | 21 | 2837343 | 433822 | 322193 | 4 | 32.72 | 2x250bp | |
| BPH2757 | SAB | AUS/NZ | 5 | 5 | 37 | 2822624 | 34 | 2821181 | 710759 | 344651 | 3 | 32.75 | 2x250bp | |
| BPH2758 | SAB | AUS/NZ | 5 | 5 | 61 | 2794025 | 54 | 2791258 | 321322 | 115537 | 9 | 32.68 | 2x250bp | |
| BPH2759 | SAB | AUS/NZ | 30 | 30 | 47 | 2796147 | 41 | 2793940 | 394914 | 150541 | 6 | 32.74 | 2x250bp | |
| BPH2760 | IE | AUS/NZ | 1 | 1 | 26 | 2732251 | 24 | 2731383 | 557897 | 454184 | 3 | 32.66 | 2x250bp | |
| BPH2761 | SAB | AUS/NZ | 30 | 30 | 145 | 2796045 | 138 | 2793589 | 125816 | 42996 | 19 | 32.73 | 2x250bp | |
| BPH2763 | SAB | AUS/NZ | 78 | 78 | 38 | 2807167 | 32 | 2804735 | 710243 | 421148 | 3 | 32.72 | 2x250bp | |
| BPH2765 | SAB | AUS/NZ | 15 | 15 | 66 | 2736280 | 48 | 2728794 | 275437 | 113546 | 7 | 32.70 | 2x250bp | |
| BPH2767 | SAB | AUS/NZ | 5 | 3628 | 334 | 3010459 | 257 | 2975101 | 448119 | 131649 | 6 | 32.71 | 2x250bp | |
| BPH2768 | SAB | AUS/NZ | 30 | 30 | 45 | 2822869 | 41 | 2821419 | 414768 | 175308 | 5 | 32.72 | 2x150bp | |
| BPH2769 | SAB | AUS/NZ | 1 | 188 | 14 | 2738365 | 13 | 2738062 | 1168200 | 439195 | 2 | 32.75 | 2x250bp | |
| BPH2770 | SAB | AUS/NZ | 45 | 45 | 86 | 2842128 | 64 | 2833051 | 307193 | 103969 | 8 | 32.69 | 2x250bp | |
| BPH2773 | SAB | AUS/NZ | 1 | 188 | 24 | 2793911 | 19 | 2792005 | 709190 | 565402 | 3 | 32.70 | 2x150bp | |
| BPH2774 | SAB | AUS/NZ | 22 | 22 | 42 | 2748664 | 36 | 2746417 | 406581 | 174182 | 5 | 32.71 | 2x150bp | |
| BPH2775 | SAB | AUS/NZ | 8 | 8 | 21 | 2752463 | 15 | 2750177 | 1179014 | 544381 | 2 | 32.66 | 2x150bp | |
| BPH2776 | SAB | AUS/NZ | 30 | 30 | 67 | 2823312 | 60 | 2820693 | 422107 | 162594 | 5 | 32.62 | 2x150bp | |
| BPH2777 | SAB | AUS/NZ | 30 | 30 | 65 | 2860689 | 60 | 2858938 | 422051 | 109919 | 7 | 32.74 | 2x150bp | |
| BPH2781 | SAB | AUS/NZ | 8 | 239 | 65 | 3049741 | 59 | 3047386 | 564157 | 142921 | 5 | 32.67 | 2x150bp | |
| BPH2782 | SAB | AUS/NZ | 123 | 10009* | 78 | 2810327 | 57 | 2801730 | 184536 | 99322 | 11 | 32.71 | 2x250bp | *Unknown ST |
| BPH2783 | SAB | AUS/NZ | 22 | 22 | 108 | 2812068 | 95 | 2806588 | 244364 | 62449 | 13 | 32.69 | 2x250bp | |
| BPH2784 | SAB | AUS/NZ | 78 | 88 | 34 | 2774871 | 26 | 2771649 | 927506 | 296934 | 3 | 32.77 | 2x150bp | |
| BPH2785 | SAB | AUS/NZ | 22 | 22 | 53 | 2805610 | 43 | 2801363 | 407043 | 126265 | 7 | 32.71 | 2x250bp | |
| BPH2786 | SAB | AUS/NZ | 5 | 5 | 67 | 2762154 | 36 | 2749848 | 577365 | 176853 | 5 | 32.68 | 2x250bp | |
| BPH2788 | SAB | AUS/NZ | 15 | 15 | 27 | 2709326 | 23 | 2707880 | 622876 | 272924 | 4 | 32.75 | 2x150bp | |
| BPH2789 | SAB | AUS/NZ | 1 | 109 | 30 | 2788644 | 23 | 2786108 | 855454 | 288060 | 3 | 32.71 | 2x150bp | |
| BPH2790 | SAB | AUS/NZ | 22 | 22 | 45 | 2789916 | 34 | 2785544 | 348318 | 174035 | 6 | 32.72 | 2x150bp | |
| BPH2792 | SAB | AUS/NZ | 45 | 45 | 57 | 2838169 | 52 | 2836369 | 497024 | 126656 | 6 | 32.72 | 2x150bp | |
| BPH2794 | SAB | AUS/NZ | 5 | 5 | 36 | 2819998 | 31 | 2818179 | 759370 | 243758 | 3 | 32.72 | 2x150bp | |
| BPH2795 | SAB | AUS/NZ | 291 | 291 | 47 | 2773116 | 43 | 2771362 | 368812 | 133038 | 8 | 32.85 | 2x250bp | |
| BPH2796 | SAB | AUS/NZ | 97 | 97 | 52 | 2742238 | 51 | 2741781 | 295844 | 119830 | 8 | 32.71 | 2x300bp | |
| BPH2797 | SAB | AUS/NZ | 12 | 12 | 19 | 2696536 | 15 | 2695077 | 972721 | 956210 | 2 | 32.72 | 2x150bp | |
| BPH2798 | SAB | AUS/NZ | 8 | 239 | 49 | 2996982 | 45 | 2995368 | 544516 | 169319 | 5 | 32.64 | 2x150bp | |
| BPH2801 | SAB | AUS/NZ | 59 | 59 | 41 | 2727769 | 36 | 2725652 | 453194 | 168081 | 6 | 32.74 | 2x150bp | |
| BPH2805 | SAB | AUS/NZ | 30 | 39 | 41 | 2745268 | 37 | 2743769 | 340220 | 140623 | 7 | 32.75 | 2x150bp | |
| BPH2806 | SAB | AUS/NZ | 45 | 45 | 25 | 2713477 | 21 | 2712096 | 982699 | 295892 | 3 | 32.77 | 2x150bp | |
| BPH2807 | SAB | AUS/NZ | 22 | 22 | 43 | 2784787 | 37 | 2782526 | 309815 | 146469 | 7 | 32.73 | 2x150bp | |
| BPH2809 | SAB | AUS/NZ | 5 | 10010* | 31 | 2732739 | 25 | 2730565 | 456382 | 215591 | 5 | 32.70 | 2x150bp | *Unknown ST |
| BPH2810 | SAB | AUS/NZ | 45 | 508 | 25 | 2783669 | 21 | 2782292 | 1040923 | 480256 | 2 | 32.78 | 2x150bp | |
| BPH2811 | SAB | AUS/NZ | 15 | 15 | 26 | 2701460 | 22 | 2699873 | 606632 | 272569 | 4 | 32.76 | 2x150bp | |
| BPH2812 | SAB | AUS/NZ | 1 | 1 | 25 | 2854249 | 23 | 2853455 | 632842 | 345698 | 3 | 32.71 | 2x300bp | |
| BPH2817 | SAB | AUS/NZ | 1 | 188 | 25 | 2746929 | 18 | 2744440 | 704954 | 417947 | 3 | 32.66 | 2x150bp | |
| BPH2818 | SAB | AUS/NZ | 5 | 5 | 38 | 2754504 | 38 | 2754504 | 288128 | 170389 | 6 | 32.76 | 2x300bp | |
| BPH2819 | SAB | AUS/NZ | 5 | 5 | 26 | 2733009 | 21 | 2731190 | 551276 | 243814 | 4 | 32.70 | 2x150bp | |
| BPH2821 | IE | AUS/NZ | 12 | 12 | 20 | 2766384 | 16 | 2764877 | 1015321 | 653694 | 2 | 32.74 | 2x150bp | |
| BPH2822 | SAB | AUS/NZ | 1 | 1 | 17 | 2767268 | 12 | 2765343 | 1411013 | 1411013 | 1 | 32.72 | 2x150bp | |
| BPH2823 | SAB | AUS/NZ | 78 | 78 | 50 | 2805080 | 46 | 2803379 | 408889 | 130650 | 7 | 32.73 | 2x300bp | |

| | | | | | | | | | | | | | | |
|---|---|---|---|---|---|---|---|---|---|---|---|---|---|---|
| BPH2824 | SAB | AUS/NZ | 8 | 239 | 69 | 3054525 | 62 | 3051718 | 564160 | 155503 | 5 | 32.67 | 2x150bp | |
| BPH2827 | SAB | AUS/NZ | 8 | 239 | 113 | 2868092 | 112 | 2867601 | 162924 | 53762 | 16 | 32.78 | 2x300bp | |
| BPH2828 | SAB | AUS/NZ | 22 | 22 | 47 | 2787873 | 40 | 2785225 | 313042 | 174072 | 6 | 32.74 | 2x150bp | |
| BPH2829 | SAB | AUS/NZ | 8 | 8 | 36 | 2755847 | 26 | 2752275 | 698587 | 277270 | 3 | 32.66 | 2x300bp | |
| BPH2830 | SAB | AUS/NZ | 78 | 78 | 94 | 2757940 | 82 | 2752988 | 125189 | 75807 | 14 | 32.77 | 2x250bp | |
| BPH2831 | SAB | AUS/NZ | 8 | 8 | 46 | 2840496 | 41 | 2838370 | 407575 | 189590 | 6 | 32.71 | 2x300bp | |
| BPH2834 | SAB | AUS/NZ | 30 | 10011* | 67 | 2813271 | 60 | 2810704 | 331472 | 150309 | 6 | 32.70 | 2x150bp | *Unknown ST |
| BPH2835 | IE | AUS/NZ | 8 | 72 | 29 | 2695997 | 20 | 2692747 | 760521 | 361294 | 3 | 32.72 | 2x150bp | |
| BPH2836 | SAB | AUS/NZ | 8 | 8 | 51 | 2733299 | 44 | 2730673 | 366400 | 156491 | 6 | 32.68 | 2x300bp | |
| BPH2838 | SAB | AUS/NZ | 12 | 12 | 82 | 2699729 | 82 | 2699729 | 240101 | 69147 | 12 | 32.74 | 2x300bp | |
| BPH2839 | SAB | AUS/NZ | 1 | 1 | 80 | 2721601 | 70 | 2717537 | 194850 | 74607 | 12 | 32.72 | 2x250bp | |
| BPH2840 | SAB | AUS/NZ | 8 | 8 | 29 | 2824814 | 25 | 2823002 | 701154 | 370554 | 3 | 32.64 | 2x250bp | |
| BPH2843 | IE | AUS/NZ | 22 | 22 | 42 | 2814233 | 39 | 2813030 | 349609 | 174387 | 6 | 32.69 | 2x300bp | |
| BPH2845 | SAB | AUS/NZ | 1 | 1 | 31 | 2795288 | 28 | 2793947 | 532013 | 271096 | 4 | 32.62 | 2x300bp | |
| BPH2846 | SAB | AUS/NZ | 12 | 10012* | 33 | 2814306 | 32 | 2813807 | 974877 | 685842 | 2 | 32.75 | 2x300bp | *Unknown ST |
| BPH2847 | IE | AUS/NZ | 22 | 22 | 53 | 2809867 | 43 | 2806173 | 252340 | 146469 | 7 | 32.68 | 2x150bp | |
| BPH2848 | IE | AUS/NZ | 1 | 1 | 24 | 2762101 | 20 | 2760551 | 482499 | 315233 | 4 | 32.68 | 2x300bp | |
| BPH2849 | SAB | AUS/NZ | 5 | 5 | 88 | 2736963 | 85 | 2735846 | 241571 | 79801 | 11 | 32.70 | 2x250bp | |
| BPH2851 | SAB | AUS/NZ | 22 | 10013* | 41 | 2795094 | 36 | 2792964 | 348417 | 174275 | 6 | 32.73 | 2x300bp | *Unknown ST |
| BPH2852 | SAB | AUS/NZ | 101 | 101 | 14 | 2721691 | 12 | 2720880 | 756085 | 686746 | 2 | 32.73 | 2x300bp | |
| BPH2854 | IE | AUS/NZ | 8 | 8 | 28 | 2797045 | 19 | 2793645 | 988136 | 312650 | 3 | 32.60 | 2x150bp | |
| BPH2855 | SAB | AUS/NZ | 15 | 15 | 29 | 2721798 | 23 | 2719127 | 567367 | 378173 | 3 | 32.70 | 2x300bp | |
| BPH2856 | SAB | AUS/NZ | 22 | 22 | 40 | 2785702 | 36 | 2784094 | 311569 | 178302 | 6 | 32.78 | 2x300bp | |
| BPH2858 | SAB | AUS/NZ | 22 | 22 | 40 | 2809164 | 35 | 2807137 | 387321 | 146626 | 6 | 32.71 | 2x300bp | |
| BPH2859 | SAB | AUS/NZ | 93 | 93 | 36 | 2812586 | 34 | 2811814 | 646195 | 168357 | 4 | 32.68 | 2x300bp | |
| BPH2860 | SAB | AUS/NZ | 78 | 88 | 31 | 2769324 | 26 | 2767305 | 720484 | 299729 | 3 | 32.73 | 2x300bp | |
| BPH2861 | SAB | AUS/NZ | 30 | 39 | 37 | 2723628 | 31 | 2721262 | 451779 | 162671 | 6 | 32.74 | 2x150bp | |
| BPH2863 | SAB | AUS/NZ | 30 | 3381 | 65 | 2804214 | 57 | 2800962 | 370487 | 98501 | 8 | 32.71 | 2x250bp | |
| BPH2866 | SAB | AUS/NZ | 15 | 15 | 36 | 2775007 | 32 | 2773227 | 473941 | 168362 | 6 | 32.74 | 2x300bp | |
| BPH2868 | SAB | AUS/NZ | 93 | 93 | 30 | 2781710 | 21 | 2778356 | 620994 | 346707 | 3 | 32.68 | 2x150bp | |
| BPH2869 | SAB | AUS/NZ | 8 | 239 | 86 | 2914855 | 79 | 2912038 | 324809 | 108878 | 9 | 32.78 | 2x300bp | |
| BPH2870 | SAB | AUS/NZ | 78 | 78 | 21 | 2772621 | 20 | 2772290 | 839983 | 751059 | 2 | 32.75 | 2x300bp | |
| BPH2871 | SAB | AUS/NZ | 78 | 88 | 19 | 2715343 | 14 | 2713512 | 975563 | 459317 | 2 | 32.75 | 2x150bp | |
| BPH2872 | IE | AUS/NZ | 5 | 10014* | 32 | 2800222 | 27 | 2798302 | 667652 | 378714 | 3 | 32.72 | 2x150bp | *Unknown ST |
| BPH2874 | SAB | AUS/NZ | 7 | 7 | 23 | 2786161 | 17 | 2783813 | 637267 | 302591 | 4 | 32.72 | 2x150bp | |
| BPH2875 | SAB | AUS/NZ | 30 | 34 | 68 | 2856929 | 62 | 2854596 | 332347 | 114866 | 7 | 32.73 | 2x150bp | |
| BPH2876 | SAB | AUS/NZ | 78 | 78 | 29 | 2788311 | 27 | 2787355 | 717324 | 329097 | 3 | 32.73 | 2x300bp | |
| BPH2878 | SAB | AUS/NZ | 30 | 30 | 57 | 2813500 | 57 | 2813500 | 281233 | 138586 | 7 | 32.73 | 2x300bp | |
| BPH2879 | SAB | AUS/NZ | 22 | 22 | 39 | 2795868 | 37 | 2795067 | 349393 | 174251 | 6 | 32.73 | 2x300bp | |
| BPH2880 | SAB | AUS/NZ | 1 | 188 | 21 | 2769470 | 19 | 2768651 | 547021 | 405087 | 3 | 32.73 | 2x250bp | |
| BPH2881 | SAB | AUS/NZ | 5 | 1756 | 41 | 2821933 | 34 | 2819152 | 425717 | 170447 | 6 | 32.70 | 2x250bp | |
| BPH2883 | SAB | AUS/NZ | 5 | 5 | 48 | 2739167 | 41 | 2736312 | 536189 | 181956 | 5 | 32.70 | 2x250bp | |
| BPH2885 | SAB | AUS/NZ | 5 | 5 | 28 | 2751752 | 25 | 2750486 | 697501 | 243901 | 4 | 32.78 | 2x300bp | |
| BPH2886 | SAB | AUS/NZ | 5 | 5 | 31 | 2741840 | 28 | 2740574 | 523564 | 394218 | 3 | 32.72 | 2x300bp | |
| BPH2887 | SAB | AUS/NZ | 45 | 45 | 36 | 2902537 | 32 | 2901114 | 776739 | 354716 | 3 | 32.70 | 2x150bp | |
| BPH2889 | SAB | AUS/NZ | 1 | 9 | 31 | 2836350 | 22 | 2832890 | 665732 | 349887 | 3 | 32.72 | 2x150bp | |
| BPH2890 | SAB | AUS/NZ | 78 | 78 | 27 | 2765449 | 14 | 2760171 | 996335 | 479734 | 2 | 32.67 | 2x150bp | |
| BPH2892 | SAB | AUS/NZ | 1 | 109 | 35 | 2803238 | 24 | 2799348 | 714632 | 270365 | 3 | 32.69 | 2x150bp | |
| BPH2893 | SAB | AUS/NZ | 30 | 30 | 224 | 2798950 | 219 | 2796988 | 111623 | 27767 | 31 | 32.80 | 2x300bp | |
| BPH2894 | SAB | AUS/NZ | 121 | 121 | 45 | 2800039 | 38 | 2797361 | 353300 | 119137 | 7 | 32.72 | 2x150bp | |
| BPH2895 | SAB | AUS/NZ | 509 | 509 | 71 | 2726262 | 58 | 2721220 | 269012 | 95969 | 10 | 32.77 | 2x150bp | |
| BPH2897 | SAB | AUS/NZ | 30 | 34 | 55 | 2810645 | 42 | 2806085 | 342943 | 136187 | 8 | 32.72 | 2x150bp | |
| BPH2900 | SAB | AUS/NZ | 22 | 22 | 39 | 2786886 | 33 | 2784750 | 406824 | 173956 | 5 | 32.72 | 2x150bp | |
| BPH2901 | SAB | AUS/NZ | 5 | 5 | 37 | 2766118 | 33 | 2764463 | 397773 | 294402 | 5 | 32.74 | 2x150bp | |
| BPH2902 | SAB | AUS/NZ | 45 | 45 | 33 | 2833623 | 26 | 2830921 | 554065 | 285558 | 4 | 32.73 | 2x150bp | |
| BPH2905 | SAB | AUS/NZ | 101 | 10015* | 47 | 2735821 | 47 | 2735821 | 312571 | 112354 | 7 | 32.72 | 2x250bp | *Unknown ST |
| BPH2906 | SAB | AUS/NZ | 30 | 30 | 47 | 2817317 | 42 | 2815369 | 477498 | 175579 | 4 | 32.76 | 2x300bp | |
| BPH2908 | SAB | AUS/NZ | 15 | 15 | 31 | 2724764 | 24 | 2721805 | 368055 | 243763 | 5 | 32.71 | 2x300bp | |
| BPH2909 | SAB | AUS/NZ | 45 | 10016* | 23 | 2754687 | 20 | 2753245 | 1033689 | 599507 | 2 | 32.80 | 2x300bp | *Unknown ST |
| BPH2910 | SAB | AUS/NZ | 97 | 953 | 51 | 2744820 | 48 | 2743422 | 390772 | 121213 | 7 | 32.71 | 2x250bp | |
| BPH2911 | SAB | AUS/NZ | 93 | 93 | 27 | 2807926 | 27 | 2807926 | 560482 | 209223 | 4 | 32.68 | 2x300bp | |
| BPH2912 | IE | AUS/NZ | 1 | 1 | 15 | 2750249 | 10 | 2748324 | 1350946 | 751276 | 2 | 32.67 | 2x150bp | |
| BPH2913 | SAB | AUS/NZ | 5 | 5 | 37 | 2742972 | 32 | 2741090 | 433712 | 307247 | 4 | 32.67 | 2x150bp | |
| BPH2914 | SAB | AUS/NZ | 5 | 5 | 24 | 2735926 | 21 | 2734560 | 855690 | 380373 | 3 | 32.69 | 2x300bp | |
| BPH2915 | SAB | AUS/NZ | 30 | 30 | 45 | 2820172 | 43 | 2819369 | 331525 | 171450 | 6 | 32.73 | 2x300bp | |
| BPH2916 | SAB | AUS/NZ | 15 | 15 | 36 | 2722667 | 30 | 2720294 | 330423 | 193757 | 6 | 32.72 | 2x300bp | |
| BPH2917 | IE | AUS/NZ | 45 | 46 | 18 | 2712741 | 16 | 2711818 | 1030222 | 544828 | 2 | 32.85 | 2x300bp | |
| BPH2918 | SAB | AUS/NZ | 78 | 78 | 30 | 2787950 | 23 | 2784938 | 733300 | 687678 | 2 | 32.73 | 2x300bp | |
| BPH2920 | SAB | AUS/NZ | 22 | 22 | 43 | 2791522 | 36 | 2788939 | 406682 | 174036 | 5 | 32.73 | 2x150bp | |
| BPH2921 | SAB | AUS/NZ | 78 | 88 | 30 | 2751656 | 22 | 2748317 | 730208 | 298272 | 3 | 32.72 | 2x150bp | |
| BPH2922 | IE | AUS/NZ | 1 | 1 | 16 | 2716833 | 11 | 2714984 | 1319163 | 747763 | 2 | 32.67 | 2x150bp | |
| BPH2927 | SAB | AUS/NZ | 45 | 45 | 31 | 2792583 | 24 | 2789478 | 437503 | 348095 | 4 | 32.70 | 2x300bp | |
| BPH2928 | IE | AUS/NZ | 101 | 101 | 23 | 2741880 | 18 | 2740171 | 972349 | 311729 | 3 | 32.69 | 2x150bp | |
| BPH2929 | SAB | AUS/NZ | 30 | 30 | 40 | 2787287 | 31 | 2783724 | 543474 | 239194 | 4 | 32.71 | 2x150bp | |
| BPH2930 | SAB | AUS/NZ | 78 | 88 | 21 | 2715916 | 15 | 2713543 | 915823 | 531229 | 2 | 32.69 | 2x150bp | |
| BPH2932 | SAB | AUS/NZ | 8 | 8 | 31 | 2810569 | 27 | 2808786 | 1342790 | 332598 | 2 | 32.71 | 2x300bp | |
| BPH2933 | SAB | AUS/NZ | 5 | 5 | 71 | 2775260 | 69 | 2774273 | 194873 | 78894 | 13 | 32.72 | 2x300bp | |
| BPH2934 | SAB | AUS/NZ | 291 | 291 | 33 | 2729802 | 26 | 2727334 | 450651 | 212242 | 4 | 32.80 | 2x150bp | |

| | | | | | | | | | | | | | | |
|---|---|---|---|---|---|---|---|---|---|---|---|---|---|---|
| BPH2935 | IE | AUS/NZ | 30 | 3383 | 56 | 2832167 | 55 | 2831846 | 324243 | 114114 | 8 | 32.72 | 2x300bp | |
| BPH2936 | SAB | AUS/NZ | 5 | 5 | 25 | 2820330 | 24 | 2819842 | 776543 | 322314 | 3 | 32.69 | 2x300bp | |
| BPH2937 | SAB | AUS/NZ | 8 | 8 | 36 | 2783264 | 27 | 2779808 | 544493 | 406689 | 3 | 32.67 | 2x150bp | |
| BPH2938 | SAB | AUS/NZ | 30 | 30 | 56 | 2809376 | 53 | 2808154 | 392790 | 151156 | 6 | 32.73 | 2x300bp | |
| BPH2939 | SAB | AUS/NZ | 5 | 5 | 33 | 2738104 | 29 | 2736432 | 856766 | 342898 | 3 | 32.71 | 2x300bp | |
| BPH2940 | SAB | AUS/NZ | 78 | 88 | 26 | 2726152 | 22 | 2724327 | 410268 | 283059 | 5 | 32.70 | 2x300bp | |
| BPH2941 | SAB | AUS/NZ | 22 | 22 | 44 | 2792172 | 41 | 2790969 | 538895 | 155798 | 5 | 32.73 | 2x300bp | |
| BPH2944 | SAB | AUS/NZ | 22 | 60 | 53 | 2725666 | 49 | 2724119 | 245550 | 128455 | 8 | 32.81 | 2x300bp | |
| BPH2945 | SAB | AUS/NZ | 30 | 30 | 53 | 2804013 | 47 | 2801829 | 331962 | 175010 | 7 | 32.71 | 2x150bp | |
| BPH2946 | SAB | AUS/NZ | 8 | 239 | 50 | 3021315 | 46 | 3019717 | 544502 | 155449 | 5 | 32.65 | 2x150bp | |
| BPH2947 | SAB | AUS/NZ | 8 | 239 | 67 | 3022687 | 60 | 3020009 | 407026 | 142918 | 7 | 32.66 | 2x150bp | |
| BPH2949 | SAB | AUS/NZ | 1 | 1 | 26 | 2799712 | 22 | 2798030 | 419106 | 224894 | 5 | 32.69 | 2x300bp | |
| BPH2950 | IE | AUS/NZ | 1 | 188 | 100 | 2709127 | 92 | 2705924 | 144581 | 57224 | 16 | 32.76 | 2x250bp | |
| BPH2951 | SAB | AUS/NZ | 8 | 340 | 32 | 2733774 | 25 | 2730803 | 651157 | 202892 | 4 | 32.66 | 2x150bp | |
| BPH2952 | SAB | AUS/NZ | 45 | 45 | 29 | 2778311 | 21 | 2775425 | 697433 | 232956 | 3 | 32.75 | 2x150bp | |
| BPH2954 | SAB | AUS/NZ | 1 | 1 | 28 | 2737574 | 21 | 2734730 | 595008 | 232337 | 4 | 32.67 | 2x300bp | |
| BPH2955 | SAB | AUS/NZ | 5 | 5 | 70 | 2748806 | 67 | 2747707 | 214468 | 88723 | 10 | 32.75 | 2x250bp | |
| BPH2956 | SAB | AUS/NZ | 22 | 22 | 38 | 2813128 | 36 | 2812230 | 407024 | 165366 | 5 | 32.71 | 2x250bp | |
| BPH2957 | SAB | AUS/NZ | 8 | 8 | 54 | 2752991 | 46 | 2749872 | 346878 | 138371 | 7 | 32.66 | 2x250bp | |
| BPH2958 | SAB | AUS/NZ | 15 | 15 | 47 | 2723638 | 42 | 2721870 | 252552 | 138014 | 8 | 32.75 | 2x300bp | |
| BPH2959 | SAB | AUS/NZ | 5 | 5 | 30 | 2737493 | 25 | 2735550 | 552055 | 296621 | 4 | 32.76 | 2x150bp | |
| BPH2962 | IE | AUS/NZ | 20 | 20 | 28 | 2755983 | 21 | 2753511 | 716297 | 201130 | 3 | 32.67 | 2x150bp | |
| BPH2967 | SAB | AUS/NZ | 78 | 78 | 20 | 2720394 | 14 | 2718064 | 637510 | 353348 | 3 | 32.72 | 2x150bp | |
| BPH2968 | SAB | AUS/NZ | 8 | 8 | 55 | 2801220 | 48 | 2798559 | 411761 | 156235 | 5 | 32.63 | 2x300bp | |
| BPH2969 | SAB | AUS/NZ | 1 | 188 | 23 | 2792754 | 20 | 2791508 | 1056435 | 408155 | 2 | 32.70 | 2x300bp | |
| BPH2972 | SAB | AUS/NZ | 15 | 582 | 26 | 2723717 | 20 | 2721561 | 611381 | 273066 | 4 | 32.73 | 2x150bp | |
| BPH2973 | SAB | AUS/NZ | 30 | 39 | 39 | 2777110 | 32 | 2774596 | 455358 | 175871 | 5 | 32.81 | 2x150bp | |
| BPH2983 | SAB | AUS/NZ | 25 | 25 | 43 | 2777938 | 39 | 2776239 | 309377 | 111070 | 7 | 32.68 | 2x250bp | |
| BPH2984 | SAB | AUS/NZ | 15 | 15 | 50 | 2801382 | 43 | 2798777 | 315685 | 158662 | 6 | 32.76 | 2x250bp | |
| BPH2985 | SAB | AUS/NZ | 5 | 5 | 47 | 2756799 | 44 | 2755787 | 352435 | 105038 | 6 | 32.74 | 2x250bp | |
| BPH2986 | SAB | AUS/NZ | 8 | 8 | 31 | 2892524 | 26 | 2890466 | 938805 | 543165 | 2 | 32.67 | 2x150bp | |
| BPH2987 | IE | AUS/NZ | 30 | 39 | 34 | 2784638 | 27 | 2782090 | 455332 | 225278 | 5 | 32.76 | 2x150bp | |
| BPH2989 | SAB | AUS/NZ | 8 | 2176 | 27 | 2814705 | 21 | 2812423 | 757910 | 313828 | 3 | 32.68 | 2x150bp | |
| BPH2992 | SAB | AUS/NZ | 45 | 10017* | 22 | 2767769 | 21 | 2767280 | 617395 | 316520 | 4 | 32.79 | 2x250bp | *Unknown ST |
| BPH2993 | SAB | AUS/NZ | 45 | 1970 | 25 | 2773796 | 23 | 2772870 | 544785 | 353028 | 4 | 32.82 | 2x300bp | |
| BPH2994 | SAB | AUS/NZ | 20 | 20 | 31 | 2788383 | 28 | 2787184 | 345898 | 250364 | 5 | 32.66 | 2x300bp | |
| BPH3201 | IE | AUS/NZ | 45 | 45 | 41 | 2722471 | 22 | 2715635 | 986563 | 421915 | 2 | 32.75 | 2x150bp | |
| BPH3203 | SAB | AUS/NZ | 101 | 101 | 25 | 2754987 | 20 | 2753237 | 370933 | 229526 | 5 | 32.74 | 2x150bp | |
| BPH3204 | SAB | AUS/NZ | 8 | 1282 | 77 | 2827867 | 70 | 2825371 | 373519 | 103759 | 8 | 32.75 | 2x150bp | |
| BPH3205 | SAB | AUS/NZ | 30 | 30 | 55 | 2789948 | 44 | 2785825 | 483485 | 175535 | 4 | 32.70 | 2x150bp | |
| BPH3207 | SAB | AUS/NZ | 1 | 188 | 51 | 2803202 | 20 | 2791916 | 684228 | 384960 | 3 | 32.74 | 2x150bp | |
| BPH3209 | SAB | AUS/NZ | 15 | 15 | 51 | 2735418 | 35 | 2729698 | 665224 | 215627 | 4 | 32.71 | 2x150bp | |
| BPH3210 | SAB | AUS/NZ | 22 | 1784 | 72 | 2811219 | 46 | 2801315 | 397547 | 150832 | 6 | 32.74 | 2x150bp | |
| BPH3211 | IE | AUS/NZ | 1 | 10018* | 34 | 2734547 | 21 | 2729950 | 669796 | 339403 | 3 | 32.67 | 2x150bp | *Unknown ST |
| BPH3212 | SAB | AUS/NZ | 22 | 22 | 105 | 2862635 | 55 | 2843011 | 401744 | 170707 | 6 | 32.76 | 2x150bp | |
| BPH3213 | SAB | AUS/NZ | 1 | 1 | 46 | 2736253 | 20 | 2726667 | 1245853 | 371089 | 2 | 32.73 | 2x150bp | |
| BPH3214 | SAB | AUS/NZ | 5 | 5 | 55 | 2767530 | 32 | 2758774 | 321768 | 159966 | 7 | 32.73 | 2x150bp | |
| BPH3216 | SAB | AUS/NZ | 1 | 188 | 24 | 2789512 | 17 | 2787065 | 681496 | 462580 | 3 | 32.70 | 2x150bp | |
| BPH3218 | SAB | AUS/NZ | 15 | 15 | 65 | 2768368 | 41 | 2759637 | 321651 | 174040 | 6 | 32.82 | 2x150bp | |
| BPH3221 | SAB | AUS/NZ | 1 | 188 | 40 | 2824116 | 24 | 2818249 | 515735 | 277296 | 4 | 32.72 | 2x150bp | |
| BPH3222 | SAB | AUS/NZ | 45 | 45 | 40 | 2721124 | 24 | 2715181 | 997189 | 423335 | 2 | 32.77 | 2x150bp | |
| BPH3223 | SAB | AUS/NZ | 59 | 59 | 51 | 2719449 | 45 | 2717161 | 317493 | 117961 | 8 | 32.70 | 2x150bp | |
| BPH3224 | SAB | AUS/NZ | 5 | 6 | 54 | 2780823 | 27 | 2770600 | 556399 | 293718 | 4 | 32.74 | 2x150bp | |
| BPH3227 | SAB | AUS/NZ | 5 | 5 | 39 | 2732040 | 31 | 2729021 | 387079 | 186600 | 5 | 32.69 | 2x150bp | |
| BPH3228 | SAB | AUS/NZ | 1 | 1 | 54 | 2893442 | 38 | 2887313 | 378072 | 292191 | 5 | 32.74 | 2x150bp | |
| BPH3229 | SAB | AUS/NZ | 30 | 30 | 94 | 2828771 | 55 | 2814408 | 331411 | 150820 | 7 | 32.76 | 2x150bp | |
| BPH3230 | SAB | AUS/NZ | 30 | 30 | 110 | 2836677 | 58 | 2817130 | 433396 | 151518 | 5 | 32.82 | 2x150bp | |
| BPH3231 | SAB | AUS/NZ | 12 | 12 | 86 | 2747601 | 43 | 2731828 | 856479 | 619752 | 2 | 32.90 | 2x150bp | |
| BPH3232 | SAB | AUS/NZ | 5 | 5 | 33 | 2849640 | 26 | 2847007 | 758631 | 265639 | 4 | 32.70 | 2x150bp | |
| BPH3233 | IE | AUS/NZ | 45 | 508 | 69 | 2888945 | 41 | 2877948 | 572588 | 442561 | 3 | 32.79 | 2x150bp | |
| BPH3234 | SAB | AUS/NZ | 8 | 239 | 74 | 2988873 | 57 | 2982207 | 559967 | 168768 | 5 | 32.61 | 2x150bp | |
| BPH3236 | SAB | AUS/NZ | 8 | 239 | 120 | 2918899 | 87 | 2906390 | 501370 | 109414 | 6 | 32.78 | 2x150bp | |
| BPH3237 | SAB | AUS/NZ | 101 | 101 | 94 | 2784522 | 40 | 2763677 | 910964 | 233169 | 3 | 32.83 | 2x150bp | |
| BPH3238 | IE | AUS/NZ | 1 | 188 | 36 | 2765446 | 29 | 2763103 | 486696 | 208973 | 4 | 32.72 | 2x150bp | |
| BPH3239 | SAB | AUS/NZ | 12 | 12 | 36 | 2677126 | 33 | 2676090 | 461458 | 162090 | 5 | 32.75 | 2x150bp | |
| BPH3240 | SAB | AUS/NZ | 5 | 5 | 40 | 2772369 | 38 | 2771538 | 327873 | 160279 | 7 | 32.70 | 2x150bp | |
| BPH3241 | SAB | AUS/NZ | 5 | 5 | 50 | 2781339 | 39 | 2777393 | 399547 | 223073 | 5 | 32.72 | 2x150bp | |
| BPH3242 | SAB | AUS/NZ | 30 | 30 | 83 | 2850425 | 66 | 2843745 | 462452 | 116132 | 7 | 32.69 | 2x150bp | |
| BPH3243 | SAB | AUS/NZ | 45 | 45 | 52 | 2757955 | 35 | 2751190 | 708713 | 311527 | 3 | 32.71 | 2x150bp | |
| BPH3245 | SAB | AUS/NZ | 78 | 78 | 41 | 2771054 | 36 | 2769043 | 385799 | 216687 | 5 | 32.70 | 2x150bp | |
| BPH3246 | SAB | AUS/NZ | 1 | 1 | 63 | 2795173 | 55 | 2792074 | 323009 | 109342 | 9 | 32.70 | 2x150bp | |
| BPH3248 | SAB | AUS/NZ | 1 | 1 | 127 | 2902142 | 53 | 2873820 | 291574 | 145599 | 7 | 32.74 | 2x150bp | |
| BPH3249 | IE | AUS/NZ | 1 | 188 | 65 | 2755424 | 32 | 2743847 | 385423 | 211270 | 5 | 32.69 | 2x150bp | |
| BPH3250 | SAB | AUS/NZ | 8 | 239 | 103 | 2895379 | 97 | 2893003 | 317029 | 103623 | 9 | 32.72 | 2x150bp | |
| BPH3251 | SAB | AUS/NZ | 50 | 50 | 50 | 2737810 | 48 | 2737070 | 395134 | 143879 | 7 | 32.71 | 2x150bp | |
| BPH3252 | SAB | AUS/NZ | 5 | 5 | 63 | 2832884 | 58 | 2830771 | 195889 | 112990 | 10 | 32.73 | 2x150bp | |
| BPH3253 | SAB | AUS/NZ | 5 | 6 | 56 | 2718891 | 51 | 2716866 | 249189 | 103870 | 9 | 32.71 | 2x150bp | |
| BPH3254 | SAB | AUS/NZ | 8 | 239 | 71 | 2880168 | 66 | 2878242 | 195287 | 103336 | 11 | 32.72 | 2x150bp | |

| | | | | | | | | | | | | | |
|---|---|---|---|---|---|---|---|---|---|---|---|---|---|
| BPH3256 | SAB | AUS/NZ | 59 | 59 | 53 | 2685324 | 50 | 2684228 | 209625 | 119250 | 9 | 32.72 | 2x150bp |
| BPH3257 | SAB | AUS/NZ | 1 | 188 | 64 | 2748953 | 41 | 2740791 | 352470 | 110892 | 7 | 32.69 | 2x150bp |
| BPH3258 | SAB | AUS/NZ | 8 | 239 | 108 | 2901679 | 97 | 2897572 | 378318 | 72982 | 10 | 32.70 | 2x150bp |
| BPH3260 | SAB | AUS/NZ | 8 | 239 | 101 | 2898236 | 96 | 2896400 | 226328 | 77047 | 11 | 32.72 | 2x150bp |
| BPH3261 | SAB | AUS/NZ | 15 | 15 | 71 | 2750980 | 56 | 2745085 | 249492 | 86570 | 11 | 32.72 | 2x150bp |
| BPH3262 | SAB | AUS/NZ | 8 | 239 | 103 | 2884455 | 91 | 2879669 | 260357 | 72962 | 11 | 32.69 | 2x150bp |
| BPH3265 | SAB | AUS/NZ | 20 | 20 | 51 | 2763515 | 42 | 2760438 | 288637 | 111764 | 8 | 32.69 | 2x150bp |
| BPH3266 | SAB | AUS/NZ | 8 | 239 | 91 | 2836189 | 89 | 2835431 | 317735 | 66731 | 12 | 32.73 | 2x150bp |
| BPH3269 | SAB | AUS/NZ | 30 | 30 | 90 | 2809640 | 85 | 2807574 | 246067 | 70759 | 12 | 32.71 | 2x150bp |
| BPH3270 | SAB | AUS/NZ | 1 | 188 | 43 | 2784902 | 39 | 2783393 | 299796 | 134712 | 7 | 32.70 | 2x150bp |
| BPH3271 | SAB | AUS/NZ | 101 | 101 | 52 | 2728531 | 46 | 2726264 | 340252 | 101432 | 9 | 32.71 | 2x150bp |
| BPH3272 | SAB | AUS/NZ | 8 | 239 | 95 | 2873521 | 93 | 2872593 | 310291 | 65715 | 11 | 32.75 | 2x150bp |
| BPH3274 | SAB | AUS/NZ | 5 | 5 | 60 | 2755678 | 35 | 2746209 | 558266 | 290190 | 4 | 32.73 | 2x150bp |
| BPH3275 | SAB | AUS/NZ | 1 | 1 | 44 | 2793339 | 30 | 2788135 | 544763 | 143554 | 6 | 32.68 | 2x150bp |
| BPH3277 | SAB | AUS/NZ | 25 | 25 | 34 | 2757516 | 32 | 2756705 | 403558 | 204506 | 5 | 32.63 | 2x150bp |
| BPH3278 | SAB | AUS/NZ | 59 | 59 | 64 | 2678880 | 56 | 2676011 | 267544 | 92638 | 10 | 32.74 | 2x150bp |
| BPH3279 | SAB | AUS/NZ | 1 | 1 | 46 | 2790951 | 30 | 2785087 | 515133 | 205099 | 5 | 32.68 | 2x150bp |
| BPH3280 | SAB | AUS/NZ | 25 | 25 | 75 | 2730129 | 61 | 2724406 | 222406 | 105764 | 10 | 32.62 | 2x150bp |
| BPH3281 | SAB | AUS/NZ | 78 | 78 | 30 | 2738746 | 22 | 2735574 | 914358 | 228860 | 3 | 32.67 | 2x150bp |
| BPH3282 | SAB | AUS/NZ | 5 | 5 | 45 | 2739352 | 41 | 2737594 | 259174 | 176347 | 7 | 32.69 | 2x150bp |
| BPH3283 | SAB | AUS/NZ | 15 | 15 | 39 | 2713344 | 37 | 2712458 | 319818 | 138249 | 6 | 32.72 | 2x150bp |
| BPH3284 | SAB | AUS/NZ | 8 | 239 | 96 | 2875080 | 91 | 2872943 | 205899 | 90276 | 11 | 32.75 | 2x150bp |
| BPH3285 | SAB | AUS/NZ | 121 | 121 | 72 | 2788239 | 64 | 2785249 | 286085 | 109573 | 9 | 32.70 | 2x150bp |
| BPH3286 | SAB | AUS/NZ | 22 | 22 | 66 | 2782737 | 60 | 2780254 | 223983 | 99745 | 10 | 32.73 | 2x150bp |
| BPH3287 | IE | AUS/NZ | 78 | 78 | 55 | 2723901 | 52 | 2722812 | 218040 | 87273 | 10 | 32.68 | 2x150bp |
| BPH3288 | SAB | AUS/NZ | 1 | 1 | 78 | 2827222 | 42 | 2813769 | 324582 | 183386 | 6 | 32.73 | 2x150bp |
| BPH3289 | SAB | AUS/NZ | 30 | 39 | 38 | 2723498 | 37 | 2723058 | 305410 | 130150 | 7 | 32.76 | 2x150bp |
| BPH3290 | SAB | AUS/NZ | 8 | 1282 | 78 | 2840736 | 78 | 2840736 | 312352 | 82863 | 10 | 32.76 | 2x150bp |
| BPH3292 | SAB | AUS/NZ | 8 | 1282 | 63 | 2778754 | 63 | 2778754 | 275842 | 61309 | 10 | 32.79 | 2x150bp |
| BPH3293 | IE | AUS/NZ | 45 | 45 | 52 | 2760520 | 42 | 2756824 | 563353 | 238486 | 4 | 32.85 | 2x150bp |
| BPH3294 | SAB | AUS/NZ | 1 | 1 | 48 | 2829853 | 42 | 2827466 | 356146 | 132929 | 7 | 32.67 | 2x150bp |
| BPH3295 | SAB | AUS/NZ | 5 | 5 | 53 | 2780818 | 41 | 2776493 | 387123 | 148158 | 6 | 32.70 | 2x150bp |
| BPH3296 | SAB | AUS/NZ | 93 | 93 | 46 | 2802139 | 42 | 2800596 | 338010 | 119859 | 8 | 32.67 | 2x150bp |
| BPH3297 | SAB | AUS/NZ | 78 | 78 | 40 | 2774491 | 36 | 2772763 | 468501 | 201728 | 5 | 32.72 | 2x150bp |
| BPH3298 | SAB | AUS/NZ | 93 | 93 | 137 | 2852236 | 71 | 2827365 | 351617 | 167667 | 6 | 32.94 | 2x150bp |
| BPH3300 | SAB | AUS/NZ | 1 | 1 | 135 | 2831329 | 62 | 2804254 | 322149 | 186194 | 6 | 32.88 | 2x150bp |
| BPH3304 | SAB | AUS/NZ | 78 | 88 | 42 | 2731864 | 37 | 2729927 | 312305 | 222870 | 5 | 32.67 | 2x150bp |
| BPH3305 | SAB | AUS/NZ | 1 | 188 | 120 | 2822697 | 48 | 2796270 | 688908 | 186281 | 4 | 32.88 | 2x150bp |
| BPH3310 | SAB | AUS/NZ | 30 | 30 | 100 | 2833360 | 91 | 2829445 | 145106 | 60694 | 15 | 32.74 | 2x150bp |
| BPH3311 | SAB | AUS/NZ | 20 | 20 | 125 | 2773839 | 74 | 2754428 | 223286 | 93423 | 10 | 32.77 | 2x150bp |
| BPH3312 | SAB | AUS/NZ | 45 | 45 | 76 | 2812519 | 47 | 2802166 | 508274 | 185000 | 5 | 32.81 | 2x150bp |
| BPH3313 | SAB | AUS/NZ | 20 | 20 | 64 | 2778037 | 43 | 2770095 | 297967 | 105931 | 8 | 32.70 | 2x150bp |
| BPH3314 | SAB | AUS/NZ | 20 | 20 | 52 | 2712007 | 49 | 2710911 | 393111 | 105446 | 8 | 32.73 | 2x150bp |
| BPH3315 | SAB | AUS/NZ | 1 | 1 | 24 | 2740384 | 18 | 2738144 | 554223 | 421652 | 3 | 32.66 | 2x150bp |
| BPH3318 | SAB | AUS/NZ | 5 | 5 | 44 | 2763419 | 35 | 2760145 | 340136 | 170079 | 6 | 32.73 | 2x150bp |
| BPH3319 | IE | AUS/NZ | 78 | 88 | 68 | 2824039 | 57 | 2819338 | 237699 | 106213 | 9 | 32.79 | 2x150bp |
| BPH3320 | SAB | AUS/NZ | 8 | 1282 | 215 | 2886368 | 133 | 2854698 | 302564 | 72030 | 10 | 33.10 | 2x150bp |
| BPH3321 | SAB | AUS/NZ | 5 | 5 | 82 | 2740429 | 49 | 2728529 | 397492 | 148567 | 6 | 32.71 | 2x150bp |
| BPH3323 | SAB | AUS/NZ | 5 | 5 | 55 | 2772825 | 52 | 2771753 | 310946 | 129884 | 7 | 32.72 | 2x150bp |
| BPH3325 | SAB | AUS/NZ | 1 | 1 | 45 | 2756331 | 37 | 2753275 | 285714 | 148298 | 7 | 32.67 | 2x150bp |
| BPH3326 | SAB | AUS/NZ | 8 | 8 | 96 | 2834565 | 89 | 2831888 | 211198 | 96501 | 11 | 32.70 | 2x150bp |
| BPH3327 | SAB | AUS/NZ | 15 | 15 | 40 | 2716845 | 38 | 2716019 | 306342 | 128653 | 8 | 32.71 | 2x150bp |
| BPH3329 | SAB | AUS/NZ | 5 | 5 | 139 | 2813606 | 89 | 2794007 | 319637 | 138557 | 7 | 33.10 | 2x150bp |
| BPH3331 | SAB | AUS/NZ | 15 | 15 | 48 | 2674972 | 39 | 2671420 | 271976 | 141934 | 7 | 32.70 | 2x150bp |
| BPH3332 | SAB | AUS/NZ | 8 | 239 | 106 | 3013991 | 100 | 3011768 | 247594 | 61030 | 13 | 32.66 | 2x150bp |
| BPH3333 | SAB | AUS/NZ | 97 | 97 | 54 | 2715279 | 34 | 2708239 | 418894 | 163348 | 5 | 32.73 | 2x150bp |
| BPH3334 | SAB | AUS/NZ | 5 | 5 | 56 | 2722606 | 53 | 2721488 | 270847 | 110280 | 9 | 32.76 | 2x150bp |
| BPH3335 | SAB | AUS/NZ | 30 | 30 | 76 | 2807224 | 64 | 2802486 | 198870 | 95310 | 11 | 32.72 | 2x150bp |
| BPH3336 | SAB | AUS/NZ | 5 | 5 | 40 | 2719800 | 38 | 2719003 | 515697 | 152845 | 6 | 32.73 | 2x150bp |
| BPH3337 | IE | AUS/NZ | 8 | 8 | 76 | 2795728 | 68 | 2792451 | 248869 | 76943 | 11 | 32.67 | 2x150bp |
| BPH3339 | SAB | AUS/NZ | 12 | 12 | 46 | 2710634 | 42 | 2709283 | 416078 | 179098 | 5 | 32.71 | 2x150bp |
| BPH3340 | SAB | AUS/NZ | 45 | 45 | 103 | 2900590 | 82 | 2892718 | 245033 | 95672 | 9 | 32.73 | 2x150bp |
| BPH3341 | SAB | AUS/NZ | 45 | 45 | 29 | 2689106 | 19 | 2685475 | 986988 | 404243 | 2 | 32.77 | 2x150bp |
| BPH3343 | SAB | AUS/NZ | 15 | 15 | 78 | 2746704 | 72 | 2744369 | 190422 | 73504 | 12 | 32.75 | 2x150bp |
| BPH3344 | SAB | AUS/NZ | 8 | 239 | 89 | 2897195 | 83 | 2894915 | 300212 | 76383 | 11 | 32.63 | 2x150bp |
| BPH3345 | IE | AUS/NZ | 30 | 30 | 72 | 2818308 | 66 | 2815911 | 198870 | 97389 | 10 | 32.74 | 2x150bp |
| BPH3346 | SAB | AUS/NZ | 8 | 239 | 109 | 3011909 | 97 | 3007453 | 247496 | 74906 | 12 | 32.65 | 2x150bp |
| BPH3348 | SAB | AUS/NZ | 8 | 239 | 96 | 3012214 | 80 | 3005978 | 271688 | 109558 | 9 | 32.65 | 2x150bp |
| BPH3351 | SAB | AUS/NZ | 30 | 30 | 77 | 2853481 | 66 | 2849140 | 174006 | 116122 | 10 | 32.76 | 2x150bp |
| BPH3352 | IE | AUS/NZ | 45 | 45 | 63 | 2871353 | 49 | 2866269 | 299644 | 159976 | 7 | 32.69 | 2x150bp |
| BPH3354 | SAB | AUS/NZ | 20 | 20 | 47 | 2720710 | 42 | 2718958 | 341477 | 127001 | 7 | 32.70 | 2x150bp |
| BPH3355 | SAB | AUS/NZ | 15 | 15 | 60 | 2784954 | 60 | 2784954 | 309651 | 104393 | 9 | 32.75 | 2x150bp |
| BPH3356 | SAB | AUS/NZ | 8 | 239 | 83 | 3001406 | 74 | 2997758 | 291319 | 109371 | 10 | 32.63 | 2x150bp |
| BPH3357 | SAB | AUS/NZ | 8 | 1152 | 64 | 2767881 | 57 | 2765200 | 188270 | 104352 | 10 | 32.67 | 2x150bp |
| BPH3359 | SAB | AUS/NZ | 5 | 5 | 75 | 2840007 | 68 | 2837164 | 269839 | 76022 | 11 | 32.73 | 2x150bp |
| BPH3361 | SAB | AUS/NZ | 8 | 8 | 52 | 2746926 | 47 | 2744864 | 483811 | 170267 | 5 | 32.67 | 2x150bp |
| BPH3363 | SAB | AUS/NZ | 101 | 101 | 30 | 2703370 | 26 | 2701789 | 353602 | 241862 | 5 | 32.73 | 2x150bp |
| BPH3365 | SAB | AUS/NZ | 12 | 12 | 30 | 2711090 | 26 | 2709658 | 461233 | 284578 | 4 | 32.71 | 2x150bp |

| | | | | | | | | | | | | | | |
|---|---|---|---|---|---|---|---|---|---|---|---|---|---|---|
| BPH3366 | SAB | AUS/NZ | 8 | 239 | 100 | 2985300 | 91 | 2981514 | 222679 | 101393 | 11 | 32.64 | 2x150bp | |
| BPH3367 | IE | AUS/NZ | 101 | 101 | 50 | 2703911 | 48 | 2703207 | 384032 | 103910 | 8 | 32.74 | 2x150bp | |
| BPH3368 | SAB | AUS/NZ | 8 | 239 | 73 | 2988235 | 72 | 2987893 | 404762 | 94930 | 9 | 32.66 | 2x150bp | |
| BPH3369 | SAB | AUS/NZ | 78 | 78 | 36 | 2733002 | 33 | 2731812 | 292031 | 156815 | 7 | 32.69 | 2x150bp | |
| BPH3370 | SAB | AUS/NZ | 8 | 239 | 93 | 3008553 | 86 | 3005735 | 208304 | 100764 | 11 | 32.64 | 2x150bp | |
| BPH3371 | SAB | AUS/NZ | 30 | 39 | 42 | 2767715 | 41 | 2767275 | 325023 | 124280 | 7 | 32.78 | 2x150bp | |
| BPH3372 | SAB | AUS/NZ | 8 | 239 | 101 | 3019075 | 81 | 3011537 | 300702 | 100796 | 8 | 32.65 | 2x150bp | |
| BPH3373 | SAB | AUS/NZ | 30 | 30 | 84 | 2828469 | 69 | 2822739 | 296468 | 116184 | 8 | 32.70 | 2x150bp | |
| BPH3374 | SAB | AUS/NZ | 5 | 6 | 47 | 2724324 | 43 | 2722820 | 339645 | 108571 | 8 | 32.71 | 2x150bp | |
| BPH3376 | SAB | AUS/NZ | 8 | 239 | 126 | 3002234 | 113 | 2997380 | 142889 | 69315 | 15 | 32.67 | 2x150bp | |
| BPH3378 | SAB | AUS/NZ | 45 | 45 | 52 | 2725031 | 49 | 2723985 | 334154 | 88328 | 9 | 32.73 | 2x150bp | |
| BPH3379 | SAB | AUS/NZ | 78 | 78 | 33 | 2731300 | 25 | 2728080 | 353113 | 232460 | 5 | 32.69 | 2x150bp | |
| BPH3380 | SAB | AUS/NZ | 7 | 7 | 45 | 2731758 | 37 | 2728886 | 318874 | 142893 | 7 | 32.69 | 2x150bp | |
| BPH3381 | SAB | AUS/NZ | 45 | 1795 | 39 | 2757902 | 32 | 2755432 | 427918 | 254089 | 5 | 32.80 | 2x150bp | |
| BPH3382 | SAB | AUS/NZ | 8 | 239 | 81 | 2937246 | 74 | 2934269 | 216834 | 101148 | 10 | 32.66 | 2x150bp | |
| BPH3384 | SAB | AUS/NZ | 22 | 22 | 89 | 2825564 | 62 | 2816148 | 289310 | 102664 | 9 | 32.70 | 2x150bp | |
| BPH3390 | SAB | AUS/NZ | 45 | 45 | 38 | 2693896 | 33 | 2691871 | 469240 | 307254 | 4 | 32.68 | 2x150bp | |
| BPH3392 | SAB | AUS/NZ | 12 | 12 | 45 | 2691727 | 41 | 2690273 | 314531 | 165225 | 6 | 32.74 | 2x150bp | |
| BPH3393 | SAB | AUS/NZ | 1 | 188 | 32 | 2729819 | 24 | 2726955 | 483157 | 271746 | 4 | 32.69 | 2x150bp | |
| BPH3395 | SAB | AUS/NZ | 30 | 30 | 68 | 2807790 | 61 | 2805039 | 383611 | 116262 | 7 | 32.72 | 2x150bp | |
| BPH3396 | SAB | AUS/NZ | 22 | 22 | 80 | 2752247 | 57 | 2743296 | 228579 | 99943 | 9 | 32.68 | 2x150bp | |
| BPH3397 | SAB | AUS/NZ | 1 | 1 | 40 | 2736420 | 25 | 2730647 | 421987 | 300791 | 4 | 32.70 | 2x150bp | |
| BPH3399 | SAB | AUS/NZ | 93 | 93 | 40 | 2802234 | 34 | 2799983 | 474323 | 205449 | 5 | 32.66 | 2x150bp | |
| BPH3400 | SAB | AUS/NZ | 1 | 188 | 42 | 2747528 | 21 | 2739430 | 461570 | 277403 | 4 | 32.67 | 2x150bp | |
| BPH3401 | IE | AUS/NZ | 8 | 239 | 82 | 3005592 | 65 | 2999172 | 291198 | 112606 | 8 | 32.66 | 2x150bp | |
| BPH3403 | SAB | AUS/NZ | 8 | 239 | 335 | 3168810 | 164 | 3106033 | 296500 | 111461 | 9 | 32.62 | 2x150bp | |
| BPH3406 | IE | AUS/NZ | 45 | 45 | 60 | 2725595 | 22 | 2711881 | 458971 | 273256 | 4 | 32.76 | 2x150bp | |
| BPH3408 | SAB | AUS/NZ | 8 | 8 | 59 | 2753033 | 47 | 2748487 | 330356 | 175800 | 6 | 32.64 | 2x150bp | |
| BPH3409 | SAB | AUS/NZ | 45 | 45 | 42 | 2701456 | 22 | 2694201 | 987316 | 428923 | 2 | 32.79 | 2x150bp | |
| BPH3410 | IE | AUS/NZ | 12 | 12 | 56 | 2703713 | 26 | 2692594 | 854468 | 325009 | 3 | 32.76 | 2x150bp | |
| BPH3411 | SAB | AUS/NZ | 22 | 22 | 91 | 2810081 | 54 | 2796307 | 327146 | 125502 | 8 | 32.76 | 2x150bp | |
| BPH3412 | SAB | AUS/NZ | 5 | 5 | 52 | 2736651 | 31 | 2729109 | 545272 | 186602 | 5 | 32.69 | 2x150bp | |
| BPH3414 | IE | AUS/NZ | 45 | 45 | 74 | 2747035 | 33 | 2731871 | 527382 | 283600 | 4 | 32.78 | 2x150bp | |
| BPH3416 | SAB | AUS/NZ | 5 | 5 | 60 | 2753804 | 35 | 2744622 | 386897 | 215594 | 5 | 32.69 | 2x150bp | |
| BPH3419 | SAB | AUS/NZ | 5 | 5 | 45 | 2830776 | 32 | 2825463 | 394704 | 188064 | 5 | 32.68 | 2x150bp | |
| BPH3420 | SAB | AUS/NZ | 45 | 508 | 342 | 2887289 | 128 | 2808044 | 510287 | 267834 | 4 | 32.80 | 2x150bp | |
| BPH3422 | IE | AUS/NZ | 5 | 5 | 54 | 2792533 | 39 | 2786560 | 415635 | 157752 | 7 | 32.73 | 2x150bp | |
| BPH3425 | SAB | AUS/NZ | 5 | 6 | 37 | 2732868 | 22 | 2727226 | 490375 | 221750 | 4 | 32.69 | 2x150bp | |
| BPH3426 | IE | AUS/NZ | 5 | 10019* | 37 | 2793189 | 32 | 2791196 | 433242 | 239696 | 5 | 32.83 | 2x150bp | *Unknown ST |
| BPH3428 | SAB | AUS/NZ | 5 | 5 | 51 | 2782514 | 43 | 2779676 | 387075 | 221756 | 5 | 32.70 | 2x150bp | |
| BPH3430 | SAB | AUS/NZ | 45 | 45 | 31 | 2737830 | 27 | 2736299 | 515143 | 385910 | 3 | 32.82 | 2x150bp | |
| BPH3432 | SAB | AUS/NZ | 12 | 12 | 39 | 2711439 | 17 | 2703379 | 974678 | 666074 | 2 | 32.72 | 2x150bp | |
| BPH3434 | SAB | AUS/NZ | 20 | 20 | 52 | 2747260 | 42 | 2743525 | 570177 | 166990 | 5 | 32.65 | 2x150bp | |
| BPH3438 | SAB | AUS/NZ | 121 | 121 | 86 | 2807297 | 65 | 2799207 | 263737 | 100926 | 9 | 32.74 | 2x150bp | |
| BPH3440 | SAB | AUS/NZ | 1 | 1 | 42 | 2759544 | 29 | 2754600 | 468672 | 281015 | 4 | 32.68 | 2x150bp | |
| BPH3441 | SAB | AUS/NZ | 1 | 9 | 60 | 2766631 | 46 | 2761225 | 890447 | 230250 | 3 | 32.74 | 2x150bp | |
| BPH3442 | SAB | AUS/NZ | 30 | 34 | 75 | 2792189 | 59 | 2786388 | 215569 | 124624 | 9 | 32.70 | 2x150bp | |
| BPH3443 | SAB | AUS/NZ | 8 | 239 | 90 | 3019505 | 75 | 3013726 | 271552 | 101278 | 10 | 32.63 | 2x150bp | |
| BPH3447 | SAB | AUS/NZ | 1 | 1 | 49 | 2798963 | 23 | 2788695 | 539932 | 421225 | 3 | 32.71 | 2x150bp | |
| BPH3451 | SAB | AUS/NZ | 8 | 239 | 88 | 2987256 | 78 | 2983410 | 277968 | 97365 | 10 | 32.69 | 2x150bp | |
| BPH3455 | SAB | AUS/NZ | 8 | 239 | 96 | 3024227 | 82 | 3019169 | 406309 | 109546 | 8 | 32.65 | 2x150bp | |
| BPH3456 | IE | AUS/NZ | 78 | 88 | 39 | 2744738 | 29 | 2740862 | 665724 | 188689 | 4 | 32.75 | 2x150bp | |
| BPH3458 | SAB | AUS/NZ | 15 | 15 | 45 | 2764158 | 35 | 2760411 | 427059 | 138089 | 6 | 32.74 | 2x150bp | |
| BPH3463 | SAB | AUS/NZ | 8 | 239 | 126 | 3037334 | 101 | 3027506 | 257991 | 101882 | 9 | 32.68 | 2x150bp | |
| BPH3466 | IE | AUS/NZ | 45 | 45 | 21 | 2691386 | 17 | 2689918 | 1087690 | 514908 | 2 | 32.78 | 2x150bp | |
| BPH3467 | SAB | AUS/NZ | 5 | 5 | 103 | 2866665 | 65 | 2853198 | 368841 | 106191 | 9 | 32.64 | 2x150bp | |
| BPH3468 | SAB | AUS/NZ | 30 | 30 | 355 | 3083276 | 234 | 3037643 | 273120 | 74644 | 11 | 32.82 | 2x150bp | |
| BPH3470 | SAB | AUS/NZ | 1 | 1 | 96 | 2736144 | 85 | 2731973 | 143843 | 63096 | 15 | 32.67 | 2x150bp | |
| BPH3472 | SAB | AUS/NZ | 5 | 3841 | 29 | 2766382 | 23 | 2763917 | 387539 | 222569 | 5 | 32.73 | 2x150bp | |
| BPH3473 | SAB | AUS/NZ | 8 | 239 | 61 | 3017327 | 55 | 3014982 | 559972 | 169318 | 5 | 32.63 | 2x150bp | |
| BPH3474 | IE | AUS/NZ | 30 | 34 | 118 | 2860382 | 102 | 2853990 | 273994 | 67754 | 13 | 32.71 | 2x150bp | |
| BPH3476 | SAB | AUS/NZ | 15 | 15 | 79 | 2827911 | 66 | 2823132 | 455348 | 214947 | 5 | 32.78 | 2x150bp | |
| BPH3477 | SAB | AUS/NZ | 45 | 45 | 44 | 2830471 | 36 | 2827445 | 568566 | 201047 | 5 | 32.66 | 2x150bp | |
| BPH3478 | IE | AUS/NZ | 5 | 6 | 27 | 2725096 | 22 | 2723124 | 430704 | 203406 | 5 | 32.70 | 2x150bp | |
| BPH3479 | SAB | AUS/NZ | 45 | 45 | 38 | 2829924 | 33 | 2827930 | 508425 | 276477 | 4 | 32.66 | 2x150bp | |
| BPH3480 | SAB | AUS/NZ | 5 | 5 | 81 | 2760397 | 49 | 2748775 | 327944 | 135553 | 7 | 32.67 | 2x150bp | |
| BPH3481 | SAB | AUS/NZ | 22 | 22 | 50 | 2757380 | 44 | 2755098 | 252234 | 148002 | 7 | 32.68 | 2x150bp | |
| BPH3482 | SAB | AUS/NZ | 20 | 20 | 37 | 2752038 | 28 | 2748742 | 393593 | 236492 | 5 | 32.66 | 2x150bp | |
| BPH3486 | SAB | AUS/NZ | 1 | 188 | 26 | 2757150 | 26 | 2757150 | 486702 | 254441 | 4 | 32.65 | 2x150bp | |
| BPH3488 | SAB | AUS/NZ | 1 | 188 | 100 | 2763695 | 91 | 2760410 | 205360 | 67773 | 13 | 32.70 | 2x150bp | |
| BPH3489 | SAB | AUS/NZ | 22 | 22 | 50 | 2847376 | 43 | 2844573 | 348385 | 146402 | 8 | 32.73 | 2x150bp | |
| BPH3492 | SAB | AUS/NZ | 398 | 398 | 33 | 2690800 | 29 | 2689220 | 625406 | 345966 | 3 | 32.78 | 2x150bp | |
| BPH3493 | SAB | AUS/NZ | 8 | 239 | 68 | 3000582 | 62 | 2998274 | 564165 | 142870 | 6 | 32.66 | 2x150bp | |
| BPH3494 | SAB | AUS/NZ | 1 | 1 | 30 | 2742823 | 25 | 2740883 | 523421 | 251991 | 4 | 32.66 | 2x150bp | |
| BPH3497 | SAB | AUS/NZ | 45 | 45 | 47 | 2882341 | 39 | 2879240 | 553733 | 244112 | 4 | 32.66 | 2x150bp | |
| BPH3499 | SAB | AUS/NZ | 8 | 239 | 76 | 3025136 | 68 | 3022184 | 406524 | 155515 | 6 | 32.65 | 2x150bp | |
| BPH3501 | SAB | AUS/NZ | 22 | 22 | 57 | 2847597 | 48 | 2844021 | 348230 | 150802 | 7 | 32.73 | 2x150bp | |
| BPH3502 | IE | AUS/NZ | 395 | 395 | 35 | 2758907 | 31 | 2757446 | 506741 | 250457 | 4 | 32.74 | 2x150bp | |

| | | | | | | | | | | | | | | |
|---|---|---|---|---|---|---|---|---|---|---|---|---|---|---|
| BPH3503 | SAB | AUS/NZ | 78 | 88 | 37 | 2753063 | 30 | 2750194 | 391444 | 188648 | 5 | 32.71 | 2x150bp | |
| BPH3505 | SAB | AUS/NZ | 45 | 45 | 36 | 2842876 | 30 | 2840500 | 508433 | 285623 | 4 | 32.73 | 2x150bp | |
| BPH3506 | SAB | AUS/NZ | 45 | 45 | 33 | 2698913 | 25 | 2695944 | 704791 | 317492 | 3 | 32.75 | 2x150bp | |
| BPH3507 | SAB | AUS/NZ | 8 | 239 | 93 | 2980395 | 87 | 2978061 | 196489 | 104733 | 11 | 32.67 | 2x150bp | |
| BPH3508 | SAB | AUS/NZ | 20 | 20 | 39 | 2753213 | 27 | 2748684 | 382038 | 240814 | 5 | 32.67 | 2x150bp | |
| BPH3509 | SAB | AUS/NZ | 15 | 15 | 41 | 2760194 | 38 | 2758962 | 339694 | 141690 | 6 | 32.74 | 2x150bp | |
| BPH3510 | SAB | AUS/NZ | 22 | 22 | 81 | 2694746 | 66 | 2688976 | 235655 | 78970 | 11 | 32.74 | 2x150bp | |
| BPH3511 | SAB | AUS/NZ | 8 | 239 | 140 | 3033810 | 86 | 3012949 | 400090 | 109382 | 9 | 32.67 | 2x150bp | |
| BPH3512 | SAB | AUS/NZ | 1 | 4100 | 49 | 2691274 | 37 | 2686573 | 324976 | 152524 | 6 | 32.69 | 2x150bp | |
| BPH3513 | SAB | AUS/NZ | 5 | 5 | 32 | 2785831 | 28 | 2784203 | 571561 | 192245 | 4 | 32.72 | 2x150bp | |
| BPH3514 | IE | AUS/NZ | 1 | 188 | 27 | 2731479 | 22 | 2729499 | 469931 | 382114 | 4 | 32.67 | 2x150bp | |
| BPH3516 | SAB | AUS/NZ | 78 | 88 | 46 | 2763357 | 39 | 2760565 | 319084 | 228731 | 5 | 32.72 | 2x150bp | |
| BPH3517 | SAB | AUS/NZ | 45 | 45 | 53 | 2866305 | 46 | 2863488 | 496250 | 156012 | 5 | 32.68 | 2x150bp | |
| BPH3518 | SAB | AUS/NZ | 15 | 15 | 39 | 2765849 | 34 | 2764012 | 490951 | 246188 | 5 | 32.75 | 2x150bp | |
| BPH3520 | IE | AUS/NZ | 1 | 188 | 34 | 2747271 | 28 | 2744932 | 483557 | 226917 | 4 | 32.66 | 2x150bp | |
| BPH3522 | SAB | AUS/NZ | 8 | 8 | 46 | 2746213 | 36 | 2742199 | 793400 | 299695 | 3 | 32.67 | 2x150bp | |
| BPH3524 | SAB | AUS/NZ | 30 | 30 | 73 | 2850728 | 65 | 2847465 | 330397 | 116197 | 8 | 32.73 | 2x150bp | |
| BPH3526 | SAB | AUS/NZ | 15 | 15 | 45 | 2682776 | 35 | 2679099 | 430881 | 165432 | 5 | 32.69 | 2x150bp | |
| BPH3527 | SAB | AUS/NZ | 8 | 239 | 118 | 3008600 | 99 | 3000741 | 199942 | 87364 | 12 | 32.61 | 2x150bp | |
| BPH3531 | SAB | AUS/NZ | 8 | 239 | 91 | 3030199 | 75 | 3024408 | 291291 | 109530 | 9 | 32.62 | 2x150bp | |
| BPH3532 | SAB | AUS/NZ | 15 | 15 | 42 | 2687095 | 31 | 2682975 | 431604 | 160493 | 6 | 32.69 | 2x150bp | |
| BPH3534 | SAB | AUS/NZ | 45 | 45 | 43 | 2779297 | 24 | 2772176 | 1001898 | 421689 | 2 | 32.79 | 2x150bp | |
| BPH3535 | SAB | AUS/NZ | 8 | 239 | 84 | 3021323 | 64 | 3014005 | 300711 | 111624 | 8 | 32.62 | 2x150bp | |
| BPH3537 | SAB | AUS/NZ | 45 | 45 | 80 | 2937905 | 49 | 2926605 | 409596 | 244159 | 5 | 32.69 | 2x150bp | |
| BPH3538 | SAB | AUS/NZ | 78 | 88 | 41 | 2775357 | 31 | 2771351 | 720639 | 188689 | 4 | 32.70 | 2x150bp | |
| BPH3539 | SAB | AUS/NZ | 45 | 45 | 46 | 2833805 | 35 | 2829660 | 507385 | 234554 | 4 | 32.66 | 2x150bp | |
| BPH3540 | IE | AUS/NZ | 8 | 72 | 31 | 2702164 | 24 | 2699428 | 500515 | 318778 | 4 | 32.71 | 2x150bp | |
| BPH3541 | SAB | AUS/NZ | 8 | 239 | 74 | 2821647 | 68 | 2819371 | 299850 | 109085 | 9 | 32.75 | 2x150bp | |
| BPH3542 | SAB | AUS/NZ | 5 | 5 | 40 | 2712304 | 33 | 2709635 | 327211 | 159318 | 6 | 32.69 | 2x150bp | |
| BPH3543 | SAB | AUS/NZ | 8 | 239 | 74 | 2879198 | 66 | 2876168 | 281487 | 106431 | 9 | 32.73 | 2x150bp | |
| BPH3545 | SAB | AUS/NZ | 8 | 239 | 78 | 3022690 | 63 | 3016942 | 298597 | 117369 | 8 | 32.63 | 2x150bp | |
| BPH3546 | SAB | AUS/NZ | 1 | 188 | 26 | 2748127 | 19 | 2745273 | 684189 | 291374 | 3 | 32.66 | 2x150bp | |
| BPH3547 | SAB | AUS/NZ | 8 | 239 | 70 | 3058597 | 60 | 3054644 | 300710 | 142793 | 7 | 32.60 | 2x150bp | |
| BPH3548 | SAB | AUS/NZ | 30 | 30 | 83 | 2854540 | 71 | 2849639 | 331322 | 125079 | 8 | 32.73 | 2x150bp | |
| BPH3549 | SAB | AUS/NZ | 8 | 239 | 76 | 2975028 | 71 | 2973129 | 232823 | 105893 | 11 | 32.68 | 2x150bp | |
| BPH3550 | SAB | AUS/NZ | 78 | 78 | 45 | 2780300 | 37 | 2777140 | 297178 | 157369 | 6 | 32.72 | 2x150bp | |
| BPH3551 | SAB | AUS/NZ | 1 | 1 | 45 | 2827328 | 31 | 2822272 | 379344 | 259967 | 5 | 32.76 | 2x150bp | |
| BPH3552 | SAB | AUS/NZ | 30 | 30 | 53 | 2801647 | 52 | 2801322 | 313176 | 110278 | 9 | 32.71 | 2x150bp | |
| BPH3553 | SAB | AUS/NZ | 78 | 88 | 44 | 2749300 | 36 | 2746166 | 363154 | 149671 | 6 | 32.70 | 2x150bp | |
| BPH3554 | SAB | AUS/NZ | 8 | 3376 | 35 | 2826912 | 29 | 2824587 | 841697 | 278178 | 3 | 32.70 | 2x150bp | |
| BPH3555 | SAB | AUS/NZ | 93 | 93 | 37 | 2803048 | 32 | 2801097 | 474324 | 292587 | 4 | 32.67 | 2x150bp | |
| BPH3557 | SAB | AUS/NZ | 8 | 72 | 36 | 2744081 | 29 | 2741175 | 399727 | 220682 | 5 | 32.65 | 2x150bp | |
| BPH3559 | SAB | AUS/NZ | 45 | 45 | 31 | 2699854 | 21 | 2696232 | 532860 | 454601 | 3 | 32.78 | 2x150bp | |
| BPH3563 | SAB | AUS/NZ | 20 | 20 | 44 | 2715937 | 27 | 2709774 | 760354 | 294475 | 3 | 32.75 | 2x150bp | |
| BPH3565 | SAB | AUS/NZ | 30 | 34 | 68 | 2843154 | 56 | 2838904 | 451283 | 135107 | 7 | 32.72 | 2x150bp | |
| BPH3566 | SAB | AUS/NZ | 22 | 22 | 45 | 2711753 | 35 | 2707964 | 336831 | 172692 | 6 | 32.73 | 2x150bp | |
| BPH3571 | SAB | AUS/NZ | 5 | 5 | 33 | 2682228 | 28 | 2680268 | 534378 | 223115 | 4 | 32.70 | 2x150bp | |
| BPH3572 | SAB | AUS/NZ | 78 | 78 | 831 | 3320596 | 306 | 3124973 | 466550 | 158894 | 6 | 32.63 | 2x150bp | |
| BPH3574 | SAB | AUS/NZ | 22 | 22 | 561 | 3222803 | 333 | 3135902 | 255668 | 70813 | 14 | 32.71 | 2x150bp | |
| BPH3575 | SAB | AUS/NZ | 5 | 5 | 42 | 2735478 | 32 | 2731636 | 386983 | 167533 | 6 | 32.68 | 2x150bp | |
| BPH3577 | SAB | AUS/NZ | 78 | 88 | 417 | 3142930 | 333 | 3110147 | 273228 | 37314 | 23 | 32.58 | 2x150bp | |
| BPH3578 | SAB | AUS/NZ | 78 | 78 | 32 | 2812266 | 25 | 2809475 | 1024313 | 240205 | 3 | 32.70 | 2x150bp | |
| BPH3579 | IE | AUS/NZ | 5 | 5 | 513 | 3148975 | 437 | 3120446 | 110459 | 19904 | 43 | 32.55 | 2x150bp | |
| BPH3580 | SAB | AUS/NZ | 22 | 22 | 60 | 2791903 | 53 | 2789048 | 349217 | 150954 | 6 | 32.72 | 2x150bp | |
| BPH3581 | SAB | AUS/NZ | 15 | 15 | 28 | 2685250 | 25 | 2684197 | 486108 | 221709 | 5 | 32.67 | 2x150bp | |
| BPH3582 | SAB | AUS/NZ | 5 | 73 | 271 | 3023228 | 170 | 2985937 | 287410 | 116656 | 7 | 32.57 | 2x150bp | |
| BPH3583 | IE | AUS/NZ | 1 | 1 | 266 | 3033097 | 170 | 2996614 | 274679 | 118129 | 9 | 32.56 | 2x150bp | |
| BPH3584 | SAB | AUS/NZ | 121 | 121 | 52 | 2830780 | 43 | 2827192 | 263632 | 151106 | 8 | 32.66 | 2x150bp | |
| BPH3585 | SAB | AUS/NZ | 5 | 5 | 39 | 2789160 | 35 | 2787515 | 371209 | 234941 | 5 | 32.69 | 2x150bp | |
| BPH3588 | SAB | AUS/NZ | 121 | 121 | 51 | 2829928 | 43 | 2826835 | 321398 | 147650 | 7 | 32.66 | 2x150bp | |
| BPH3590 | SAB | AUS/NZ | 22 | 22 | 58 | 2853653 | 49 | 2850191 | 209983 | 125484 | 9 | 32.65 | 2x150bp | |
| BPH3593 | SAB | AUS/NZ | 15 | 15 | 34 | 2673563 | 28 | 2671467 | 611666 | 250107 | 4 | 32.69 | 2x150bp | |
| BPH3595 | SAB | AUS/NZ | 97 | 97 | 31 | 2738368 | 27 | 2736752 | 473325 | 199833 | 5 | 32.69 | 2x150bp | |
| BPH3596 | SAB | AUS/NZ | 1 | 10020* | 59 | 2864241 | 51 | 2861235 | 299316 | 143784 | 7 | 32.69 | 2x150bp | *Unknown ST |
| BPH3598 | SAB | AUS/NZ | 291 | 291 | 40 | 2720847 | 33 | 2718149 | 308259 | 194286 | 6 | 32.79 | 2x150bp | |
| BPH3599 | IE | AUS/NZ | 1 | 1 | 24 | 2747820 | 18 | 2745540 | 552433 | 427905 | 3 | 32.66 | 2x150bp | |
| BPH3600 | SAB | AUS/NZ | 78 | 78 | 55 | 2843994 | 46 | 2840359 | 291113 | 193928 | 6 | 32.72 | 2x150bp | |
| BPH3601 | SAB | AUS/NZ | 78 | 78 | 54 | 2765706 | 44 | 2761471 | 280879 | 135561 | 7 | 32.74 | 2x150bp | |
| BPH3603 | SAB | AUS/NZ | 30 | 39 | 71 | 2751806 | 54 | 2744981 | 209724 | 105161 | 10 | 32.69 | 2x150bp | |
| BPH3604 | SAB | AUS/NZ | 22 | 22 | 50 | 2804708 | 44 | 2802259 | 348329 | 125498 | 7 | 32.70 | 2x150bp | |
| BPH3605 | SAB | AUS/NZ | 101 | 101 | 22 | 2736948 | 17 | 2735062 | 928145 | 237543 | 3 | 32.66 | 2x150bp | |
| BPH3607 | SAB | AUS/NZ | 5 | 5 | 49 | 2824528 | 44 | 2822482 | 548700 | 169907 | 5 | 32.73 | 2x150bp | |
| BPH3609 | SAB | AUS/NZ | 15 | 15 | 33 | 2728983 | 29 | 2727284 | 657535 | 193937 | 4 | 32.70 | 2x150bp | |
| BPH3610 | SAB | AUS/NZ | 5 | 5 | 48 | 2784917 | 39 | 2781337 | 481056 | 153123 | 6 | 32.68 | 2x150bp | |
| BPH3611 | SAB | AUS/NZ | 5 | 5 | 39 | 2793747 | 31 | 2790645 | 722539 | 161202 | 4 | 32.67 | 2x150bp | |
| BPH3614 | SAB | AUS/NZ | 1 | 1 | 20 | 2753307 | 16 | 2751831 | 836453 | 569503 | 2 | 32.67 | 2x150bp | |
| BPH3615 | IE | AUS/NZ | 20 | 20 | 53 | 2764750 | 37 | 2758708 | 391392 | 160032 | 6 | 32.65 | 2x150bp | |
| BPH3616 | SAB | AUS/NZ | 5 | 73 | 62 | 2821420 | 41 | 2813783 | 351804 | 159855 | 6 | 32.69 | 2x150bp | |

| | | | | | | | | | | | | | | | |
|---|---|---|---|---|---|---|---|---|---|---|---|---|---|---|---|
| BPH3618 | SAB | AUS/NZ | | 78 | 78 | 32 | 2800848 | 25 | 2798134 | 353231 | 232423 | 5 | 32.70 | 2x150bp | |
| BPH3620 | SAB | AUS/NZ | | 5 | 73 | 46 | 2785795 | 40 | 2783583 | 318978 | 185582 | 6 | 32.64 | 2x150bp | |
| BPH3623 | SAB | AUS/NZ | | 93 | 93 | 43 | 2821280 | 34 | 2817865 | 515159 | 257928 | 4 | 32.72 | 2x150bp | |
| BPH3628 | SAB | AUS/NZ | | 1 | 10020* | 34 | 2863580 | 26 | 2860415 | 593460 | 299352 | 4 | 32.69 | 2x150bp | *Unknown ST |
| BPH3631 | SAB | AUS/NZ | | 93 | 93 | 45 | 2821777 | 38 | 2819168 | 432215 | 205565 | 5 | 32.72 | 2x150bp | |
| BPH3632 | SAB | AUS/NZ | | 8 | 239 | 98 | 2895539 | 85 | 2890342 | 235468 | 100646 | 10 | 32.68 | 2x150bp | |
| BPH3633 | SAB | AUS/NZ | | 30 | 30 | 35 | 2738375 | 30 | 2736582 | 605388 | 169847 | 5 | 32.74 | 2x150bp | |
| BPH3634 | SAB | AUS/NZ | | 8 | 239 | 82 | 2897213 | 74 | 2894214 | 326442 | 104642 | 9 | 32.72 | 2x150bp | |
| BPH3639 | IE | AUS/NZ | | 78 | 88 | 41 | 2745493 | 24 | 2738909 | 638932 | 228957 | 4 | 32.68 | 2x150bp | |
| BPH3645 | SAB | AUS/NZ | | 78 | 78 | 29 | 2690222 | 20 | 2686554 | 634664 | 229064 | 4 | 32.73 | 2x150bp | |
| BPH3690 | SAB | AUS/NZ | | 22 | 22 | 113 | 2788958 | 92 | 2781137 | 169097 | 80856 | 13 | 32.75 | 2x150bp | |
| BPH3695 | SAB | AUS/NZ | | 101 | 101 | 86 | 2740221 | 35 | 2720722 | 963065 | 233315 | 3 | 32.81 | 2x150bp | |
| BPH3716 | SAB | AUS/NZ | | 5 | 5 | 39 | 2803388 | 35 | 2801908 | 327997 | 168349 | 7 | 32.69 | 2x150bp | |
| BPH3719 | SAB | AUS/NZ | | 7 | 7 | 88 | 2801663 | 36 | 2782495 | 466680 | 216047 | 4 | 32.77 | 2x150bp | |
| BPH3727 | SAB | AUS/NZ | | 8 | 368 | 189 | 3061194 | 124 | 3036760 | 156233 | 72462 | 13 | 32.73 | 2x150bp | |
| ST20091659 | IE | FR | | 1 | 188 | 23 | 2731110 | 14 | 2727744 | 1037373 | 292011 | 3 | 32.71 | 2x150bp | |
| ST20091660 | IE | FR | | 15 | 15 | 42 | 2769464 | 24 | 2762583 | 412494 | 244926 | 5 | 32.75 | 2x150bp | |
| ST20091661 | IE | FR | | 15 | 15 | 37 | 2711904 | 28 | 2708750 | 332507 | 215211 | 6 | 32.72 | 2x150bp | |
| ST20091662 | SAB | FR | | 59 | 59 | 44 | 2688726 | 37 | 2686199 | 317224 | 128229 | 7 | 32.72 | 2x150bp | |
| ST20091663 | IE | FR | | 15 | 15 | 31 | 2723187 | 23 | 2720336 | 545769 | 437075 | 3 | 32.69 | 2x150bp | |
| ST20091665 | IE | FR | | 30 | 30 | 102 | 2826386 | 60 | 2810622 | 326876 | 157669 | 7 | 32.72 | 2x150bp | |
| ST20091666 | IE | FR | Mice pool | 5 | 5 | 39 | 2747097 | 36 | 2745968 | 437246 | 208196 | 5 | 32.69 | 2x150bp | |
| ST20091669 | IE | FR | | 8 | 8 | 38 | 2844715 | 29 | 2841449 | 699553 | 544858 | 3 | 32.65 | 2x150bp | |
| ST20091692 | IE | FR | | 8 | 8 | 22 | 2726690 | 15 | 2724043 | 819212 | 806093 | 2 | 32.68 | 2x150bp | |
| ST20091693 | IE | FR | Mice pool | 45 | 54 | 31 | 2705100 | 25 | 2703073 | 655145 | 343041 | 3 | 32.75 | 2x150bp | |
| ST20091694 | IE | FR | | 1 | 1 | 26 | 2744201 | 17 | 2740881 | 518721 | 379296 | 3 | 32.66 | 2x150bp | |
| ST20091729 | IE | FR | | 22 | 22 | 61 | 2694714 | 47 | 2689478 | 251647 | 164133 | 7 | 32.75 | 2x150bp | |
| ST20091949 | IE | FR | Mice pool | 5 | 5 | 34 | 2757611 | 27 | 2754490 | 606326 | 422703 | 3 | 32.78 | 2x250bp | |
| ST20091950 | SAB | FR | | 8 | 8 | 77 | 2739814 | 67 | 2736120 | 289724 | 108235 | 8 | 32.65 | 2x150bp | |
| ST20091954 | SAB | FR | | 30 | 30 | 73 | 2874146 | 61 | 2869676 | 350042 | 85290 | 8 | 32.68 | 2x150bp | |
| ST20091965 | SAB | FR | | 182 | 3803 | 19 | 2701719 | 12 | 2699029 | 1328088 | 1010749 | 2 | 32.67 | 2x150bp | |
| ST20100400 | IE | FR | Mice pool | 45 | 45 | 24 | 2737310 | 18 | 2735201 | 1117137 | 601412 | 2 | 32.79 | 2x150bp | |
| ST20100856 | IE | FR | | 8 | 8 | 33 | 2820499 | 24 | 2817198 | 736873 | 379384 | 3 | 32.67 | 2x150bp | |
| ST20100857 | IE | FR | | 398 | 398 | 30 | 2683075 | 21 | 2679779 | 723973 | 406258 | 3 | 32.82 | 2x150bp | |
| ST20100859 | IE | FR | Mice pool | 5 | 5 | 24 | 2772192 | 22 | 2771314 | 544854 | 428438 | 3 | 32.78 | 2x250bp | |
| ST20100860 | IE | FR | | 30 | 30 | 50 | 2814186 | 42 | 2811286 | 314859 | 175386 | 6 | 32.71 | 2x150bp | |
| ST20100871 | SAB | FR | | 30 | 30 | 50 | 2808419 | 43 | 2805870 | 478475 | 170637 | 6 | 32.71 | 2x150bp | |
| ST20100872 | IE | FR | | 121 | 121 | 43 | 2752579 | 35 | 2749651 | 371539 | 176553 | 6 | 32.71 | 2x150bp | |
| ST20100875 | IE | FR | | 8 | 27 | 17 | 2704384 | 11 | 2702089 | 1342060 | 656382 | 2 | 32.68 | 2x150bp | |
| ST20100878 | IE | FR | | 78 | 78 | 20 | 2759693 | 15 | 2757810 | 1303860 | 623517 | 2 | 32.73 | 2x150bp | |
| ST20100880 | SAB | FR | | 30 | 34 | 53 | 2793745 | 46 | 2790969 | 473432 | 174951 | 5 | 32.72 | 2x150bp | |
| ST20100881 | IE | FR | Mice pool | 45 | 45 | 28 | 2732392 | 19 | 2729149 | 621868 | 487691 | 3 | 32.79 | 2x150bp | |
| ST20100882 | IE | FR | | 1 | 188 | 23 | 2736477 | 13 | 2732768 | 1081446 | 301004 | 2 | 32.69 | 2x150bp | |
| ST20101138 | IE | FR | Mice pool | 45 | 45 | 23 | 2745367 | 18 | 2743530 | 1160604 | 598484 | 2 | 32.80 | 2x150bp | |
| ST20101156 | IE | FR | Mice pool | 45 | 45 | 22 | 2755725 | 18 | 2754247 | 987369 | 625911 | 2 | 32.78 | 2x150bp | |
| ST20101161 | IE | FR | | 1 | 9 | 18 | 2733049 | 13 | 2731228 | 1231295 | 1011826 | 2 | 32.66 | 2x150bp | |
| ST20101186 | IE | FR | | 15 | 15 | 35 | 2754158 | 29 | 2751881 | 366627 | 249952 | 5 | 32.77 | 2x150bp | |
| ST20101417 | SAB | FR | | 12 | 1954 | 18 | 2698642 | 12 | 2696514 | 1297027 | 660796 | 2 | 32.74 | 2x150bp | |
| ST20101418 | IE | FR | | 15 | 10021* | 45 | 2727997 | 25 | 2720584 | 576184 | 411649 | 3 | 32.70 | 2x150bp | *Unknown ST |
| ST20101419 | SAB | FR | | 121 | 121 | 50 | 2831137 | 40 | 2827742 | 466143 | 185032 | 5 | 32.76 | 2x150bp | |
| ST20101420 | IE | FR | Mice pool + Rabbit | 5 | 5 | 28 | 2801217 | 25 | 2800175 | 643319 | 289139 | 3 | 32.68 | 2x150bp | |
| ST20101424 | IE | FR | | 121 | 121 | 39 | 2706980 | 32 | 2704540 | 371263 | 137197 | 6 | 32.69 | 2x150bp | |
| ST20101427 | IE | FR | Mice pool | 2867* | 2867 | 19 | 2747149 | 11 | 2744234 | 1256145 | 703164 | 2 | 32.74 | 2x150bp | assigned as CC5 and used as such in the mice pool |
| ST20101428 | IE | FR | | 12 | 12 | 23 | 2674023 | 17 | 2671981 | 973452 | 630608 | 2 | 32.71 | 2x150bp | |
| ST20101429 | IE | FR | | 398 | 398 | 33 | 2738722 | 26 | 2736181 | 459575 | 307399 | 4 | 32.83 | 2x150bp | |
| ST20101433 | IE | FR | | 97 | 97 | 21 | 2737423 | 17 | 2735889 | 796339 | 672735 | 2 | 32.69 | 2x150bp | |
| ST20101434 | SAB | FR | Mice pool | 45 | 45 | 103 | 2797938 | 35 | 2772729 | 725666 | 389279 | 3 | 32.79 | 2x150bp | |
| ST20101435 | SAB | FR | Mice pool | 5 | 5 | 26 | 2738696 | 21 | 2736809 | 821773 | 367215 | 3 | 32.78 | 2x150bp | |
| ST20101437 | SAB | FR | | 30 | 34 | 40 | 2819877 | 33 | 2817044 | 645522 | 175489 | 4 | 32.71 | 2x150bp | |
| ST20101438 | IE | FR | Mice pool | 45 | 45 | 53 | 2744381 | 23 | 2733371 | 1144247 | 1027145 | 2 | 32.80 | 2x150bp | |
| ST20101445 | IE | FR | | 15 | 15 | 30 | 2732020 | 23 | 2729396 | 413172 | 249956 | 5 | 32.69 | 2x150bp | |
| ST20101531 | IE | FR | | 7 | 7 | 42 | 2821651 | 29 | 2817025 | 707135 | 422963 | 3 | 32.76 | 2x150bp | |
| ST20101696 | SAB | FR | | 30 | 30 | 34 | 2789011 | 28 | 2786754 | 495557 | 223981 | 4 | 32.70 | 2x150bp | |
| ST20101705 | IE | FR | | 97 | 464 | 22 | 2719612 | 17 | 2717681 | 853436 | 711047 | 2 | 32.70 | 2x150bp | |
| ST20101718 | IE | FR | | 15 | 15 | 29 | 2740919 | 21 | 2738194 | 437240 | 314928 | 4 | 32.78 | 2x150bp | |
| ST20101737 | SAB | FR | Mice pool | 5 | 5 | 28 | 2738860 | 21 | 2736067 | 660488 | 388986 | 3 | 32.75 | 2x150bp | |
| ST20101785 | SAB | FR | Mice pool | 45 | 45 | 28 | 2762786 | 21 | 2760319 | 1282211 | 393569 | 2 | 32.83 | 2x150bp | |
| ST20101786 | SAB | FR | Mice pool | 45 | 45 | 32 | 2732782 | 23 | 2729550 | 533242 | 423025 | 3 | 32.82 | 2x150bp | |
| ST20101789 | IE | FR | Mice pool + Rabbit | 5 | 10022* | 29 | 2703636 | 23 | 2701494 | 810426 | 681648 | 2 | 32.74 | 2x150bp | *Unknown ST |
| ST20101791 | SAB | FR | Mice pool + Rabbit | 5 | 5 | 775 | 3038806 | 312 | 2835251 | 94407 | 16345 | 47 | 32.69 | 2x250bp | |
| ST20101794 | SAB | FR | Mice pool | 5 | 5 | 36 | 2752843 | 33 | 2751657 | 340255 | 208199 | 5 | 32.68 | 2x150bp | |
| ST20101796 | SAB | FR | | 22 | 22 | 48 | 2722555 | 38 | 2718688 | 326530 | 160461 | 7 | 32.72 | 2x150bp | |
| ST20101934 | IE | FR | | 8 | 8 | 31 | 2787237 | 23 | 2784238 | 736897 | 488553 | 3 | 32.62 | 2x150bp | |
| ST20102127 | SAB | FR | | 12 | 12 | 24 | 2719084 | 16 | 2716205 | 974813 | 660571 | 2 | 32.70 | 2x150bp | |
| ST20102139 | IE | FR | | 398 | 398 | 39 | 2732600 | 33 | 2730602 | 423468 | 237713 | 5 | 32.81 | 2x150bp | |
| ST20102292 | IE | FR | Mice pool | 5 | 5 | 26 | 2744331 | 21 | 2742213 | 1021502 | 841289 | 2 | 32.82 | 2x250bp | |
| ST20102293 | IE | FR | | 1 | 1 | 18 | 2775733 | 11 | 2773123 | 1105777 | 1057800 | 2 | 32.70 | 2x150bp | |
| ST20102295 | IE | FR | Mice pool + Rabbit | 5 | 5 | 32 | 2719305 | 24 | 2716013 | 973843 | 646985 | 2 | 32.75 | 2x250bp | |

| | | | | | | | | | | | | | | |
|---|---|---|---|---|---|---|---|---|---|---|---|---|---|---|
| ST20102297 | SAB | FR | | 22 | 22 | 73 | 2690006 | 65 | 2687170 | 208980 | 74110 | 12 | 32.70 | 2x150bp |
| ST20102298 | SAB | FR | | 8 | 8 | 30 | 2842349 | 21 | 2838999 | 864751 | 699093 | 2 | 32.63 | 2x150bp |
| ST20102300 | SAB | FR | | 97 | 97 | 22 | 2730445 | 19 | 2729281 | 792622 | 287304 | 3 | 32.73 | 2x150bp |
| ST20102301 | SAB | FR | | 12 | 12 | 18 | 2696949 | 12 | 2694922 | 1296308 | 660892 | 2 | 32.74 | 2x150bp |
| ST20102302 | IE | FR | Mice pool | 45 | 1165 | 25 | 2747676 | 21 | 2746216 | 1097886 | 597881 | 2 | 32.79 | 2x150bp |
| ST20102303 | SAB | FR | | 59 | 59 | 43 | 2748900 | 35 | 2745956 | 331871 | 149185 | 6 | 32.75 | 2x150bp |
| ST20102304 | SAB | FR | | 121 | 3213 | 74 | 2769389 | 66 | 2766526 | 451743 | 86710 | 8 | 32.76 | 2x150bp |
| ST20102307 | IE | FR | | 398 | 398 | 24 | 2690337 | 17 | 2687833 | 723767 | 338943 | 3 | 32.79 | 2x150bp |
| ST20102308 | SAB | FR | | 1 | 1 | 18 | 2780742 | 11 | 2778018 | 1266919 | 1029549 | 2 | 32.69 | 2x150bp |
| ST20110014 | IE | FR | Mice pool | 45 | 45 | 25 | 2705814 | 20 | 2704020 | 1117181 | 648572 | 2 | 32.77 | 2x150bp |
| ST20110196 | IE | FR | | 8 | 72 | 29 | 2709015 | 21 | 2706070 | 760576 | 709847 | 2 | 32.71 | 2x150bp |
| ST20110285 | IE | FR | Mice pool | 5 | 146 | 23 | 2704407 | 17 | 2702233 | 590218 | 324845 | 3 | 32.72 | 2x150bp |
| ST20110286 | IE | FR | | 101 | 101 | 17 | 2699055 | 11 | 2696981 | 1347382 | 632012 | 2 | 32.74 | 2x150bp |
| ST20110552 | SAB | FR | | 15 | 15 | 86 | 2716316 | 78 | 2713526 | 251173 | 71812 | 12 | 32.71 | 2x150bp |
| ST20110553 | SAB | FR | | 398 | 398 | 32 | 2736537 | 24 | 2733554 | 536962 | 410510 | 3 | 32.82 | 2x150bp |
| ST20110554 | IE | FR | | 7 | 7 | 27 | 2795613 | 23 | 2794253 | 422392 | 282858 | 5 | 32.83 | 2x150bp |
| ST20110555 | IE | FR | | 12 | 4302 | 18 | 2695997 | 12 | 2693869 | 1295814 | 660832 | 2 | 32.74 | 2x150bp |
| ST20110557 | SAB | FR | | 97 | 97 | 21 | 2715399 | 17 | 2713955 | 706561 | 671374 | 2 | 32.72 | 2x150bp |
| ST20110558 | SAB | FR | | 8 | 8 | 33 | 2891348 | 26 | 2888846 | 813003 | 780146 | 2 | 32.69 | 2x150bp |
| ST20110559 | SAB | FR | | 30 | 30 | 68 | 2851063 | 58 | 2847387 | 421628 | 150428 | 6 | 32.74 | 2x150bp |
| ST20110560 | IE | FR | Mice pool + Rabbit | 5 | 5 | 54 | 2811481 | 45 | 2807511 | 1020972 | 408518 | 2 | 32.79 | 2x250bp |
| ST20110561 | SAB | FR | | 97 | 97 | 34 | 2723571 | 26 | 2720632 | 556254 | 204005 | 4 | 32.71 | 2x150bp |
| ST20110562 | SAB | FR | Mice pool | 5 | 5 | 52 | 2809865 | 44 | 2806585 | 483329 | 196056 | 4 | 32.77 | 2x250bp |
| ST20110890 | IE | FR | | 8 | 646 | 24 | 2760367 | 17 | 2757729 | 1217134 | 685009 | 2 | 32.65 | 2x150bp |
| ST20111082 | SAB | FR | | 398 | 398 | 51 | 2773251 | 33 | 2766254 | 565121 | 200985 | 5 | 32.83 | 2x150bp |
| ST20111220 | IE | FR | | 121 | 121 | 46 | 2782352 | 38 | 2779526 | 402453 | 143153 | 7 | 32.75 | 2x150bp |
| ST20111347 | IE | FR | | 15 | 15 | 29 | 2764305 | 21 | 2761370 | 571777 | 364751 | 3 | 32.74 | 2x150bp |
| ST20111348 | IE | FR | | 30 | 30 | 69 | 2847264 | 57 | 2842494 | 369131 | 124447 | 7 | 32.74 | 2x150bp |
| ST20111367 | IE | FR | Mice pool | 45 | 46 | 33 | 2748889 | 19 | 2743749 | 985713 | 543299 | 2 | 32.81 | 2x150bp |
| ST20111368 | IE | FR | | 15 | 15 | 135 | 2662211 | 121 | 2657106 | 159667 | 46125 | 19 | 32.74 | 2x150bp |
| ST20111370 | SAB | FR | Mice pool | 5 | 5 | 24 | 2697982 | 21 | 2696827 | 761983 | 329645 | 3 | 32.74 | 2x150bp |
| ST20111371 | SAB | FR | Mice pool | 45 | 45 | 56 | 2748603 | 28 | 2738284 | 699420 | 444664 | 3 | 32.82 | 2x150bp |
| ST20111372 | SAB | FR | Mice pool + Rabbit | 5 | 5 | 182 | 2854068 | 88 | 2816283 | 241389 | 77621 | 13 | 32.70 | 2x250bp |
| ST20111441 | IE | FR | | 1 | 188 | 20 | 2736687 | 12 | 2733678 | 1017971 | 593142 | 2 | 32.66 | 2x150bp |
| ST20111452 | IE | FR | Mice pool | 5 | 146 | 38 | 2720296 | 30 | 2716915 | 534687 | 358572 | 3 | 32.75 | 2x250bp |
| ST20111461 | SAB | FR | | 80 | 80 | 26 | 2780447 | 20 | 2778300 | 542091 | 248802 | 4 | 32.77 | 2x150bp |
| ST20111792 | IE | FR | Mice pool | 5 | 3014 | 45 | 2768555 | 39 | 2766408 | 583843 | 200793 | 4 | 32.71 | 2x150bp |
| ST20112201 | SAB | FR | | 25 | 25 | 16 | 2719351 | 14 | 2718583 | 1353516 | 770842 | 2 | 32.67 | 2x150bp |
| ST20112206 | SAB | FR | | 30 | 30 | 131 | 2779884 | 63 | 2754806 | 478158 | 167601 | 4 | 32.70 | 2x150bp |
| ST20112208 | SAB | FR | Mice pool | 5 | 5 | 29 | 2819845 | 24 | 2818037 | 506404 | 266951 | 4 | 32.73 | 2x150bp |
| ST20112212 | IE | FR | | 30 | 30 | 456 | 3026907 | 318 | 2968288 | 273338 | 50964 | 16 | 32.76 | 2x150bp |
| ST20112213 | IE | FR | | 30 | 30 | 120 | 2862041 | 71 | 2844058 | 192805 | 97010 | 10 | 32.74 | 2x150bp |
| ST20112214 | IE | FR | | 398 | 398 | 30 | 2722354 | 22 | 2719401 | 560557 | 410546 | 3 | 32.86 | 2x150bp |
| ST20112215 | IE | FR | Mice pool | 45 | 45 | 26 | 2772997 | 19 | 2770536 | 594636 | 519923 | 3 | 32.79 | 2x150bp |
| ST20112216 | SAB | FR | | 121 | 121 | 45 | 2781315 | 38 | 2778666 | 418301 | 162523 | 5 | 32.74 | 2x150bp |
| ST20112222 | SAB | FR | Mice pool | 45 | 45 | 25 | 2747903 | 20 | 2746070 | 743556 | 426821 | 3 | 32.80 | 2x150bp |
| ST20112223 | SAB | FR | | 152 | 152 | 31 | 2719590 | 24 | 2716983 | 446264 | 289433 | 4 | 32.69 | 2x150bp |
| ST20112455 | IE | FR | Mice pool | 5 | 5 | 34 | 2747673 | 26 | 2744555 | 457508 | 238405 | 4 | 32.78 | 2x150bp |
| ST20120191 | IE | FR | | 15 | 15 | 34 | 2781982 | 24 | 2778419 | 569291 | 315196 | 3 | 32.71 | 2x150bp |
| ST20120194 | IE | FR | | 8 | 8 | 36 | 2833399 | 28 | 2830345 | 463839 | 307614 | 4 | 32.72 | 2x150bp |
| ST20120196 | IE | FR | | 15 | 15 | 30 | 2685077 | 23 | 2682486 | 570484 | 249954 | 4 | 32.69 | 2x150bp |
| ST20120197 | SAB | FR | | 15 | 15 | 89 | 2740119 | 35 | 2720369 | 611503 | 271497 | 4 | 32.72 | 2x150bp |
| ST20120198 | SAB | FR | | 8 | 8 | 13 | 2798262 | 9 | 2796664 | 1373470 | 1030731 | 2 | 32.60 | 2x150bp |
| ST20120202 | IE | FR | | 398 | 398 | 29 | 2695465 | 20 | 2692211 | 533066 | 261797 | 4 | 32.81 | 2x150bp |
| ST20120204 | SAB | FR | Mice pool | 5 | 5 | 24 | 2752621 | 21 | 2751338 | 589625 | 335222 | 3 | 32.74 | 2x250bp |
| ST20120205 | SAB | FR | | 30 | 30 | 39 | 2844828 | 35 | 2843450 | 523903 | 196306 | 5 | 32.74 | 2x150bp |
| ST20120206 | SAB | FR | Mice pool + Rabbit | 5 | 5 | 27 | 2744744 | 23 | 2742972 | 856426 | 766401 | 2 | 32.71 | 2x250bp |
| ST20120207 | SAB | FR | Mice pool | 5 | 5 | 23 | 2726591 | 17 | 2724301 | 1018704 | 702359 | 2 | 32.77 | 2x150bp |
| ST20120208 | SAB | FR | | 30 | 4796 | 99 | 2832435 | 45 | 2813181 | 392372 | 175391 | 6 | 32.72 | 2x150bp |
| ST20120209 | SAB | FR | | 8 | 72 | 27 | 2720648 | 22 | 2718810 | 760524 | 671631 | 2 | 32.68 | 2x150bp |
| ST20120210 | SAB | FR | | 121 | 121 | 50 | 2798984 | 42 | 2796220 | 371214 | 137135 | 6 | 32.73 | 2x150bp |
| ST20120211 | SAB | FR | Mice pool + Rabbit | 5 | 5 | 33 | 2772972 | 27 | 2770558 | 1021794 | 766627 | 2 | 32.78 | 2x250bp |
| ST20120234 | IE | FR | | 8 | 8 | 29 | 2828746 | 23 | 2826526 | 652091 | 326179 | 3 | 32.70 | 2x150bp |
| ST20120263 | SAB | FR | Mice pool | 45 | 45 | 37 | 2744409 | 26 | 2740004 | 682303 | 347566 | 3 | 32.78 | 2x150bp |
| ST20120276 | SAB | FR | Mice pool | 45 | 45 | 28 | 2725655 | 20 | 2722671 | 986983 | 464019 | 2 | 32.79 | 2x150bp |
| ST20120283 | SAB | FR | Mice pool | 45 | 45 | 27 | 2716282 | 22 | 2714494 | 672320 | 421044 | 3 | 32.79 | 2x150bp |
| ST20120303 | SAB | FR | Mice pool | 45 | 45 | 29 | 2761656 | 21 | 2758628 | 781928 | 342248 | 3 | 32.77 | 2x150bp |
